# BOCS: DNA k-mer content and scoring for rapid genetic biomarker identification at low coverage

**DOI:** 10.1101/631333

**Authors:** Lee E. Korshoj, Prashant Nagpal

**Affiliations:** Department of Chemical and Biological Engineering, University of Colorado Boulder, Boulder, CO 80303, USA; Renewable and Sustainable Energy Institute (RASEI), University of Colorado Boulder, Boulder, CO 80303, USA; Materials Science and Engineering, University of Colorado Boulder, Boulder, CO 80303, USA

**Keywords:** DNA sequencing, optical sequencing, high-throughput diagnostics, biomarker detection, multidrug-resistant bacteria, Raman spectroscopy

## Abstract

A single, inexpensive diagnostic test capable of rapidly identifying a wide range of genetic biomarkers would prove invaluable in precision medicine. Previous work has demonstrated the potential for high-throughput, label-free detection of A-G-C-T content in DNA k-mers, providing an alternative to single-letter sequencing while also having inherent lossy data compression and massively parallel data acquisition. Here, we apply a new bioinformatics algorithm – block optical content scoring (BOCS) – capable of using the high-throughput content k-mers for rapid, broad-spectrum identification of genetic biomarkers. BOCS uses content-based sequence alignment for probabilistic mapping of k-mer contents to gene sequences within a biomarker database, resulting in a probability ranking of genes on a content score. Simulations of the BOCS algorithm reveal high accuracy for identification of single antibiotic resistance genes, even in the presence of significant sequencing errors (100% accuracy for no sequencing errors, and > 90% accuracy for sequencing errors at 20%), and at well below full coverage of the genes. Simulations for detecting multiple resistance genes within a methicillin-resistant *Staphylococcus aureus* (MRSA) strain showed 100% accuracy at an average gene coverage of merely 0.515, when the k-mer lengths were variable and with 4% sequencing error within the k-mer blocks. Extension of BOCS to cancer and other genetic diseases met or exceeded the results for resistance genes. Combined with a high-throughput content-based sequencing technique, the BOCS algorithm potentiates a test capable of rapid diagnosis and profiling of genetic biomarkers ranging from antibiotic resistance to cancer and other genetic diseases.

## 1. Introduction

In the push for precision medicine, there is an increasing demand for inexpensive, non-specific assays capable of broad-spectrum diagnostics, where a single test can rapidly screen an array of biomarkers[1,2]. One immediate application of such a technology is to address the growing threat of antibiotic resistance, a public health crisis that affects nearly two million people in the U.S. annually[3,4]. Rapid, affordable identification of drug-resistance in clinically relevant microbial strains is vital for prescribing patients with appropriate treatment plans to reduce mortality rates and the development of further resistances[5]. Current resistance diagnostics and profiling assays are often performed only after initial antibiotics fail. Most of these assays rely on cell culturing, PCR amplification, and microarray analyses[6–13]. Not only do these tests require hours to days and significant costs, but they are specific for detecting resistances of one or a few well-characterized strains. Next-generation, whole-genome sequencing approaches to resistance screening have shown promise; however, applications of this technology to diagnostics has been limited by lack of standardization protocols and the need for data interpretation leading to long diagnosis times[14–17].

A rapid, broad-spectrum diagnostic technique would also prove invaluable in the screening of cancers and other genetic diseases. Point-of-care diagnostic devices for sensitive and specific detection of cancer biomarkers has long been a goal of the biosensing community[18]. Moreover, scientists and clinicians have long struggled to identify rare, novel, and undiagnosed disorders as evident by initiatives such as the National Institutes of Health (NIH) Undiagnosed Diseases Network[19,20]. For cancers and other genetic diseases, early detection is crucial for patient survival. Current and emerging diagnostics continue to rely on the identification of the protein, peptide, or gene expression biomarkers[21–23]. These diagnostic devices apply an array of nanoelectronic and optical techniques, but like antibiotic resistance assays, are specific for detecting merely one or a few biomarkers for which the device is constructed.

In this study, we present a robust algorithmic platform called block optical content scoring (BOCS) that facilitates rapid, broad-spectrum genetic biomarker identification from DNA k-mer content. This algorithm builds upon previous work demonstrating the use of Raman spectroscopy measurements for high-throughput, label-free detection of A-G-C-T content in DNA k-mers, called block optical sequencing (BOS)[24]. This BOS method is an alternative to single-letter sequencing and has the potential to simultaneously measure DNA k-mer content from millions of fragments simultaneously, thereby converting it into useful genetic information. This approach is akin to sharing and streaming of large multimedia files across the world wide web using a combination of lossless and lossy data compression techniques. Our bioinformatics approach, BOCS, uses the DNA k-mer content for identification of genetic biomarkers through probabilistic mapping of the k-mer content to gene databases. Comprehensive simulations show accurate and specific recognition of antibiotic resistance genes, as well as cancer and other genetic disease genes with less than full coverage of the genes and in the presence of sequencing error. The results shown here for the BOCS algorithm pave the way for a single, inexpensive diagnostic test capable of rapidly identifying a wide range of genetic biomarkers.

## 2. Results

### 2.1 The BOCS algorithm

Given the capability of high-throughput single-molecule Raman spectroscopy measurements in determining DNA k-mer content, the need arises for a way to correlate these content measurements into meaningful genetic information. The potential for coupling a high-throughput measurement system with a broad-spectrum genetic biomarker identification method could lead to a diagnostic platform for rapid point-of-care genetic profiling. Direct applications range from providing clinicians with the information they need to effectively treat multidrug-resistant (MDR) bacterial infections to early detection of cancers and other genetic diseases that previously had no screening techniques. Therefore, we introduce the BOCS algorithm, which uses DNA k-mer content for broad-spectrum genetic biomarker recognition.

In designing BOCS (schematic in Figure 1), we took inspiration from probability-based sequence analyzers such as those employed for protein identification from mass spectrometry data[25–27], as well as alignment programs used to map next-generation sequencing reads to reference genomes[28,29]. In a similar nature to these methods, the BOCS algorithm relies on probabilistic content alignments to reference sequences for genetic biomarkers. The BOCS algorithm requires 1) the log of all k-mer blocks and their content and 2) a database containing gene sequences for the genetic biomarkers being investigated (e.g., antibiotic resistance, cancer, or other genetic diseases). The algorithm cycles through each k-mer block and performs a content-based alignment with each gene sequence in the database, translating through the gene sequence one nucleotide at a time and tracking the number of match locations – where the k-mer block content matches the content of the k-length gene sequence. A probability is calculated for each gene after each block is aligned with it. This raw probability (*P*_*R*_) is simply the number of observed matches divided by the calculated number of matches that are statistically expected to occur randomly. It is based on the fundamental idea that genes in the database that are most similar to the k-mer blocks in terms of their content should have the most matches during alignment, and therefore deviate the most significantly from the random case. The raw probability is calculated from the number of match locations (*m*), the length of the k-mer block (*k*) and its content in terms of the number of A-G-C-T nucleotides, and length of the gene (*g*_*L*_), shown below for an arbitrary gene (*x*):

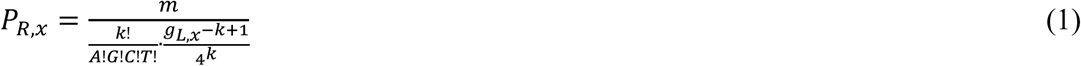

**Fig. 1.**
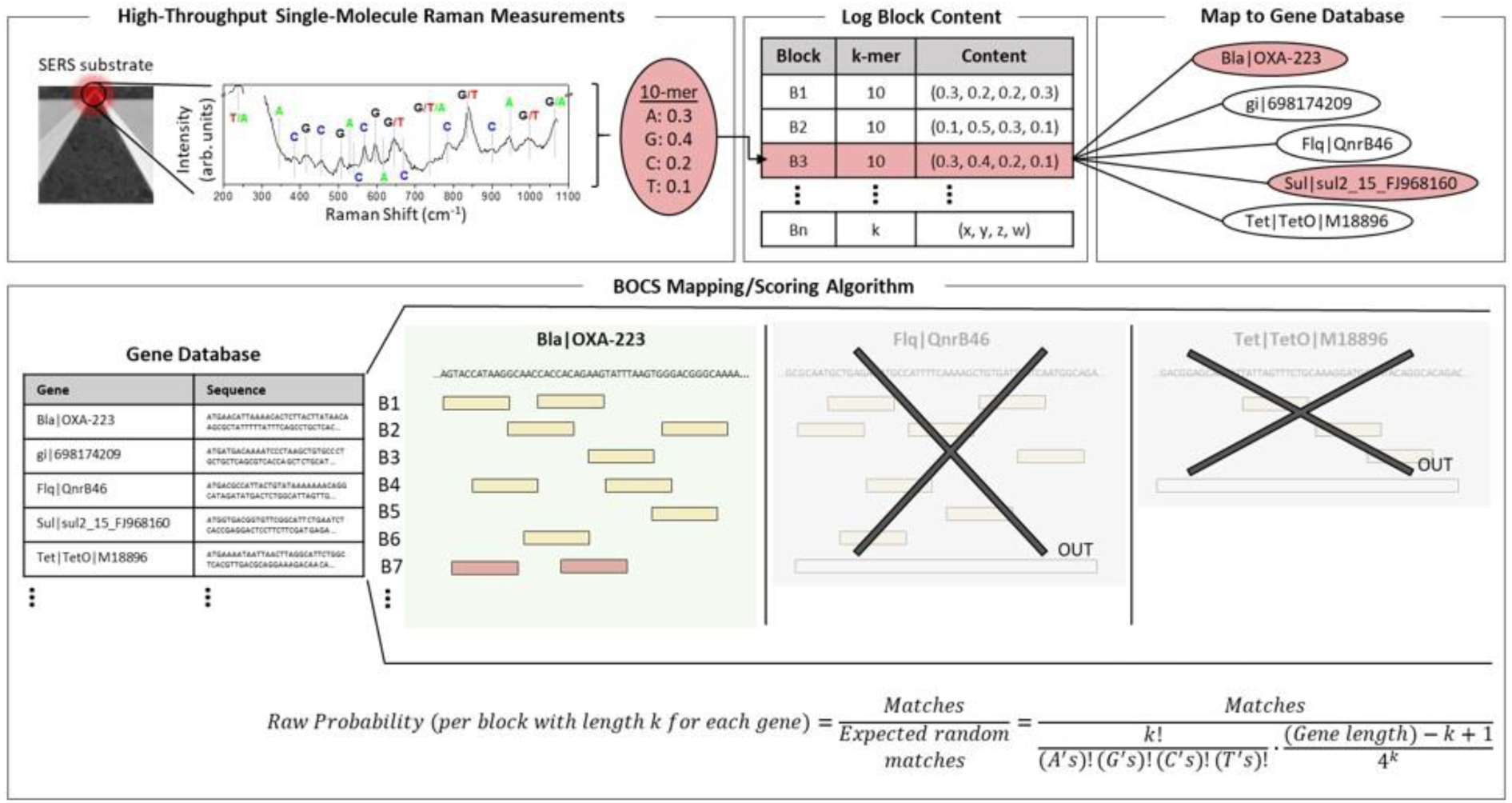
Schematic for the BOCS algorithm. The BOCS algorithm takes measurements of DNA k-mer content from high-throughput Raman spectroscopy measurements and maps them to gene databases for probabilistic determination of genetic biomarkers at low coverages. Starting with a log of measured k-mer content blocks (Bl…Bn as shown) and a genetic biomarker database (excerpts from the MEGARes antibiotic resistance database are shown), the blocks are individually aligned to each gene in the database based on content. This alignment consists of finding all match locations for the k-mer block content within a gene via translating through the gene one nucleotide at a time and looking at fragments of length k. For each block, a raw probability can be calculated for each gene based on the number of matches for the k-mer block content within the gene, length of the k-mer block, and length of the gene (calculation shown in the schematic). As more blocks are analyzed, probabilities are compounded and genes in the database are ranked. The gene(s) from which the Raman-analyzed k-mer blocks originate quickly generate the top probabilities and can often be determined in coverages ≪ 1.0, meaning that only a small fraction of the gene blocks need to be analyzed for identification of a specific genetic biomarker.

In the case where no matches are found for a gene, the gene is given a penalty score in place of the raw probability (adjustable parameter for the algorithm, normally in the range of 0.01-0.10). After the analysis of a block (i.e., when the block has been content aligned to each gene in the database), this raw probability is normalized by the maximum raw probability observed for all genes (*P*_*R*_ becomes 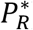). While this raw probability itself is not the score on which biomarker identifications are made, it is the basis for many of the six probability factors that make up the overall content score.

After the content alignment of a block has been completed for all genes, and the raw probabilities are calculated for each gene, six probability factors (*PF*) that make up the content score (*CS*) are calculated for each gene. These *PF* values are designed as pattern recognition elements for a customized machine learning enhancement to the algorithm. They were designed to account for repeated trends observed throughout comprehensive analyses of match patterns during content alignment. The first probability factor (*PF*_1_) is the cumulative percent difference from average of the normalized raw probability (*PDiff*) multiplied by the normalized cumulative raw probability, shown below for an arbitrary gene (*x*) after an arbitrary block (*b*_*n*_) in terms of normalized raw probabilities:

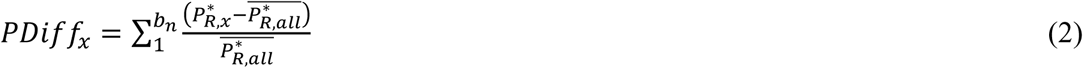

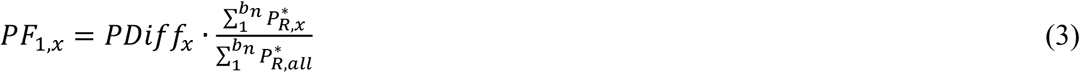

The second probability factor (*PF*_2_) is the total number of blocks, up to the current block, having at least one match from the content alignment:

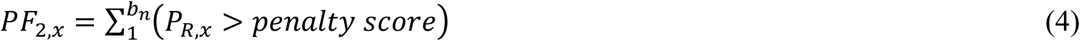

The third probability factor (*PF*_3_) is the product of all normalized raw probabilities taken as the log base 2 sum. Since this leads to negative values, they are flipped by subtracting from the most negative value:

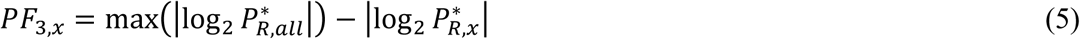

The fourth probability factor (*PF*_4_) is an exponential of the gene coverage (*g*_*cov*_), indicating the fractional number of nucleotides within the gene that have been matched during content alignment:

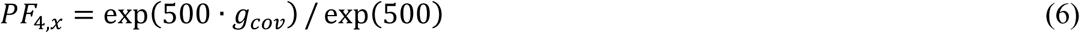

The fifth probability factor (*PF*_5_) is the cumulative slope (*S*_*PF*5_) calculated from the percent difference from average of the normalized raw probability (*PDiff*, equation 2). The slope is calculated for the current block and the nine previous blocks; therefore, this factor does not take effect until the tenth block:

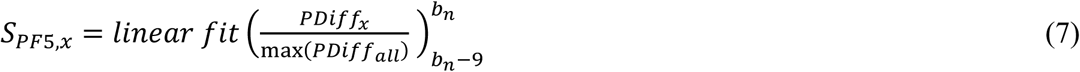

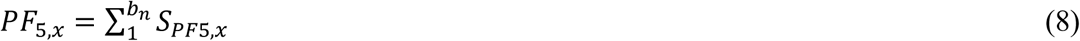

The sixth probability factor (*PF*_6_) is the cumulative difference from average of the normalized raw probability:

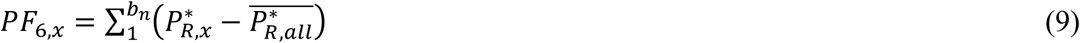

Each of the six *PF* values are normalized individually by the maximum *PF* observed for all genes (*PF* becomes *PF*^*^). This normalization by the maximum ensures equal weighting for the factors when they are added together to give the *CS*:

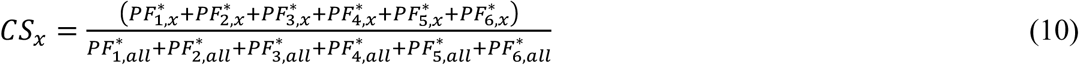

Notice that the *CS* is also normalized; however, here it is by the sum of *CS* values for all of the genes instead of the maximum as for the *PFs*. As each block is analyzed, the *CS* for each gene accumulates, leading to a probabilistic ranking of genes in the database. As demonstrated in the results, the compounded probabilistic content scoring is robust, and can often correlate the k-mer block contents to a positive genetic biomarker identification well below full coverage of the gene.

### 2.2 BOCS for detection of antibiotic resistance

The BOCS algorithm was built into a simulation for large-scale analyses (code available at https://github.com/lkorshoj/Block-Optical-Content-Scoring). The simulation takes gene sequences from a biomarker database and creates k-mer blocks of A-G-C-T content to simulate BOS reads. These simulated BOS reads are then run through the BOCS algorithm against the biomarker database. The goal of the simulation is to see how well the BOCS algorithm can identify the correct gene (out of all others in the database) using merely randomized k-mer blocks of A-G-C-T content. A specific gene from the database can be pulled or a random gene can be selected. The k-mer block lengths, gene coverage, and the number of errors within the blocks can all be set.

For comprehensive testing of the BOCS algorithm, we used the MEGARes database of antimicrobial resistance, composed of 3824 total resistance gene sequences[30]. Due to the phylogeny of annotated genes in MEGARes and other gene databases, the BOCS analysis uses three levels for gene detection. In the order of most broad to most specific they include – class, sub-class, and specific gene. For example, a gene leading to resistance of tetracycline antibiotics could have a class: tetracycline ribosomal protection proteins, sub-class: TETO, and specific gene: TETO-x,y,z (where x, y, z are specific mutations of TETO). Note that we deviate from the MEGARes three-level annotation system for more wide-range applicability with other genetic databases (as demonstrated later). For our BOCS benchmarking analyses, we randomly selected 70 genes having unique sub-classes from the MEGARes database (see the Supplementary Table S1 for details of the genes) and ran 25 repeat simulations on each, where each simulation repeat represents different split locations for the k-mer blocks and a different randomized order in which the blocks are analyzed. In this first set of 1750 simulations, the k-mer blocks were set at k=10, single gene coverage, and no block errors (results are shown in Figure 2 – red). In analyzing the simulation results, we are interested in four main metrics: accuracy, coverage at which a gene is identified, false positives, and specificity. The accuracy is a measure of how often the selected gene, which has been fragmented into randomized k-mers of A-G-C-T content, can be identified. The coverage at which a gene is identified indicates how many blocks less than the total (all blocks correspond to a coverage = 1.0) are needed, eluding to the rapid, robust nature of the algorithm. False positives are a measure of the sensitivity in detection (more false positives means less sensitive). The specificity shows how significantly the gene database can be narrowed as consecutive blocks are analyzed. All of these factors depend on when an identification is made, which is determined as the point where a gene within the database adopts the highest content score and remains there and/or separates itself probabilistically from the rest. False positives arise when genes other than the selected gene meet this identification criterion. Genes within the database can be eliminated when a block shows no content matches during the alignment (this elimination scheme can only be used when there is single coverage for the genes and no block errors). In this first simulation with 70 resistance genes, 100% accuracy (with no false positives) was achieved while requiring an average coverage of merely 0.271 ±0.064 (Figure 2A – red). Additionally, roughly 90% of the genes in the MEGARes database were eliminated by 0.20 coverage (Figure 2B – red). Results for four individual genes within the set of 70 are shown in Figure 2C-E. Although variation in the coverage for identification and specificity are observed, both metrics remain highly favorable (identifications made and the majority of genes in the database eliminated at coverages ≪ 1.0, Figure 2C,D – red). Figure 2E – red shows the rapid separation, and hence identification, of genes from the content scoring. In the case where content scoring separation does not appear as significant (such as for the TEM class A beta-lactamase), this is because all of the top-ranking genes (red line and gray lines in close proximity) are of the same TEM sub-class. Full results for this simulation can be found in Supplementary Table S2.

**Fig. 2.**
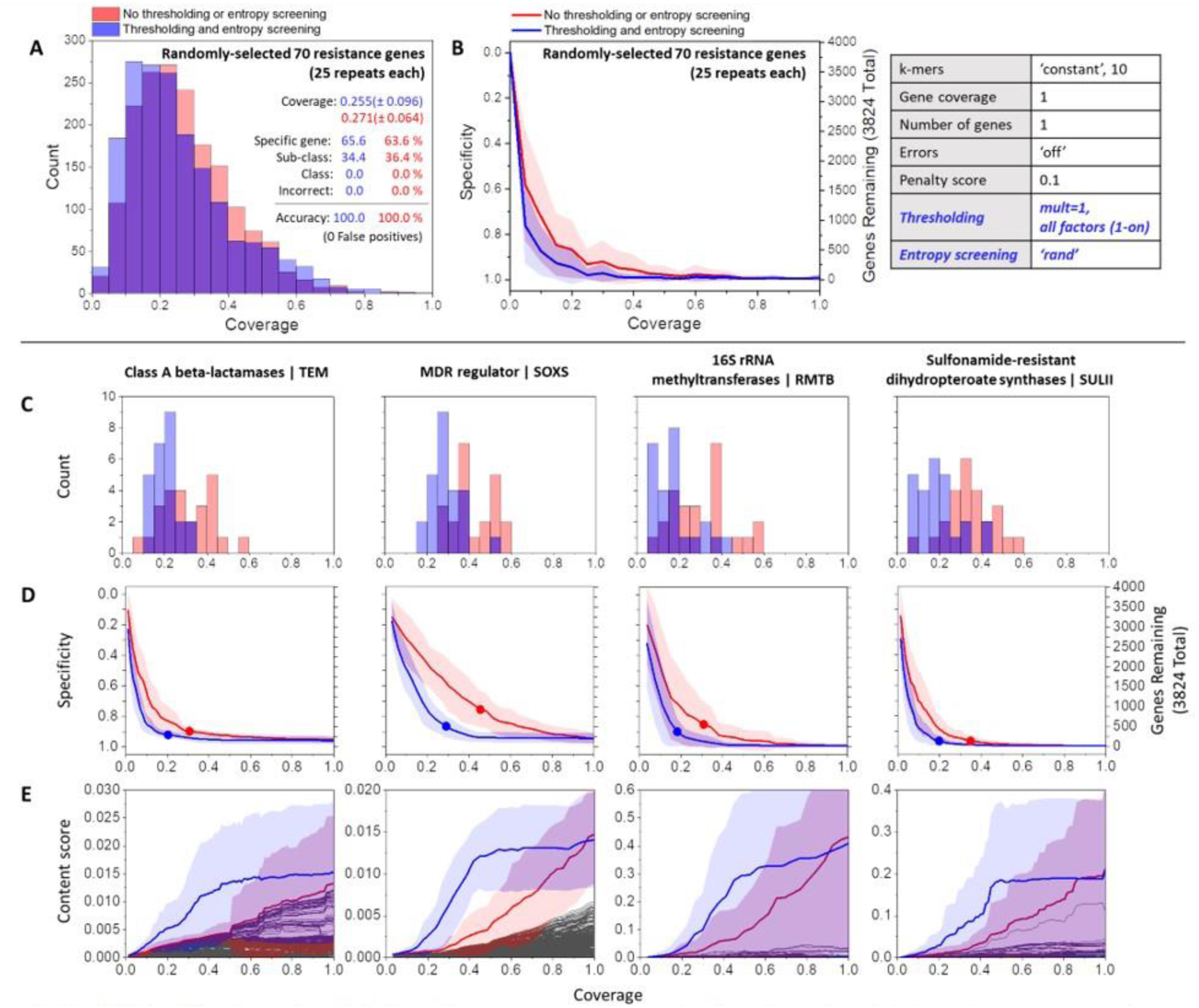
Rapid identification of antibiotic resistance genes. 70 randomly-selected antibiotic resistance genes from the MEGARes database were each run through the BOCS simulation with 25 repeats, for a total of 1750 simulations. Results are shown for both the cases of no thresholding and entropy screening (red) and with thresholding and entropy screening (blue). **(A**) Histogram of the coverage at which a resistance gene is identified combined for all 70-gene simulations. Details of the average coverage and accuracy are given in the inset. Results demonstrate that most genes can be identified with 100% accuracy at merely 0.15–0.30 coverage of the gene. With thresholding and entropy screening, the average coverage decreased and led to more specific gene identifications. **(B**) Specificity with increasing coverage for all compiled 70-gene simulations. It is demonstrated that about 90% of the genes in the database can be eliminated at coverages as low as 0.10. With thresholding and entropy screening, more genes are eliminated at lower coverages leading to higher specificity in the identification process. **(C)** Histograms of the average coverage at which a resistance gene is identified for four individual genes (gene labels shown at the top). The histograms show a clear shift towards lower coverages for the thresholding and entropy screening case (data from the 25 simulations for each case are shown). **(D)** Specificity with increasing coverage for the four individual genes. Dots indicate the average locations at which a gene is identified. Again, significant shifts towards lower coverages are seen in the case with thresholding and entropy screening. **(E)** Increasing content scores with coverage for four individual genes. The selected genes are colored blue/red for cases with/without thresholding and entropy screening. The grayed lines are all other genes in the resistance gene database (3823 of the 3824 total). As coverage increases (i.e., as more blocks are analyzed), the selected genes quickly separate themselves from the others probabilistically, leading to their identification at low coverages. The separation happens sooner, and more significantly, in the case of thresholding and entropy screening.

When looking at the content scoring for this first set of simulations on antibiotic resistance genes, we observed the most significant spikes in probabilities when the number of permutations for a particular block content was low (i.e., the value *k*!/(*A*! *G*! *C*! *T*!) was low). This led to the idea of preferably analyzing these ‘low entropy’ blocks before others in a process we call entropy screening. In the simulation, entropy screening can be applied in a random fashion (in the random order to which the blocks are scattered) or an ideal fashion (in order of low entropy to high entropy). Moreover, we noticed that in the majority of simulations, genes within the database that had probabilistically become irrelevant were still being analyzed as potential candidates. To alleviate this, we implemented a thresholding system to remove genes with lowest probability ranks after each round of block analyses. This type of thresholding based on content score ranking is also necessary to eliminate genes for the cases when there are more than a single gene or gene coverage as well as sequencing errors, where eliminations based on no content matches to a block would lead to significant identification error and decrease the overall accuracy. In the simulation, thresholding can be implemented based on the rank of the content score, as well as each of the individual probability factors, and each can be multiplied by a factor to increase/decrease the sensitivity of thresholding. With the thresholding and entropy screening in place, the first simulation with 70 resistance genes was re-run (again with k-mer blocks set at k=10, single gene coverage, and no block errors, with 25 repeat simulations per gene). Looking at the results shown in Figure 2A,B – blue, we again see 100% accuracy (with no false positives), this time achieved at an average coverage of only 0.255 ±0.096, and roughly 90% of the genes in the database were eliminated by 0.10 coverage. For four individual gene examples (Figure 2C-E) significant improvements in BOCS metrics were seen for the case of thresholding and entropy screening. Not only did we achieve significant shifts towards lower coverage (Figure 2C – blue) and higher specificity (Figure 2D – blue), but we see faster, more prominent increases in the content scores for the genes we are attempting to identify (Figure 2E – blue). Full results for this simulation can be found in Supplementary Table S3. This first round of simulations clearly demonstrated the rapidness to which BOCS can identify genes based merely on randomized k-mer content blocks, and improvements can be further seen with thresholding and entropy screening.

### 2.3 BOCS with sequencing variability

We next sought to test the limits of the BOCS algorithm by introducing sequencing variability in the form of fluctuating k-mer block lengths, block errors, and using blocks from multiple genes. All of these settings can be input on the BOCS simulation, and each of the simulations were run with the thresholding (using all probability factors and content score) and random entropy screening. First looking at k-mer lengths, we ran two sets of simulations with constant k-mer lengths different from the k=10 case used previously – one with k=8 and another with k=12. Then another set of simulations were run for varying k-mer lengths centered around k=10. For this, k-mer lengths for each block are randomly picked from a normal distribution centered around k=10, leading to a distribution of k-mer lengths in the range k=6-14. For each of these simulations, the same 70 MEGARes genes were used, again with 25 repeats. Results in Figure 3A-C show that accuracy, coverage for identification, and the false positive rate are weakly correlated with the k-mer length variability. For all k-mer trials, the accuracy remains > 99%, coverage for identification remains < 0.40, and false positives remain ≪ 1. Full results for these simulations can be found in Supplementary Tables S4-S6.

**Fig. 3.**
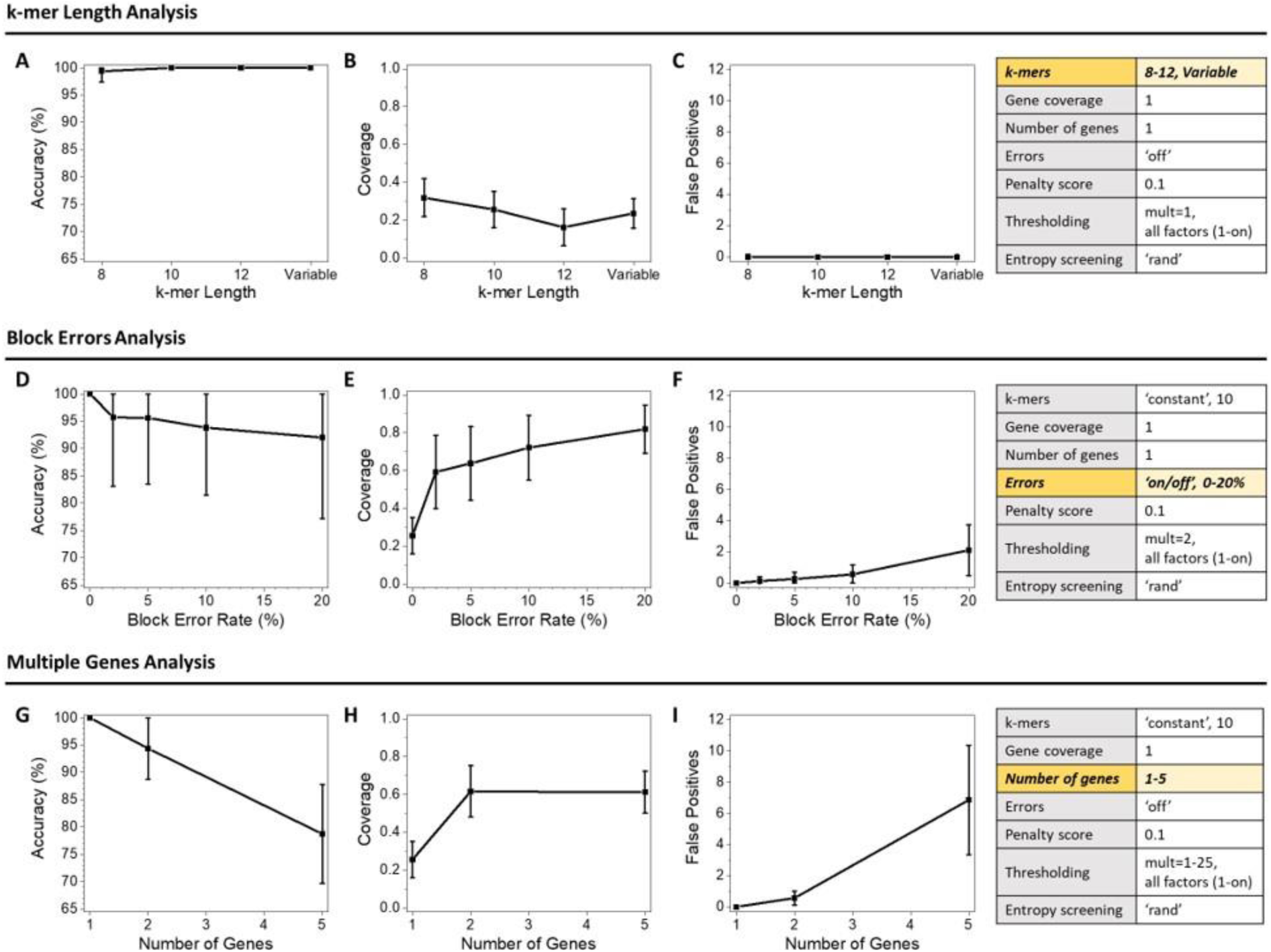
Robust antibiotic resistance gene identification with sequencing variability. (**A. B. C**) The effect on accuracy, coverage for identification, and false positives as k-mer length is varied. For values of k—8, 10, and 12, all blocks are set to length k. For the ‘Variable’ mode, block lengths are sampled from a normal distribution centered around k=10, leading to a distribution of block lengths from ∼6-14. Accuracy, coverage for identification, and false positive rate are all weakly dependent upon k-mer length. For all k-mer trials, the accuracy remains > 99%, coverage remains < 0.40. and false positives remain ≪ 1. (**D. E. F**) The effect on accuracy, coverage for identification, and false positives with errors in the blocks. Even at 20% error rates, the average accuracy remains > 90%, the coverage for identification never reaches 1.0. and false positives are low. (**G. H. I**) The effect on accuracy, coverage for identification, and false positives as blocks from multiple genes are analyzed. Accuracy decreases linearly with an increasing number of genes in the analysis, but remains near 80% for five genes, with average coverage of around 0.60. The main hindrance with an increasing number of genes is the large false positive rate. For the k-mer length and errors analyses in parts A-F, each data point on the graphs is a result of 70 randomly-selected antibiotic resistance genes from the MEGARes database each run through the BOCS simulation with 25 repeats, for a total of 1750 simulations. For the multiple genes analysis in parts G-I, the 2-gene and 5-gene results are from 10 random 2-gene selections and 5 random 5-gene selections from the base set of 70 randomly-selected antibiotic resistance genes, each with 25 repeats.

Next looking at block errors, a set of simulations (for the 70 resistance genes with 25 repeats) were run for each of four error rates within the blocks: 2, 5, 10, and 20%. Note that when using content as a sequencing platform, the error rates become double the rates that would normally be seen in single-letter sequencing. This is because a single point error within a k-mer block affects the resulting content of two nucleotides – the letter corresponding to the correct nucleotide, and the letter corresponding to the incorrect nucleotide. In the BOCS simulation, the error rates are entered as fractional error rates for the gene sequence, not the content; therefore, the error rates shown here (2, 5, 10, and 20%) were entered as 0.01, 0.025, 0.05, and 0.10. The results in Figure 3D-F indicate that accuracy, coverage for identification, and false positive rate are more strongly correlated to block errors than is the k-mer length, although all of these metrics remain strong even under extreme error rates. At error rates as high as 20%, the average accuracy remains > 90%, the coverage for identification never reaches 1.0, and false positives are low (under 2 false positives on average). Full results for these simulations can be found in Supplementary Tables S7-S10.

Lastly looking at using k-mer blocks from multiple genes instead of a single gene (and therefore trying to identify all genes from which the blocks are compiled), we ran two sets of simulations using sets of k-mer blocks from two and five genes. The 2-gene simulations are for 10 random 2-gene selections from the base set of 70 resistance genes, each with 25 repeats. The 5-gene simulations are for 5 random 5-gene selections from the base set of 70 resistance genes, each with 25 repeats. Figure 3G-I shows accuracy decreases linearly with an increasing number of genes, but remains near 80% for five genes, with average coverage around 0.60. The main hindrance with an increasing number of genes is the large false positive rate, which reaches an average of > 6 when the blocks are comprised of five genes. This makes sense when thinking about the relative signal from each gene – when the k-mer blocks are comprised of five different genes, the signal-to-noise level can be as low as 1:4 for each of the genes. The fact that an 80% accuracy rate is observed despite this low signal-to-noise level is impressive, and in the future, more advanced machine learning techniques could be applied to the BOCS algorithm to help reduce the false positive rate. Full results for these simulations can be found in Supplementary Tables S11-S12. In all, the BOCS algorithm proved very robust under the pressures of variable k-mer lengths, high block error rates, and in the presence of blocks comprised of multiple genes.

### 2.4 BOCS for determining clinical MDR bacterial strains

We applied our BOCS simulation towards the detection of a very relevant clinical MDR bacterial strain. Methicillin-resistant *Staphylococcus aureus* (MRSA), has become a leading cause of bacterial infections in healthcare and the community. It is the most clinically-relevant *Staphylococcus* species, with a large prevalence of tissue and bloodstream infections due to chronic skin conditions and surgical procedures. Through horizontal gene transfer, MRSA strains show resistance to most beta-lactam antibiotics, leading to endemics in healthcare facilities worldwide[31,32]. Diagnosis is most commonly performed with phenotypic cell culture assays. These assays look for the presence of the *mecA* gene encoding the PBP2a penicillin-binding protein with a cefoxitin (a beta-lactam, with resistance being of the type OXA class D) antibiotic inducer. The culture tests must incubate for > 24 hours, with overall time for testing usually being > 46 hours[31].

To demonstrate detection of MRSA with BOCS, we designed a simulation looking for two genes: 1) *mecA* gene encoding the PBP2a penicillin-binding protein and 2) OXA beta-lactamase (class D). The simulation used variable length k-mer blocks centered around k=10 (for a range of k=6-14), and a 4% error rate within the blocks. Thresholding (with multiplier and selected factors) and random entropy screening were also applied, and the simulation was run with 50 repeats. The BOCS algorithm once again showed powerful performance in identification of the two resistance genes of interest, leading to MRSA detection even in the presence of block errors and variable k-mer lengths (results in Figure 4). Accuracy was 100%, with identification being made at an average coverage of 0.515 ±0.363. The false positive rate was 4.24 ±3.67. This number is deceptively large, as most of the false positives were genes conferring beta-lactam resistance or general MDR effluxes and would not inhibit proper diagnosis. Figure 4A shows a histogram of the coverage for identification of both the *mecA* and OXA genes throughout all 50 repeats, and Figure 4B shows the specificity as coverage increased. Figure 4C shows increasing content score with coverage, clearly illustrating how the *mecA* and OXA genes of interest probabilistically identify themselves from the rest of the genes in the database. This MDR detection simulation further demonstrates the robustness of the BOCS algorithm and its potential for clinical diagnostics.

**Fig. 4.**
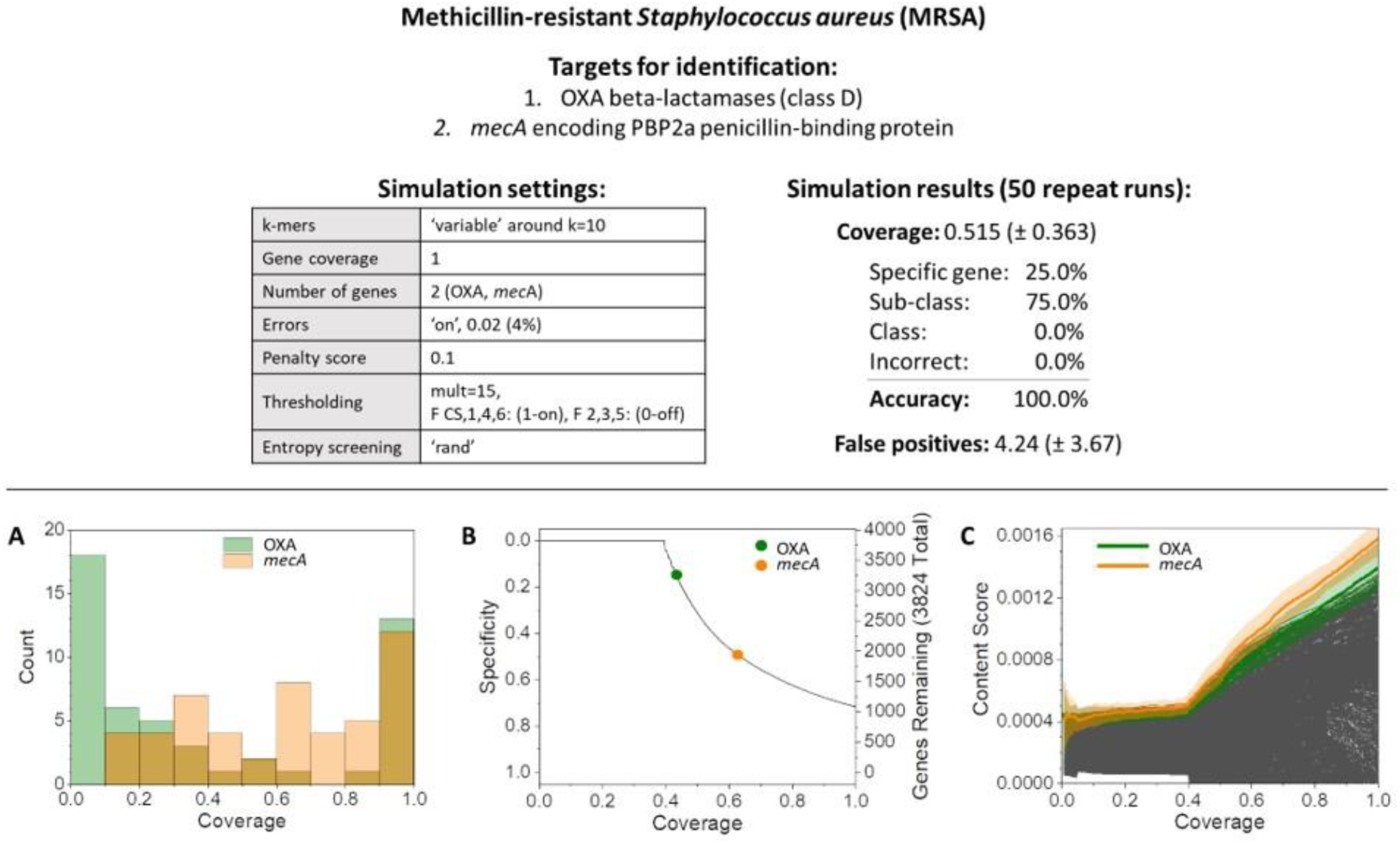
MRSA detection with BOCS. A BOCS simulation was set tip to test the viability of detecting a generic MRS A strain on the basis of two resistance genes (a class D beta-lactamase OXA gene and a *mecA* gene for the penicillin-binding protein PBP2a), which are the norm for both phenotypic and non-phenotypic diagnostic methods. The simulation also included sequencing inconsistencies in the form of variable k-mer block lengths centered around k=10 and a 4% error rate within the blocks. 50 repeat simulations were run for the statistics presented. (**A**) Histogram of the coverage at which the resistance genes are identified in each of the 50 repeat simulations. (**B**) Specificity with increasing coverage. Dots indicate the average coverages at which the OXA and *mecA* genes were identified. The lag where specificity remains at zero during low coverages is a result of a high thresholding multiplier, which was set at 15. (**C**) Increasing content scores for the OXA and *mecA* genes with coverage. The grayed lines are all other genes in the resistance gene database (3822 of the 3824 total). As coverage increases (i.e., as more blocks are analyzed), the genes of interest quickly separate themselves from the others probabilistically, leading to MRSA detection at low coverages.

### 2.5 Applying BOCS to cancer and other genetic disease databases

Expanding BOCS to other areas benefiting from broad-spectrum diagnostics, we ran simulations with the COSMIC cancer database[33] and a custom compiled database of other genetic diseases including many listed by the NIH Undiagnosed Diseases Network (see more information on the custom database in the Supplementary data). Note for these databases, there is no class level identification, only sub-class and specific gene. For each database, 10 randomly-selected genes were run with 10 repeats, for 100 total simulations with constant k-mers at k=10, no block errors, and thresholding and entropy screening (results in Figure 5). Cancer genes (Figure 5A,B) showed 100% accuracy (no false positives) at an average coverage for identification of 0.340 ±0.105 and specificity on par with that of the resistance genes. The other genetic diseases (Figure 5C,D) showed 100% accuracy (no false positives) with an average coverage and specificity significantly better than the resistance genes. The average coverage for identification was 0.132 ±0.136, and roughly 95% of the genes within the database were eliminated by 0.10 coverage. Full results for these simulations can be found in Supplementary Tables S13-S16. The fact that other genetic biomarker databases perform as well or superior to our results with the resistance database adds to the vast potential of the BOCS algorithm in its ability for broad-spectrum diagnostics.

**Fig. 5.**
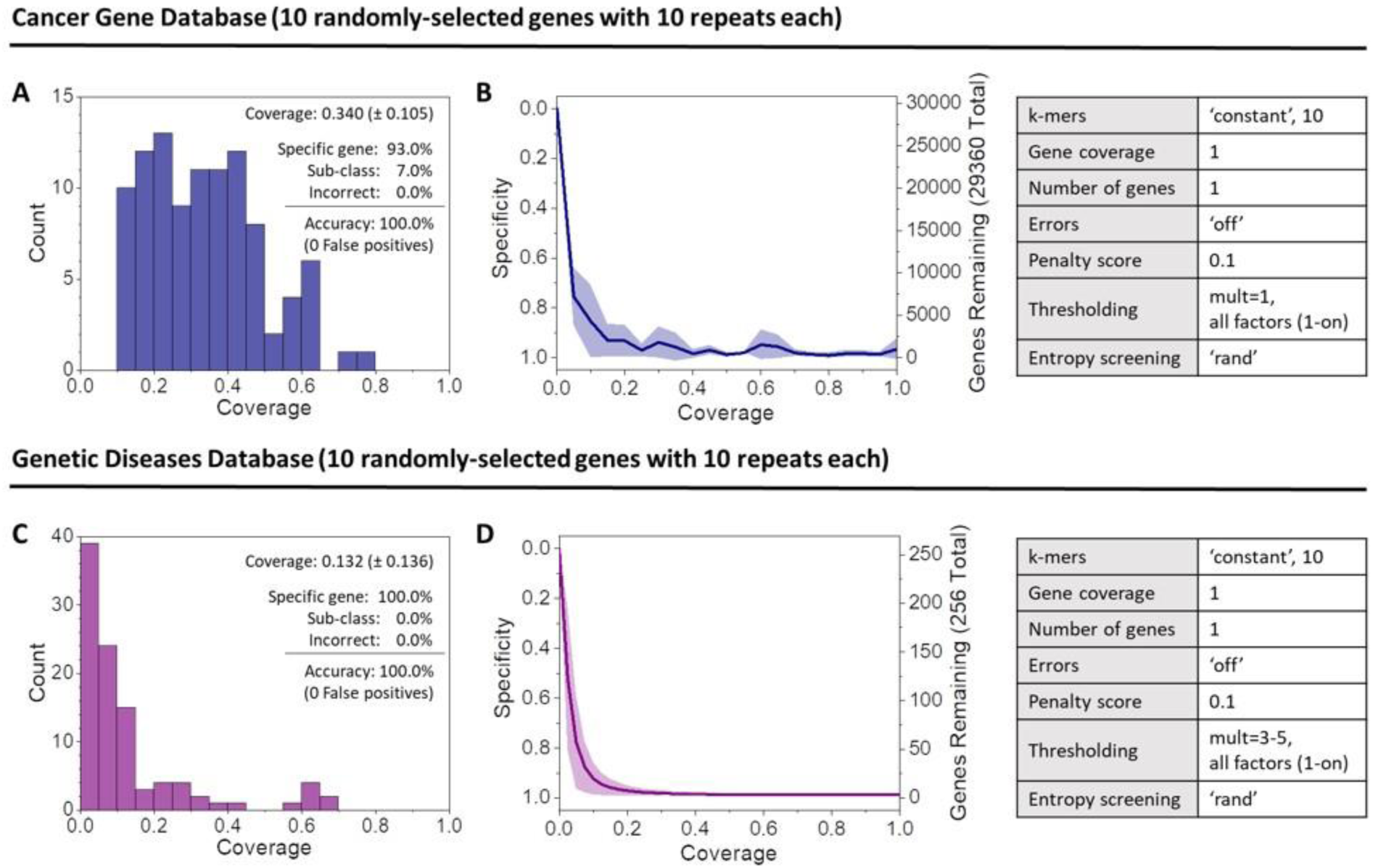
BOCS applied to other genetic biomarkers. To demonstrate the versatility of the BOCS algorithm, simulations were run for identifying single genes from databases for cancer genes (COSMIC database) and other genetic diseases (custom compiled database – see the Supplementary data for more details). For each database, 10 randomly-selected genes were run with 10 repeats, for 100 total simulations. (**A. B**) Histogram of the coverage at which the cancer genes are identified and the specificity with increasing coverage for the cancer genes detection. Accuracy is 100% with an average identification coverage of 0.34, and about 90% of the 29360 genes are eliminated after merely 0.10 coverage. (**C. D**) Histogram of the coverage at which the genetic disease genes are identified and the specificity with increasing coverage for the genetic disease genes detection. Accuracy is 100% with an average identification coverage of only 0.132, and about 95% of the 256 genes are eliminated after just 0.10 coverage.

## 3. Discussion

The results presented here describe a new, powerful bioinformatics tool capable of using DNA k-mer content for rapid genetic biomarker identification, called block optical content scoring (BOCS). The BOCS algorithm uses content-based alignment for probabilistic mapping of k-mer contents to gene sequences within a biomarker database. The algorithm applies elements from pattern recognition and machine learning to rank biomarkers based on a content score. Simulations of the BOCS algorithm showed 100% accurate and highly-specific identification of single antibiotic resistance genes at average coverages of merely 0.255 ±0.096. Further simulations demonstrated robust performance of the BOCS algorithm in the presence of variable k-mer lengths and high sequencing error rates. With errors as high as 20%, over 90% accuracy in gene identification was achieved at less than full gene coverages. Additionally, BOCS has the ability to identify multiple genes when the k-mer fragments from the multiple genes are randomly mixed. When applied to a clinically relevant MDR bacterial strain, the BOCS algorithm showed 100% accuracy with a low false positive rate for detection of two resistance genes (*mecA* and OXA for MRSA identification) at an average coverage of 0.515 ±0.363, with a block error rate of 4% and variable k-mer lengths. BOCS applied to cancer and other genetic diseases also showed detection at 100% accuracy with coverages at or below the values for resistance genes. When coupled with a high-throughput content-based sequencing platform, the BOCS algorithm can provide a biomarker detection tool applicable for rapid, broad-spectrum diagnostics.

## 4. Methods

### 4.1 Entropy screening in the BOCS algorithm

The most significant spikes in raw probabilities occur when the number of permutations for a particular k-mer block is low (i.e., the value *k*!/(*A*! *G*! *C*! *T*!) is low). Preferably analyzing these ‘low entropy’ blocks before others therefore enhances the BOCS algorithm by allowing for genetic biomarker identification at lower coverages, in a process we call entropy screening.

### 4.2 Thresholding in the BOCS algorithm

As more k-mer blocks are analyzed and content scores become compounded, genes within the biomarker database that have probabilistically become irrelevant need to be eliminated. For the case of analyzing k-mer blocks from a single gene at single coverage and no errors, genes can be eliminated when no content matches for a block occur. However, this elimination scheme cannot be implemented in the presence of errors, higher coverages, or the case of multiple genes comprising the k-mer blocks as it will lead to significant decreases in accuracy. To account for this, we implemented a thresholding system within BOCS to remove genes with lowest probability ranks after each consecutive round of block analyses. Thresholding is based on the rank of the content score, as well as each of the individual probability factors, and can be multiplied by a factor to increase/decrease the sensitivity of the eliminations being made.

### 4.3 Accounting for special characters in the genetic databases

Some genetic biomarker database FASTA files contain special nucleic acid code characters (e.g., N signifies that either A, G, C, or T can be substituted into the sequence at that location). When performing content-based sequence alignment, this creates multiple possibilities for content within the two sequences being aligned (the k-mer block and genetic biomarker sequence). To account for these special characters, the BOCS algorithm tests all possible substitutions of A, G, C, and T for the character code used, and a match is awarded if any of the possible substitutions lead to equal content between block and gene sequence.

### 4.4 Making genetic biomarker identifications

The BOCS algorithm uses three levels for gene detection. In the order of most broad to most specific they include – class, sub-class, and specific gene. For example, a gene leading to resistance of beta-lactam antibiotics could have a class: class A beta-lactamase, sub-class: TEM, and specific gene: TEM-x,y,z (where x, y, z are specific mutations of TEM). Based on the level of phylogeny present in the genetic biomarker database, some or all of these classes are used. Each of these levels are tracked in terms of content score ranking throughout the k-mer blocks analysis, and an identification can be made for each level. Identification is determined as the point where a gene within the database adopts one of the n-highest content scores (for n genes comprising the blocks) and remains there and/or separates itself probabilistically from the rest. False positives arise when genes other than the selected gene(s) meet this identification criterion.

### 4.5 Implementing a BOCS simulation

To generate large amounts of data on which to benchmark the BOCS algorithm without the need for experimental data, we built the BOCS algorithm into a simulation. The simulation was implemented in MATLAB R2017b, and is available at https://github.com/lkorshoj/Block-Optical-Content-Scoring. The simulation uses gene sequences from a biomarker database to create k-mer blocks of A-G-C-T content as would be output from high-throughput BOS experiments. The simulated BOS reads are then run through the BOCS algorithm against the biomarker database. The goal of the simulation is to see how well the BOCS algorithm can identify the correct gene (out of all others in the database) using merely randomized k-mer blocks of A-G-C-T content. A specific gene from the database can be pulled or a random gene can be selected. The k-mer block lengths, gene coverage, and the number of errors within the blocks can all be set.

### 4.6 Simulating DNA k-mer blocks

Blocks of DNA k-mer content within the BOCS simulation are generated from one (or more, based on simulation inputs) of the gene sequences within the biomarker database being used. Prior to fragmenting a gene sequence into k-mer blocks, random errors can be added at any specified rate. The gene sequence is split into k-mers based on the set value of k and whether k-mers are to be of constant length or variable length. For the variable length setting, lengths are randomly chosen from a normal distribution centered around the set value for k (with restrictions limiting the length to deviate no more than ±4). Note that the first and last fragments of the gene sequence can deviate from the settings in order to include the entire gene. After errors have been added to the sequence and the gene has been split into k-mers, fractional content for each k-mer is calculated and logged. This process is repeated for however many genes are selected for the analysis and for whatever integer the coverage is set to (for each additional +1X coverage, split locations for the blocks are different). The k-mer block contents for all genes selected for the analysis and all coverages are combined into a single randomized pool to be introduced into the BOCS algorithm. For each repeat simulation, split locations for the k-mer blocks and their randomized ordering will vary.

### 4.7 Simulation inputs/outputs

The following inputs can be set and tuned when running the BOCS simulation (see the Supplementary data for more details):

- Genetic biomarker database – any genetic database in FASTA format
- How simulated k-mers are split (from the overall gene) – at a constant or variable length for k
- Average length of k-mers
- The overall coverage of the gene present throughout all blocks
- The number of genes comprising the blocks
- Error rate within the blocks
- A penalty score is given to genes within the database when no matches to a block are observed
- Multipliers for how sensitive genes within the database are to be eliminated
- Entropy screening method – a randomized or idealized fashion
- The setpoint for what is considered low entropy

The BOCS simulation outputs a text file with the following data used for analysis (see the Supplementary data for more details):

- Simulation runtime, all inputs, and selected gene (gene from the database used to create k-mer blocks)
- All k-mer blocks sequence and content, as well as the randomized order in which they were analyzed in the BOCS algorithm
- Gene coverage as blocks are analyzed
- Specificity as blocks are analyzed
- Classification of the top-ranked genes within the database
- Content scores for all genes in the database

## Availability of code and data

The BOCS simulation MATLAB code, along with genetic biomarker databases used for testing are available at https://github.com/lkorshoj/Block-Optical-Content-Scoring.

## Author contributions

LEK and PN designed the BOCS algorithm. LEK implemented the algorithm and performed simulations. LEK wrote the manuscript with PN making revisions and providing input.

## Acknowledgements

The authors acknowledge computational resources and support from the RMACC Summit Research Computing services at the University of Colorado Boulder.

## Supplementary data

Supplementary data for this article can be found online.

