## Supplementary for "BOCS: DNA k-mer content and scoring for rapid genetic biomarker identification at low coverage"

**Content:**

I. Gene databases
II. Running the BOCS simulation
III. BOCS output
IV. Detailed simulation results
**Table S1.** 70 randomly-selected resistance genes
**Table S2.** Resistance genes simulations with no thresholding or entropy screening
**Table S3.** Resistance genes simulations with thresholding and entropy screening
**Table S4.** Resistance genes simulations with 8-mer blocks
**Table S5.** Resistance genes simulations with 12-mer blocks
**Table S6.** Resistance genes simulations with variable-mer blocks centered around k=10
**Table S7.** Resistance genes simulations with 0.01 (2%) error rate
**Table S8.** Resistance genes simulations with 0.025 (5%) error rate
**Table S9.** Resistance genes simulations with 0.05 (10%) error rate
**Table S10.** Resistance genes simulations with 0.10 (20%) error rate
**Table S11.** Simulations with 2 resistance genes (gene combinations and results)
**Table S12.** Simulations with 5 resistance genes (gene combinations and results)
**Table S13.** 10 randomly-selected cancer genes
**Table S14.** Cancer genes simulations
**Table S15.** 10 randomly-selected genetic disease genes
**Table S16.** Genetic disease genes simulations

### I. Gene databases

The following gene databases were used throughout the study:

1. MEGARes – Antibiotic resistance genes

   Lakin, S. M.; Dean, C.; Noyes, N. R.; Dettenwanger, A.; Ross, A. S.; Doster, E.; Rovira, P.; Abdo, Z.; Jones, K. L.; Ruiz, J.; et al. MEGARes: An Antimicrobial Resistance Database for High Throughput Sequencing. *Nucleic Acids Res.* **2017**, *45* (D1), D574–D580.
2. COSMIC – Cancer gene/somatic mutations

   Forbes, S. A.; Beare, D.; Gunasekaran, P.; Leung, K.; Bindal, N.; Boutselakis, H.; Ding, M.; Bamford, S.; Cole, C.; Ward, S.; et al. COSMIC: Exploring the World's Knowledge of Somatic Mutations in Human Cancer. *Nucleic Acids Res.* **2015**, *43* (D1), D805–D811.
3. Genetic disease genes (custom compiled)

   Database contents…

   - Achondroplasia: FGFR3
   - Alpha-1 antitrypsin deficiency (AATD): SERPINA1
   - Antiphospholipid syndrome (APS): ADAMTS-13
   - Autism: ADNP, ANK2, ARID1B, ASXL3, CACNA1H, CHD2, CHD8, CNTN4, CNTNAP2, CTNND2, DYRK1A, GABRB3, GRIN2B, KDM5B, MECP2, MYT1L, NLGN3, NRXN1, POGZ, PTCHD1, PTEN, RELN, SCN2A, SHANK2, SHANK3, SYNGAP1, TBR1, RPL10, NLGN4X, SNRPN
   - Autosomal dominant polycystic kidney disease: PKD1, PKD2
   - Breast cancer: BRCA1, BRCA2, PALB2, TP53, PTEN, ATM, CDH1, CHEK2, NBN, NF1, STK11, BARD1, BRIP1, CASP8, CTLA4, CYP19A1, FGFR2, H19, LSP1, MAP3K1, MRE11, RAD51, RAD51C, TERT, TOX3, XRCC2, XRCC3
   - Charcot-Marie-Tooth: GARS
   - Colon cancer: APC, MSH2, MLH1, PMS2, MSH6, PMS1
   - Cri du chat: CTNND2, chromosome 5
   - Crohn's disease: ATG16L1, IL23R, IRGM, NOD2, HLA-DRB1, 1L10, 1L12B, JAK2, LRRK2, MUC2, SLC22A4, SLC22A5, STAT3, TYK2
   - Cystic fibrosis: CFTR
   - Dercum disease (a.k.a. Adiposis dolorosa): cause unknown, associated genes unknown
   - Down syndrome: chromosome 21
   - Duane syndrome: CHN1, SALL4
   - Duchenne muscular dystrophy: DMD
   - Factor V Leiden thrombophilia: F5
   - Familial hypercholesterolemia: APOB, LDLR, LDLRAP1, PCSK9
   - Familial Mediterranean fever: MEFV, SAA1
   - Fragile X syndrome: FMR1
   - Gaucher disease: GBA
   - Hemochromatosis: HAMP, HFE, HJV, PNPLA3, SLC40A1, TFR2
   - Hemophilia: F8, F9
   - Holoprosencephaly: DISP1, FGF8, FOXH1, GLIZ, NODAL, PTCH1, SHH, SIX3, TDGF1, ZIC2
   - Huntington disease: HTT
   - Klinefelter syndrome: chromosome x

- Marfan syndrome: FBN1
- Myotonic dystrophy: CNBP, DMPK
- Neurofibromatosis: NF1, NF2
- Noonan syndrome: A2ML1, BRAF, KRAS, LZTR1, MAP2K1, NRAS, PTPN11, RAF1, RASA2, RIT1, RRAS, SOS1, SOS2
- Osteogenesis imperfecta: COL1A1, COL1A2, CRTAP, P3H1
- Parkinson's disease: ATP13A2, GBA, LRRK2, PARK7, PRKN, SNCA, UCHL1, VPS35
- Phenylketonuria: PAH
- Porphyria: ALAD, ALAS2, CPOX, FECH, HFE, HMBS, PPOX, UROD, UROS
- Progeria syndrome: LMNA
- Prostate cancer: AR, BRCA1, BRCA2, CD82, CDH1, CHEK2, EHBP1, ELAC2, EP300, EPHB2, EZH2, FGFR2, FGFR4, GNMT, HNF1B, HOXB13, IGF2, ITGA6, KLF6, LRP2, MAD1L1, MED12, MSMB, MSR1, MXI1, NBN, PCNT, PTEN, RNASEL, SRD5A2, STAT3, TGFBR1, WRN, WT1, ZFHX3
- Retinitis pigmentosa: ABCA4, BEST1, C2orf71, CA4, CERKL, CLRN1, CNGA1, CNGB1, CRB1, CRX, EYS, FAM161A, FSCN2, GUCA1B, IDH3B, IMPDH1, IMPG2, KLH7, LRAT, MERTK, NR2E3, NRL, PDE6A, PDE6B, PDE6G, PRCD, PROM1, PRPF8, PRPF3, PRPF31, PRPH2, RBP3, RDH12, RGR, RHO, RLBP1, ROM1, RP1, RP2, RP9, RPE65, RPGR, SAG, SEMA4A, SNRNP200, SPATA7, TOPORS, TTC8, TULP1, USH2A, WDR19, ZNF513
- Severe combined immunodeficiency (SCID): IL2RG, JAK3, ZAP70
- Sickle cell disease: HBB
- Skin cancer: CDKN2A, CDK4, CDK6, BAP1, BRCA2, PTCH1, PTCH2
- Spinal muscular atrophy: DYNC1H1, SMN1, SMN2, UBA1, VAPB
- Tay-Sachs disease: HEXA
- Thalassemia: HBA1, HBA2, HBB, ATRX
- Trimethylaminuria: FMO3
- Turner syndrome: SHOX
- Velocardiofacial syndrome: COMT, TBX1, chromosome 22
- WAGR syndrome: BDNF, PAX6, WT1, chromosome 11
- Wilson disease: ATP7B, PRNP

### II. Running the BOCS simulation

The following options for inputs/settings are available in BOCS. Within the main text figures, tables are shown summarizing the important inputs that were used for each of the simulations. These include 3, 4, 5, 6, 7, 8, 9 below. The other inputs are not shown in the main text figures, and are merely user options dictating database options, file locations, output settings, and figure displays for further analysis.

1. Choose database – Specify the (1) database type and (2) name of the file. Note that if deviating from the three built-in database types, coding changes must be made. The file must be in the location 'Data/{database_name}/.fasta', and the file must be in the .fasta format.
   Variables to set…
   - g_database
   - database_name
2. Output file location – Specify the folder location for the output .txt file to be written.

Variable to set…

- file_output_loc

3. Length of k-mers – Specify (1) the k-mer splitting method and the (2) k-mer length.
   Variables to set…
   - kmer_split_method – Choose ‘constant’ for k-mers of the same length or ‘variable’ for k-mers of varying length centered around the avg specified by kmer_length, picked from a normal distribution with stdev=2.
   - kmer_length
4. Coverage per nucleotide – Specify the coverage at which each nucleotide in the sequence is seen in the blocks. Breaks for blocks are made in different locations for each additional +1X coverage.
   Variable to set…
   - gene_coverage (must be an integer)
5. Number of genes and select genes– Specify (1) the number of genes from which the blocks are comprised and (2) the number(s) within the database of the specific genes (if any) to use. The genes will be split into blocks and randomized in a batch with blocks from all.
   Variables to set…
   - num_genes (must be an integer)
   - sel_genes – Enter the numbers in an array for the specific genes within the database being used. This is an optional input, as random gene(s) will be selected if nothing is entered. The number of entries in the array must match the value entered for num_genes.
6. Errors – Specify (1) whether random errors should be inserted and (2) the rate at which they are seen. Note that the specified error rate corresponds to the number of random point errors, which is actually only half of the error rate observed in content-based sequencing. So, the actual error rate in the block optical method is double the entered value.
   Variables to set…
   - error_mode – Choose ‘on’ or ‘off’ (in the ‘off’ state, the err_rate is neglected)
   - err_rate
7. Penalty score – Specify the score given to genes when no matches are found for a specific block. It is suggested that a value of 0.1 is used for starting and for most normal analyses.
   Variable to set…
   - penalty_score
8. Thresholding parameters – Specify (1) the multiplier to be multiplied to each of the standard thresholding trends and (2) which of the probability factors to use for thresholding.
   Variables to set…
   - thresh_multiplier – This can be thought of as a sensitivity, where values >1 correspond to a LESS sensitive state (i.e., more genes remain in consideration after each block is analyzed), and values <1 correspond to a MORE sensitive state (i.e., fewer genes remain in consideration after each block is analyzed).
   - thresh_prob_facts_CS – Choose 1/0 (on/off)
   - thresh_prob_facts_F1– Choose 1/0 (on/off)
   - thresh_prob_facts_F2– Choose 1/0 (on/off)
   - thresh_prob_facts_F3 – Choose 1/0 (on/off)
   - thresh_prob_facts_F4 – Choose 1/0 (on/off)

- thresh_prob_facts_F5 – Choose 1/0 (on/off)
- thresh_prob_facts_F6 – Choose 1/0 (on/off)

9. Entropy screening – Specify (1) the entropy screening mode and (2) the threshold for what is considered 'high entropy'.
   Variables to set…
   - entropy_screening_mode – Options are 'rand' for random entropy screening in whichever order the blocks are randomized, 'ideal' for entropy screening idealized from lowest to highest, and 'none' for no entropy screening.
   - perms_thresh – It is suggested to use 10000 as the marker for high entropy since there is a natural break in possible entropy values near this number.
10. Analysis/Troubleshooting/Output options – Specify (1) what kind of analysis is being done, (2) if factor analysis is needed (i.e., figures are displayed for each factor comprising the content score after each block), and (3) the level at which to track gene class.
    Variables to set…
    - analysis_type – Select 'standard' for normal operation and output or 'benchmarking' for extra output including all of the factor values for all of the genes in the database, for establishing new thresholding trends.
    - disp_fact_figs – Choose 'yes' or 'no'
    - tracking_level – This is the number of unique sub-classes of genes with the top content scores after each consecutive block is analyzed. The number used here should be increased as more genes are combined, and helps in analyzing the level of identification of the selected genes (i.e., positive and false positive identifications).

### III. BOCS output

The following sections are output in the results .txt file. The .txt files can be analyzed for overall simulation performance and metrics such as coverage at which the selected gene(s) was identified, accuracy, and false positives.

1. Runtime – Displays the runtime of the content mapping scoring section (i.e., BOCS algorithm)
2. Inputs – Displays all user inputs and options in the following order…
   a. g_database
   b. database_name
   c. file_output_loc
   d. kmer_split_method
   e. kmer_length
   f. gene_coverage
   g. num_genes
   h. sel_genes
   i. err_mode
   j. err_rate
   k. penalty_score
   l. thresh_multiplier
   m. array of thresh_prob_facts_X, where X=CS, F1, F2, F3, F4, F5, F6
   n. entropy_screening_mode

o. perms_thresh
p. analysis_type
q. tracking_level

3. Selected genes data – Information on the selected genes in the study in the order…
    a. Gene number in database
    b. Gene sub-class
    c. Gene class (if the resistance database)
    d. Full gene header/name
    e. Gene sequence
    f. Gene sequence with errors (if err_mode is ‘on’)
4. Blocks for each selected gene(s) – For each coverage of each gene, the columns show…
    a. Block number
    b. Block sequence
    c. A content
    d. G content
    e. C content
    f. T content
5. Randomized blocks – For the combined genes and all coverages, the columns show…
    a. Block number
    b. Gene to which the block belongs
    c. Block sequence
    d. Block entropy
    e. A content
    f. G content
    g. C content
    h. T content
6. Blocks ordered for the final analysis – For the combined genes and all coverages, the columns show…
    a. Block number
    b. Gene to which the block belongs
    c. Block sequence
    d. Block entropy
    e. A content
    f. G content
    g. C content
    h. T content
7. Increasing coverage – Coverage is shown for each individual gene and the overall coverage, with columns showing…
    a. Block number
    b. Coverage for individual genes (each with its own column, 1…num_genes)
    c. Coverage for all genes overall
8. Specificity – Specificity for the overall algorithm with columns…
    a. Block number
    b. Remaining genes (integer)

c. Specificity (fraction in range 0-1)

9. Class analysis – The sub-classes and classes (depending on the database used) of the top content scoring genes after each block is analyzed, with columns in the following order…
    a. Block number
    b. Whether the specific selected gene(s) is identified – 1='yes' and 0='no', there is a column for each selected gene (1…num_genes)
    c. The sub-classes with top content scores (1…tracking_level)
    d. The content scores for the top sub-classes (1…tracking_level)
    e. If the resistance database is being used, The classes with top content scores (1…tracking_level)
    f. If the resistance database is being used, the content scores for the top classes (1…tracking_level)
10. All probability factors (analysis_type='benchmarking' mode only) – All probability factors (and some slope analyses) are output for each block for each gene in the database in a matrix with dimensions (number genes × number blocks +1), with columns…
    a. Gene number
    b. Cumulative probability factor (or slope analysis) for each block (1…number of blocks)
11. All content scores – All content scores are output for each block for each gene in the database in a matrix with dimensions (number genes × number blocks +1), with columns…
    a. Gene number
    b. Content score (1…number of blocks)
12. Content scores extracted for the selected genes – Content scores with columns…
    a. Block number
    b. Content score for selected genes (1…num_genes)

### IV. Detailed simulation results

The following pages include tables of detailed results for the figures presented in the main text. This includes information on all of the individual genes used in the simulations, as well as full simulation results for single-gene studies with and without entropy screening, varying k-mer lengths, and block errors; multiple-gene studies; and cancer and other genetic disease results.

**Table S1.** 70 randomly-selected resistance genes

| No. | Gene database No. | Sub-class | Class | Full gene name (from MEGARes database) |
|---|---|---|---|---|
| 1 | 4 | VANZA | VanA-type_accessory_protein | 959\|M97297.1\|TRNVAN\|Glycopeptides\|VanA-type_accessory_protein\|VANZA |
| 2 | 52 | VANWG | VanG-type_accessory_protein | Gly\|VanW-G_1_AY271782\|Glycopeptides\|VanG-type_accessory_protein\|VANWG |
| 3 | 62 | CTX | Class_A_betalactamases | Bla\|CTX-M-9\|AF174129\|1-876\|876\|betalactams\|Class_A_betalactamases\|CTX |
| 4 | 68 | MEFA | Macrolide_resistance_efflux_pumps | MLS\|NC_023287.1.18156494\|MLS\|Macrolide_resistance_efflux_pumps\|MEFA |
| 5 | 76 | OXA | Class_D_betalactamases | Bla\|OXA-208\|FR853176\|1-825\|825\|betalactams\|Class_D_betalactamases\|OXA |
| 6 | 92 | CATB | Chloramphenicol_acetyltransferases | Phe\|FJ460181.2\|gene2\|Phenicol\|Chloramphenicol_acetyltransferases\|CATB\|RequiresSNPConfirmation |
| 7 | 162 | EREA | Macrolide_esterases | MLS\|ereA_4_AF512546\|MLS\|Macrolide_esterases\|EREA |
| 8 | 174 | TEM | Class_A_betalactamases | Bla\|TEM-59\|AF062386\|31-862\|832\|betalactams\|Class_A_betalactamases\|TEM |
| 9 | 193 | CML | Phenicol_efflux_pumps | 1597\|HQ713678.1\|HQ713678\|Phenicol\|Phenicol_efflux_pumps\|CML |
| 10 | 207 | NIMA | nim_nitroimidazole_reductase | Met\|nimH_1_FJ969397\|Metronidazole\|nim_nitroimidazole_reductase\|NIMA |
| 11 | 239 | DHFR | Dihydrofolate_reductase | Tmt\|dfrA1_2_AJ419168\|Trimethoprim\|Dihydrofolate_reductase\|DHFR\|RequiresSNPConfirmation |
| 12 | 246 | FOLP | Sulfonamide-resistant_dihydropteroate_synthases | Sul\|CP001581.1\|gene938\|Sulfonamides\|Sulfonamide-resistant_dihydropteroate_synthases\|FOLP\|RequiresSNPConfirmation |
| 13 | 274 | PARC | Fluoroquinolone-resistant_DNA_topoisomerases | Flq\|NC_011586.7046300\|Fluoroquinolones\|Fluoroquinolone-resistant_DNA_topoisomerases\|PARC\|RequiresSNPConfirmation |
| 14 | 276 | SHV | Class_A_betalactamases | gi\|28912444\|gb\|AY210887.1\|betalactams\|Class_A_betalactamases\|SHV |
| 15 | 284 | DFRA | Dihydrofolate_reductase | CARD\|phgb\|AM403715\|302-854\|ARO:3002857\|dfrA26\|Trimethoprim\|Dihydrofolate_reductase\|DFRA |
| 16 | 295 | SOXS | MDR_regulator | CARD\|pvgb\|NC_003197\|4503969-4504293\|ARO:3003383\|Salmonella\|Multi-drug_resistance\|MDR_regulator\|SOXS\|RequiresSNPConfirmation |
| 17 | 321 | TSNR | Thiostrepton_23S_rRNA_methyltransferases | CARD\|phgb\|AL123456\|1853605-1854388\|ARO:3003060\|tsnr\|Thiostrepton\|Thiostrepton_23S_rRNA_methyltransferases\|TSNR |
| 18 | 413 | OMPD | Mutant_porin_proteins | Bla\|CP004022.1\|gene834\|betalactams\|Mutant_porin_proteins\|OMPD |
| 19 | 486 | LMRA | Lincomycin-resistant_lmrA | CARD\|pvgb\|AL009126\|290131-290698\|ARO:3003028\|lmrA\|MLS\|Lincomycin-resistant_lmrA\|LMRA\|RequiresSNPConfirmation |

| | | | | |
|---|---|---|---|---|
| 20 | 505 | CARB | Class_A_betalactamases | CARD\|phgb\|HF953351\|2461-3358\|ARO:3002255\|CARB-16\|betalactams\|Class_A_betalactamases\|CARB |
| 21 | 506 | ANT3-DPRIME | Aminoglycoside_O-nucleotidyltransferases | CARD\|phgb\|NC_010410\|3621491-3622283\|ARO:3002601\|aadA\|Aminoglycosides\|Aminoglycoside_O-nucleotidyltransferases\|ANT3-DPRIME |
| 22 | 555 | QNRS | Quinolone_resistance_protein_Qnr | Flq\|QnrS6_1_HQ631376\|Fluoroquinolones\|Quinolone_resistance_protein_Qnr\|QNRS |
| 23 | 558 | VANWI | VanI-type_accessory_protein | CARD\|phgb\|NC_007907.1\|2195400-2196522\|ARO:3003724\|vanWI\|Glycopeptides\|VanI-type_accessory_protein\|VANWI |
| 24 | 578 | TETX | Tetracycline_inactivation_enzymes | Tet\|tetX_1_GU014535\|Tetracyclines\|Tetracycline_inactivation_enzymes\|TETX |
| 25 | 652 | VGA | ABC_transporter | 105\|FR772051.1\|FR772051\|MLS\|ABC_transporter\|VGA |
| 26 | 682 | MOX | Class_C_betalactamases | Bla\|MOX-2\|AJ276453\|4620-5768\|1149\|betalactams\|Class_C_betalactamases\|MOX |
| 27 | 694 | ACT | Class_C_betalactamases | gi\|595583477\|gb\|KF992026.1\|betalactams\|Class_C_betalactamases\|ACT |
| 28 | 717 | VANRE | VanE-type_regulator | CARD\|phgb\|FJ872411\|43513-44203\|ARO:3002924\|vanRE\|Glycopeptides\|VanE-type_regulator\|VANRE |
| 29 | 749 | RMTB | 16S_rRNA_methyltransferases | CARD\|phgb\|FJ483516.1\|0-252\|ARO:3000860\|rmtB\|Aminoglycosides\|16S_rRNA_methyltransferases\|RMTB |
| 30 | 778 | DHA | Class_C_betalactamases | gi\|698174199\|gb\|KM087854.1\|betalactams\|Class_C_betalactamases\|DHA |
| 31 | 789 | CMY | Class_C_betalactamases | Bla\|CMY-4\|AF420597\|1-1146\|1146\|betalactams\|Class_C_betalactamases\|CMY |
| 32 | 797 | FACT | Elfamycin_efflux_pumps | CARD\|phgb\|JQ768046\|7760-9440\|ARO:3001313\|facT\|Elfamycins\|Elfamycin_efflux_pumps\|FACT |
| 33 | 819 | OKP | Class_A_betalactamases | Bla\|OKP-A-3\|AM051140\|1-861\|861\|betalactams\|Class_A_betalactamases\|OKP |
| 34 | 973 | IMP | Class_B_betalactamases | 1570\|LC031883.1\|LC031883\|betalactams\|Class_B_betalactamases\|IMP |
| 35 | 1048 | VIM | Class_B_betalactamases | Bla\|VIM-37\|JX982636\|1-801\|801\|betalactams\|Class_B_betalactamases\|VIM |
| 36 | 1135 | PBP1B | Penicillin_binding_protein | CARD\|phgb\|NC_003098\|1886035-1888501\|ARO:3003044\|PBP1b\|betalactams\|Penicillin_binding_protein\|PBP1B |
| 37 | 1146 | MPHE | Macrolide_phosphotransferases | 1507\|unknown_id\|unknown_name\|MLS\|Macrolide_phosphotransferases\|MPHE |
| 38 | 1182 | VATE | Streptogramin_A_O-acetyltransferase | MLS\|vatE_3_AF153312\|MLS\|Streptogramin_A_O-acetyltransferase\|VATE |
| 39 | 1214 | OPRJ | MDR_mutant_porin_proteins | 155\|U57969.1\|PAU57969\|Multi-drug_resistance\|MDR_mutant_porin_proteins\|OPRJ |
| 40 | 1254 | FOSB | Fosfomycin_thiol_transferases | Fos\|Fcyn\|FosB\|AHLO01000073\|63139-63558\|417\|Fosfomycin\|Fosfomycin_thiol_transferases\|FOSB |

| | | | | |
|---|---|---|---|---|
| 41 | 1271 | VPH | Viomycin_phosphotransferases | CARD\|phgb\|X02393\|96-960\|ARO:3003061\|viomycin\|Mycobacterium_tuberculosis-specific_Drug\|Viomycin_phosphotransferases\|VPH |
| 42 | 1283 | SULI | Sulfonamide-resistant_dihydropteroate_synthases | Sul\|sul1_22_AY115475\|Sulfonamides\|Sulfonamide-resistant_dihydropteroate_synthases\|SULI |
| 43 | 1297 | TET40 | Tetracycline_resistance_ribosomal_protection_proteins | Tet\|JQ740052.1\|gene2\|Tetracyclines\|Tetracycline_resistance_ribosomal_protection_proteins\|TET40 |
| 44 | 1389 | CPXAR | MDR_regulator | Mdr\|CP000034.1\|gene3834\|Multi-drug_resistance\|MDR_regulator\|CPXAR |
| 45 | 1392 | AAC6-PRIME | Aminoglycoside_N-acetyltransferases | AGly\|Aac6-32\|EF614235\|2247-2801\|555\|Aminoglycosides\|Aminoglycoside_N-acetyltransferases\|AAC6-PRIME |
| 46 | 1422 | VGBB | Streptogramin_B_ester_bond_cleavage | CARD\|phgb\|AF015628\|398-1286\|ARO:3001308\|VgbB\|MLS\|Streptogramin_B_ester_bond_cleavage\|VGBB |
| 47 | 1440 | FOSC | Fosfomycin_phosphorylation | CARD\|phgb\|Z33413\|386-935\|ARO:3000380\|FosC\|Fosfomycin\|Fosfomycin_phosphorylation\|FOSC |
| 48 | 1535 | LNUA | Lincosamide_nucleotidyltransferases | MLS\|AM399080.1\|gene2\|MLS\|Lincosamide_nucleotidyltransferases\|LNUA |
| 49 | 1569 | PARE | Aminocoumarin-resistant_DNA_topoisomerases | ACou\|CP000675.2\|gene802\|Aminocoumarins\|Aminocoumarin-resistant_DNA_topoisomerases\|PARE\|RequiresSNPConfirmation |
| 50 | 1695 | NDM | Class_B_betalactamases | CARD\|phgb\|FN396876\|2406-3219\|ARO:3000589\|NDM-1\|betalactams\|Class_B_betalactamases\|NDM |
| 51 | 1702 | SPG | Class_B_betalactamases | CARD\|phgb\|KP109680\|1254-2112\|ARO:3003720\|SPG-1\|betalactams\|Class_B_betalactamases\|SPG |
| 52 | 1753 | VEB | Class_A_betalactamases | Bla\|VEB-1_1_HM370393\|betalactams\|Class_A_betalactamases\|VEB |
| 53 | 1953 | LNUB | Lincosamide_nucleotidyltransferases | MLS\|LnuB\|AJ238249\|127-930\|804\|MLS\|Lincosamide_nucleotidyltransferases\|LNUB |
| 54 | 2026 | ERMA | 23S_rRNA_methyltransferases | 126\|AJ579365.1\|AJ579365\|MLS\|23S_rRNA_methyltransferases\|ERMA |
| 55 | 2357 | SULII | Sulfonamide-resistant_dihydropteroate_synthases | Sul\|sul2_12_AF497970\|Sulfonamides\|Sulfonamide-resistant_dihydropteroate_synthases\|SULII |
| 56 | 2517 | TET37 | Tetracycline_inactivation_enzymes | CARD\|phgb\|AF540889\|0-327\|ARO:3002871\|tet37\|Tetracyclines\|Tetracycline_inactivation_enzymes\|TET37 |
| 57 | 2822 | EMRK | Multi-drug_efflux_pumps | CARD\|phgb\|D78168\|536-1592\|ARO:3000206\|emrK\|Multi-drug_resistance\|Multi-drug_efflux_pumps\|EMRK |
| 58 | 2999 | MPHB | Macrolide_phosphotransferases | 424\|D85892.1\|D85892\|MLS\|Macrolide_phosphotransferases\|MPHB |
| 59 | 3024 | VANYM | VanM-type_accessory_protein | 299\|FJ349556.1\|FJ349556\|Glycopeptides\|VanM-type_accessory_protein\|VANYM |
| 60 | 3041 | MECC | Penicillin_binding_protein | CARD\|phgb\|AB037671\|24420-26427\|ARO:3001209\|mecC\|betalactams\|Penicillin_binding_protein\|MECC |
| 61 | 3128 | TUFAB | EF-Tu_inhibition | Elf\|CP000647.1\|gene3761\|Elfamycins\|EF-Tu_inhibition\|TUFAB\|RequiresSNPConfirmation |

| | | | | |
|---|---|---|---|---|
| 62 | 3176 | AMRB | Multi-drug_efflux_pumps | CARD\|phgb\|NC_002516\|2208168-2211306\|ARO:3002983\|amrB\|Multi-drug_resistance\|Multi-drug_efflux_pumps\|AMRB |
| 63 | 3270 | IRI | Monooxygenase | CARD\|phgb\|U56415\|279-1719\|ARO:3002884\|iri\|Rifampin\|Monooxygenase\|IRI |
| 64 | 3314 | RPOB | Rifampin-resistant_beta-subunit_of_RNA_polymerase_RpoB | Rif\|NC_002758.1120515\|Rifampin\|Rifampin-resistant_beta-subunit_of_RNA_polymerase_RpoB\|RPOB\|RequiresSNPConfirmation |
| 65 | 3332 | TET35 | Tetracycline_resistance_major_facilitator_superfamily_MFS_efflux_pumps | CARD\|phgb\|AF353562\|0-1110\|ARO:3000481\|tet35\|Tetracyclines\|Tetracycline_resistance_major_facilitator_superfamily_MFS_efflux_pumps\|TET35 |
| 66 | 3370 | CFRA | Florfenicol_methyltransferases | MLS\|CfrA\|AM408573\|10028-11077\|1050\|Phenicol\|Florfenicol_methyltransferases\|CFRA |
| 67 | 3513 | BRP | Bleomycin_resistance_protein | CARD\|phgb\|NC_012547\|21239-21638\|ARO:3001205\|bleomycin\|Glycopeptides\|Bleomycin_resistance_protein\|BRP |
| 68 | 3613 | APH3-PRIME | Aminoglycoside_O-phosphotransferases | AGly\|APH-Stph\|HE579073\|1778413-1779213\|801\|Aminoglycosides\|Aminoglycoside_O-phosphotransferases\|APH3-PRIME |
| 69 | 3697 | TETM | Tetracycline_resistance_ribosomal_protection_proteins | Tet\|tetM_6_M21136\|Tetracyclines\|Tetracycline_resistance_ribosomal_protection_proteins\|TETM |
| 70 | 3778 | IND | Class_B_betalactamases | Bla\|IND-11\|HM245379\|57-788\|732\|betalactams\|Class_B_betalactamases\|IND |

These 70 genes were used throughout all studies and simulations with the MEGARes antibiotic resistance database.

**Table S2.** Resistance genes simulations with no thresholding or entropy screening

| No. | Database No. | Gene sub-class | Avg block for identification | StDev block for identification | Avg coverage for identification | StDev coverage for identification | Accuracy | Identification level (fraction of trials) | | | | Avg false positives | StDev false positives |
|---|---|---|---|---|---|---|---|---|---|---|---|---|---|
| | | | | | | | | Specific gene | Sub-class | Class | Incorrect | | |
| 1 | 4 | VANZA | 15.240 | 7.655 | 0.310 | 0.156 | 1.000 | 1.000 | 0.000 | 0.000 | 0.000 | 0.000 | 0.000 |
| 2 | 52 | VANWG | 20.240 | 7.721 | 0.238 | 0.091 | 1.000 | 0.280 | 0.720 | 0.000 | 0.000 | 0.000 | 0.000 |
| 3 | 62 | CTX | 26.560 | 12.553 | 0.302 | 0.142 | 1.000 | 0.000 | 1.000 | 0.000 | 0.000 | 0.000 | 0.000 |
| 4 | 68 | MEFA | 28.080 | 11.169 | 0.229 | 0.090 | 1.000 | 0.400 | 0.600 | 0.000 | 0.000 | 0.000 | 0.000 |
| 5 | 76 | OXA | 20.360 | 9.827 | 0.246 | 0.119 | 1.000 | 0.000 | 1.000 | 0.000 | 0.000 | 0.000 | 0.000 |
| 6 | 92 | CATB | 15.920 | 8.093 | 0.249 | 0.127 | 1.000 | 0.520 | 0.480 | 0.000 | 0.000 | 0.000 | 0.000 |
| 7 | 162 | EREA | 31.360 | 13.187 | 0.254 | 0.106 | 1.000 | 0.080 | 0.920 | 0.000 | 0.000 | 0.000 | 0.000 |
| 8 | 174 | TEM | 25.720 | 9.965 | 0.306 | 0.119 | 1.000 | 0.000 | 1.000 | 0.000 | 0.000 | 0.000 | 0.000 |
| 9 | 193 | CML | 17.040 | 8.904 | 0.129 | 0.068 | 1.000 | 0.000 | 1.000 | 0.000 | 0.000 | 0.000 | 0.000 |
| 10 | 207 | NIMA | 13.560 | 7.235 | 0.292 | 0.154 | 1.000 | 1.000 | 0.000 | 0.000 | 0.000 | 0.000 | 0.000 |
| 11 | 239 | DHFR | 13.680 | 5.949 | 0.285 | 0.125 | 1.000 | 0.280 | 0.720 | 0.000 | 0.000 | 0.000 | 0.000 |
| 12 | 246 | FOLP | 28.680 | 10.664 | 0.240 | 0.090 | 1.000 | 1.000 | 0.000 | 0.000 | 0.000 | 0.000 | 0.000 |
| 13 | 274 | PARC | 36.040 | 11.374 | 0.162 | 0.051 | 1.000 | 0.400 | 0.600 | 0.000 | 0.000 | 0.000 | 0.000 |
| 14 | 276 | SHV | 24.760 | 11.399 | 0.283 | 0.130 | 1.000 | 0.040 | 0.960 | 0.000 | 0.000 | 0.000 | 0.000 |
| 15 | 284 | DFRA | 18.440 | 7.932 | 0.331 | 0.142 | 1.000 | 1.000 | 0.000 | 0.000 | 0.000 | 0.000 | 0.000 |
| 16 | 295 | SOXS | 15.120 | 4.494 | 0.456 | 0.135 | 1.000 | 1.000 | 0.000 | 0.000 | 0.000 | 0.000 | 0.000 |
| 17 | 321 | TSNR | 23.840 | 8.030 | 0.302 | 0.101 | 1.000 | 1.000 | 0.000 | 0.000 | 0.000 | 0.000 | 0.000 |
| 18 | 413 | OMPD | 26.440 | 9.739 | 0.244 | 0.090 | 1.000 | 1.000 | 0.000 | 0.000 | 0.000 | 0.000 | 0.000 |
| 19 | 486 | LMRA | 10.760 | 4.807 | 0.187 | 0.084 | 1.000 | 1.000 | 0.000 | 0.000 | 0.000 | 0.000 | 0.000 |
| 20 | 505 | CARB | 21.400 | 8.436 | 0.236 | 0.093 | 1.000 | 1.000 | 0.000 | 0.000 | 0.000 | 0.000 | 0.000 |
| 21 | 506 | ANT3-DPRIME | 19.320 | 6.606 | 0.242 | 0.083 | 1.000 | 0.920 | 0.080 | 0.000 | 0.000 | 0.000 | 0.000 |
| 22 | 555 | QNRS | 17.080 | 8.067 | 0.258 | 0.122 | 1.000 | 0.360 | 0.640 | 0.000 | 0.000 | 0.000 | 0.000 |
| 23 | 558 | VANWI | 20.640 | 9.591 | 0.183 | 0.085 | 1.000 | 1.000 | 0.000 | 0.000 | 0.000 | 0.000 | 0.000 |
| 24 | 578 | TETX | 30.360 | 13.391 | 0.259 | 0.115 | 1.000 | 0.280 | 0.720 | 0.000 | 0.000 | 0.000 | 0.000 |
| 25 | 652 | VGA | 38.120 | 14.001 | 0.242 | 0.089 | 1.000 | 0.880 | 0.120 | 0.000 | 0.000 | 0.000 | 0.000 |
| 26 | 682 | MOX | 42.960 | 14.226 | 0.372 | 0.123 | 1.000 | 0.400 | 0.600 | 0.000 | 0.000 | 0.000 | 0.000 |
| 27 | 694 | ACT | 41.480 | 18.136 | 0.330 | 0.145 | 1.000 | 0.120 | 0.880 | 0.000 | 0.000 | 0.000 | 0.000 |
| 28 | 717 | VANRE | 23.160 | 8.740 | 0.331 | 0.124 | 1.000 | 1.000 | 0.000 | 0.000 | 0.000 | 0.000 | 0.000 |
| 29 | 749 | RMTB | 8.320 | 3.859 | 0.315 | 0.151 | 1.000 | 1.000 | 0.000 | 0.000 | 0.000 | 0.000 | 0.000 |
| 30 | 778 | DHA | 23.760 | 11.921 | 0.208 | 0.104 | 1.000 | 0.000 | 1.000 | 0.000 | 0.000 | 0.000 | 0.000 |
| 31 | 789 | CMY | 31.160 | 16.570 | 0.271 | 0.144 | 1.000 | 0.000 | 1.000 | 0.000 | 0.000 | 0.000 | 0.000 |
| 32 | 797 | FACT | 25.440 | 16.269 | 0.151 | 0.096 | 1.000 | 1.000 | 0.000 | 0.000 | 0.000 | 0.000 | 0.000 |
| 33 | 819 | OKP | 30.840 | 15.763 | 0.355 | 0.181 | 1.000 | 0.040 | 0.960 | 0.000 | 0.000 | 0.000 | 0.000 |
| 34 | 973 | IMP | 20.240 | 10.849 | 0.270 | 0.146 | 1.000 | 0.200 | 0.800 | 0.000 | 0.000 | 0.000 | 0.000 |
| 35 | 1048 | VIM | 18.640 | 9.420 | 0.230 | 0.116 | 1.000 | 0.040 | 0.960 | 0.000 | 0.000 | 0.000 | 0.000 |
| 36 | 1135 | PBP1B | 34.640 | 13.997 | 0.140 | 0.056 | 1.000 | 1.000 | 0.000 | 0.000 | 0.000 | 0.000 | 0.000 |
| 37 | 1146 | MPHE | 23.000 | 10.794 | 0.259 | 0.122 | 1.000 | 0.400 | 0.600 | 0.000 | 0.000 | 0.000 | 0.000 |
| 38 | 1182 | VATE | 16.760 | 6.597 | 0.258 | 0.102 | 1.000 | 0.600 | 0.400 | 0.000 | 0.000 | 0.000 | 0.000 |
| 39 | 1214 | OPRJ | 49.200 | 25.987 | 0.339 | 0.179 | 1.000 | 1.000 | 0.000 | 0.000 | 0.000 | 0.000 | 0.000 |
| 40 | 1254 | FOSB | 14.600 | 5.370 | 0.343 | 0.124 | 1.000 | 1.000 | 0.000 | 0.000 | 0.000 | 0.000 | 0.000 |
| 41 | 1271 | VPH | 28.960 | 16.592 | 0.333 | 0.191 | 1.000 | 1.000 | 0.000 | 0.000 | 0.000 | 0.000 | 0.000 |
| 42 | 1283 | SULI | 26.160 | 11.912 | 0.309 | 0.140 | 1.000 | 0.120 | 0.880 | 0.000 | 0.000 | 0.000 | 0.000 |
| 43 | 1297 | TET40 | 19.960 | 8.801 | 0.162 | 0.071 | 1.000 | 0.480 | 0.520 | 0.000 | 0.000 | 0.000 | 0.000 |
| 44 | 1389 | CPXAR | 18.720 | 8.483 | 0.265 | 0.121 | 1.000 | 0.520 | 0.480 | 0.000 | 0.000 | 0.000 | 0.000 |
| 45 | 1392 | AAC6-PRIME | 11.440 | 5.973 | 0.204 | 0.106 | 1.000 | 0.720 | 0.280 | 0.000 | 0.000 | 0.000 | 0.000 |
| 46 | 1422 | VGBB | 32.960 | 10.737 | 0.367 | 0.120 | 1.000 | 1.000 | 0.000 | 0.000 | 0.000 | 0.000 | 0.000 |
| 47 | 1440 | FOSC | 16.400 | 8.201 | 0.293 | 0.146 | 1.000 | 1.000 | 0.000 | 0.000 | 0.000 | 0.000 | 0.000 |

| | | | | | | | | | | | | | |
|---|---|---|---|---|---|---|---|---|---|---|---|---|---|
| 48 | 1535 | LNUA | 12.160 | 6.743 | 0.249 | 0.138 | 1.000 | 0.520 | 0.480 | 0.000 | 0.000 | 0.000 | 0.000 |
| 49 | 1569 | PARE | 50.680 | 20.954 | 0.268 | 0.111 | 1.000 | 0.680 | 0.320 | 0.000 | 0.000 | 0.000 | 0.000 |
| 50 | 1695 | NDM | 17.960 | 10.990 | 0.218 | 0.133 | 1.000 | 0.960 | 0.040 | 0.000 | 0.000 | 0.000 | 0.000 |
| 51 | 1702 | SPG | 23.480 | 14.726 | 0.273 | 0.171 | 1.000 | 1.000 | 0.000 | 0.000 | 0.000 | 0.000 | 0.000 |
| 52 | 1753 | VEB | 26.640 | 14.059 | 0.294 | 0.155 | 1.000 | 0.120 | 0.880 | 0.000 | 0.000 | 0.000 | 0.000 |
| 53 | 1953 | LNUB | 24.000 | 9.069 | 0.296 | 0.112 | 1.000 | 0.840 | 0.160 | 0.000 | 0.000 | 0.000 | 0.000 |
| 54 | 2026 | ERMA | 21.160 | 10.172 | 0.286 | 0.137 | 1.000 | 0.920 | 0.080 | 0.000 | 0.000 | 0.000 | 0.000 |
| 55 | 2357 | SULII | 27.960 | 8.763 | 0.342 | 0.108 | 1.000 | 0.200 | 0.800 | 0.000 | 0.000 | 0.000 | 0.000 |
| 56 | 2517 | TET37 | 11.040 | 4.335 | 0.332 | 0.133 | 1.000 | 1.000 | 0.000 | 0.000 | 0.000 | 0.000 | 0.000 |
| 57 | 2822 | EMRK | 30.440 | 11.285 | 0.287 | 0.106 | 1.000 | 1.000 | 0.000 | 0.000 | 0.000 | 0.000 | 0.000 |
| 58 | 2999 | MPHB | 26.960 | 11.156 | 0.295 | 0.123 | 1.000 | 1.000 | 0.000 | 0.000 | 0.000 | 0.000 | 0.000 |
| 59 | 3024 | VANYM | 25.000 | 8.401 | 0.351 | 0.118 | 1.000 | 1.000 | 0.000 | 0.000 | 0.000 | 0.000 | 0.000 |
| 60 | 3041 | MECC | 67.560 | 60.175 | 0.336 | 0.299 | 1.000 | 1.000 | 0.000 | 0.000 | 0.000 | 0.000 | 0.000 |
| 61 | 3128 | TUFAB | 30.400 | 14.543 | 0.255 | 0.122 | 1.000 | 0.680 | 0.320 | 0.000 | 0.000 | 0.000 | 0.000 |
| 62 | 3176 | AMRB | 67.960 | 30.847 | 0.216 | 0.098 | 1.000 | 1.000 | 0.000 | 0.000 | 0.000 | 0.000 | 0.000 |
| 63 | 3270 | IRI | 53.080 | 22.156 | 0.366 | 0.153 | 1.000 | 1.000 | 0.000 | 0.000 | 0.000 | 0.000 | 0.000 |
| 64 | 3314 | RPOB | 40.120 | 14.661 | 0.113 | 0.041 | 1.000 | 0.000 | 1.000 | 0.000 | 0.000 | 0.000 | 0.000 |
| 65 | 3332 | TET35 | 19.920 | 10.352 | 0.178 | 0.093 | 1.000 | 1.000 | 0.000 | 0.000 | 0.000 | 0.000 | 0.000 |
| 66 | 3370 | CFRA | 35.000 | 13.994 | 0.331 | 0.133 | 1.000 | 1.000 | 0.000 | 0.000 | 0.000 | 0.000 | 0.000 |
| 67 | 3513 | BRP | 10.920 | 3.121 | 0.270 | 0.078 | 1.000 | 1.000 | 0.000 | 0.000 | 0.000 | 0.000 | 0.000 |
| 68 | 3613 | APH3-PRIME | 17.640 | 8.602 | 0.218 | 0.105 | 1.000 | 1.000 | 0.000 | 0.000 | 0.000 | 0.000 | 0.000 |
| 69 | 3697 | TETM | 55.200 | 26.608 | 0.286 | 0.138 | 1.000 | 0.480 | 0.520 | 0.000 | 0.000 | 0.000 | 0.000 |
| 70 | 3778 | IND | 23.840 | 7.548 | 0.322 | 0.102 | 1.000 | 0.040 | 0.960 | 0.000 | 0.000 | 0.000 | 0.000 |

Simulation settings:

k-mers: ‘constant’, k=10
Gene coverage: 1
Number of genes: 1
Errors: ‘off’
Penalty score: 0.1
Thresholding and entropy screening was deactivated

**Table S3.** Resistance genes simulations with thresholding and entropy screening

| No. | Database No. | Gene sub-class | Avg block for identification | StDev block for identification | Avg coverage for identification | StDev coverage for identification | Accuracy | Identification level (fraction of trials) | | | | Avg false positives | StDev false positives |
|---|---|---|---|---|---|---|---|---|---|---|---|---|---|
| | | | | | | | | Specific gene | Sub-class | Class | Incorrect | | |
| 1 | 4 | VANZA | 18.960 | 9.977 | 0.385 | 0.203 | 1.000 | 1.000 | 0.000 | 0.000 | 0.000 | 0.000 | 0.000 |
| 2 | 52 | VANWG | 15.600 | 7.444 | 0.183 | 0.087 | 1.000 | 0.400 | 0.600 | 0.000 | 0.000 | 0.000 | 0.000 |
| 3 | 62 | CTX | 19.080 | 14.003 | 0.217 | 0.158 | 1.000 | 0.000 | 1.000 | 0.000 | 0.000 | 0.000 | 0.000 |
| 4 | 68 | MEFA | 30.600 | 14.471 | 0.249 | 0.119 | 1.000 | 0.520 | 0.480 | 0.000 | 0.000 | 0.000 | 0.000 |
| 5 | 76 | OXA | 14.600 | 7.599 | 0.175 | 0.091 | 1.000 | 0.000 | 1.000 | 0.000 | 0.000 | 0.000 | 0.000 |
| 6 | 92 | CATB | 14.880 | 6.431 | 0.229 | 0.099 | 1.000 | 0.440 | 0.560 | 0.000 | 0.000 | 0.000 | 0.000 |
| 7 | 162 | EREA | 18.680 | 8.513 | 0.150 | 0.069 | 1.000 | 0.240 | 0.760 | 0.000 | 0.000 | 0.000 | 0.000 |
| 8 | 174 | TEM | 17.120 | 4.927 | 0.202 | 0.059 | 1.000 | 0.000 | 1.000 | 0.000 | 0.000 | 0.000 | 0.000 |
| 9 | 193 | CML | 17.560 | 9.070 | 0.133 | 0.068 | 1.000 | 0.240 | 0.760 | 0.000 | 0.000 | 0.000 | 0.000 |
| 10 | 207 | NIMA | 9.520 | 4.144 | 0.204 | 0.091 | 1.000 | 1.000 | 0.000 | 0.000 | 0.000 | 0.000 | 0.000 |
| 11 | 239 | DHFR | 12.200 | 5.824 | 0.252 | 0.119 | 1.000 | 0.160 | 0.840 | 0.000 | 0.000 | 0.000 | 0.000 |
| 12 | 246 | FOLP | 30.800 | 21.221 | 0.257 | 0.179 | 1.000 | 1.000 | 0.000 | 0.000 | 0.000 | 0.000 | 0.000 |
| 13 | 274 | PARC | 59.480 | 30.040 | 0.266 | 0.134 | 1.000 | 0.280 | 0.720 | 0.000 | 0.000 | 0.000 | 0.000 |
| 14 | 276 | SHV | 19.040 | 9.158 | 0.216 | 0.105 | 1.000 | 0.120 | 0.880 | 0.000 | 0.000 | 0.000 | 0.000 |
| 15 | 284 | DFRA | 14.960 | 5.504 | 0.265 | 0.099 | 1.000 | 1.000 | 0.000 | 0.000 | 0.000 | 0.000 | 0.000 |
| 16 | 295 | SOXS | 9.560 | 2.501 | 0.283 | 0.075 | 1.000 | 1.000 | 0.000 | 0.000 | 0.000 | 0.000 | 0.000 |
| 17 | 321 | TSNR | 23.560 | 15.202 | 0.296 | 0.192 | 1.000 | 1.000 | 0.000 | 0.000 | 0.000 | 0.000 | 0.000 |
| 18 | 413 | OMPD | 26.720 | 10.073 | 0.246 | 0.093 | 1.000 | 1.000 | 0.000 | 0.000 | 0.000 | 0.000 | 0.000 |
| 19 | 486 | LMRA | 9.320 | 3.648 | 0.160 | 0.062 | 1.000 | 1.000 | 0.000 | 0.000 | 0.000 | 0.000 | 0.000 |
| 20 | 505 | CARB | 21.760 | 8.171 | 0.238 | 0.089 | 1.000 | 1.000 | 0.000 | 0.000 | 0.000 | 0.000 | 0.000 |
| 21 | 506 | ANT3-DPRIME | 20.200 | 5.701 | 0.247 | 0.072 | 1.000 | 1.000 | 0.000 | 0.000 | 0.000 | 0.000 | 0.000 |
| 22 | 555 | QNRS | 23.160 | 6.656 | 0.347 | 0.101 | 1.000 | 0.400 | 0.600 | 0.000 | 0.000 | 0.000 | 0.000 |
| 23 | 558 | VANWI | 16.480 | 5.363 | 0.145 | 0.047 | 1.000 | 1.000 | 0.000 | 0.000 | 0.000 | 0.000 | 0.000 |
| 24 | 578 | TETX | 28.840 | 17.112 | 0.246 | 0.146 | 1.000 | 0.560 | 0.440 | 0.000 | 0.000 | 0.000 | 0.000 |
| 25 | 652 | VGA | 36.120 | 19.633 | 0.229 | 0.124 | 1.000 | 0.880 | 0.120 | 0.000 | 0.000 | 0.000 | 0.000 |
| 26 | 682 | MOX | 35.440 | 16.971 | 0.305 | 0.146 | 1.000 | 0.040 | 0.960 | 0.000 | 0.000 | 0.000 | 0.000 |
| 27 | 694 | ACT | 35.280 | 13.430 | 0.279 | 0.106 | 1.000 | 0.520 | 0.480 | 0.000 | 0.000 | 0.000 | 0.000 |
| 28 | 717 | VANRE | 23.240 | 9.212 | 0.329 | 0.131 | 1.000 | 1.000 | 0.000 | 0.000 | 0.000 | 0.000 | 0.000 |
| 29 | 749 | RMTB | 4.720 | 2.542 | 0.173 | 0.098 | 1.000 | 1.000 | 0.000 | 0.000 | 0.000 | 0.000 | 0.000 |
| 30 | 778 | DHA | 15.600 | 5.431 | 0.136 | 0.048 | 1.000 | 0.000 | 1.000 | 0.000 | 0.000 | 0.000 | 0.000 |
| 31 | 789 | CMY | 24.880 | 13.618 | 0.216 | 0.118 | 1.000 | 0.000 | 1.000 | 0.000 | 0.000 | 0.000 | 0.000 |
| 32 | 797 | FACT | 24.800 | 18.241 | 0.146 | 0.108 | 1.000 | 1.000 | 0.000 | 0.000 | 0.000 | 0.000 | 0.000 |
| 33 | 819 | OKP | 26.920 | 11.849 | 0.307 | 0.135 | 1.000 | 0.080 | 0.920 | 0.000 | 0.000 | 0.000 | 0.000 |
| 34 | 973 | IMP | 14.920 | 8.441 | 0.198 | 0.112 | 1.000 | 0.240 | 0.760 | 0.000 | 0.000 | 0.000 | 0.000 |
| 35 | 1048 | VIM | 12.320 | 4.964 | 0.149 | 0.060 | 1.000 | 0.200 | 0.800 | 0.000 | 0.000 | 0.000 | 0.000 |
| 36 | 1135 | PBP1B | 29.440 | 11.832 | 0.119 | 0.048 | 1.000 | 1.000 | 0.000 | 0.000 | 0.000 | 0.000 | 0.000 |
| 37 | 1146 | MPHE | 22.760 | 9.858 | 0.255 | 0.110 | 1.000 | 0.200 | 0.800 | 0.000 | 0.000 | 0.000 | 0.000 |
| 38 | 1182 | VATE | 16.240 | 6.139 | 0.248 | 0.093 | 1.000 | 0.640 | 0.360 | 0.000 | 0.000 | 0.000 | 0.000 |
| 39 | 1214 | OPRJ | 56.720 | 23.183 | 0.389 | 0.160 | 1.000 | 1.000 | 0.000 | 0.000 | 0.000 | 0.000 | 0.000 |
| 40 | 1254 | FOSB | 22.720 | 9.222 | 0.527 | 0.216 | 1.000 | 0.960 | 0.040 | 0.000 | 0.000 | 0.000 | 0.000 |
| 41 | 1271 | VPH | 17.040 | 10.382 | 0.196 | 0.119 | 1.000 | 1.000 | 0.000 | 0.000 | 0.000 | 0.000 | 0.000 |
| 42 | 1283 | SULI | 29.040 | 14.607 | 0.339 | 0.173 | 1.000 | 0.080 | 0.920 | 0.000 | 0.000 | 0.000 | 0.000 |
| 43 | 1297 | TET40 | 10.240 | 3.666 | 0.083 | 0.030 | 1.000 | 0.360 | 0.640 | 0.000 | 0.000 | 0.000 | 0.000 |
| 44 | 1389 | CPXAR | 17.680 | 3.891 | 0.248 | 0.052 | 1.000 | 0.680 | 0.320 | 0.000 | 0.000 | 0.000 | 0.000 |
| 45 | 1392 | AAC6-PRIME | 6.720 | 3.736 | 0.119 | 0.066 | 1.000 | 0.760 | 0.240 | 0.000 | 0.000 | 0.000 | 0.000 |
| 46 | 1422 | VGBB | 37.960 | 11.043 | 0.420 | 0.122 | 1.000 | 1.000 | 0.000 | 0.000 | 0.000 | 0.000 | 0.000 |
| 47 | 1440 | FOSC | 13.200 | 5.164 | 0.233 | 0.089 | 1.000 | 1.000 | 0.000 | 0.000 | 0.000 | 0.000 | 0.000 |

| | | | | | | | | | | | | | |
|---|---|---|---|---|---|---|---|---|---|---|---|---|---|
| 48 | 1535 | LNUA | 8.360 | 5.195 | 0.171 | 0.105 | 1.000 | 0.480 | 0.520 | 0.000 | 0.000 | 0.000 | 0.000 |
| 49 | 1569 | PARE | 38.560 | 15.565 | 0.202 | 0.082 | 1.000 | 0.920 | 0.080 | 0.000 | 0.000 | 0.000 | 0.000 |
| 50 | 1695 | NDM | 19.960 | 14.519 | 0.241 | 0.176 | 1.000 | 1.000 | 0.000 | 0.000 | 0.000 | 0.000 | 0.000 |
| 51 | 1702 | SPG | 17.680 | 10.213 | 0.205 | 0.118 | 1.000 | 1.000 | 0.000 | 0.000 | 0.000 | 0.000 | 0.000 |
| 52 | 1753 | VEB | 33.480 | 21.219 | 0.369 | 0.236 | 1.000 | 0.040 | 0.960 | 0.000 | 0.000 | 0.000 | 0.000 |
| 53 | 1953 | LNUB | 28.000 | 9.305 | 0.344 | 0.115 | 1.000 | 0.920 | 0.080 | 0.000 | 0.000 | 0.000 | 0.000 |
| 54 | 2026 | ERMA | 23.600 | 12.832 | 0.317 | 0.172 | 1.000 | 0.920 | 0.080 | 0.000 | 0.000 | 0.000 | 0.000 |
| 55 | 2357 | SULII | 16.120 | 7.918 | 0.196 | 0.096 | 1.000 | 0.080 | 0.920 | 0.000 | 0.000 | 0.000 | 0.000 |
| 56 | 2517 | TET37 | 11.640 | 5.057 | 0.348 | 0.155 | 1.000 | 1.000 | 0.000 | 0.000 | 0.000 | 0.000 | 0.000 |
| 57 | 2822 | EMRK | 53.120 | 11.591 | 0.499 | 0.110 | 1.000 | 1.000 | 0.000 | 0.000 | 0.000 | 0.000 | 0.000 |
| 58 | 2999 | MPHB | 37.360 | 13.853 | 0.407 | 0.153 | 1.000 | 1.000 | 0.000 | 0.000 | 0.000 | 0.000 | 0.000 |
| 59 | 3024 | VANYM | 24.200 | 8.005 | 0.338 | 0.113 | 1.000 | 1.000 | 0.000 | 0.000 | 0.000 | 0.000 | 0.000 |
| 60 | 3041 | MECC | 75.400 | 59.431 | 0.374 | 0.295 | 1.000 | 1.000 | 0.000 | 0.000 | 0.000 | 0.000 | 0.000 |
| 61 | 3128 | TUFAB | 22.280 | 9.689 | 0.186 | 0.081 | 1.000 | 0.840 | 0.160 | 0.000 | 0.000 | 0.000 | 0.000 |
| 62 | 3176 | AMRB | 57.400 | 25.120 | 0.182 | 0.079 | 1.000 | 1.000 | 0.000 | 0.000 | 0.000 | 0.000 | 0.000 |
| 63 | 3270 | IRI | 79.120 | 12.640 | 0.543 | 0.087 | 1.000 | 1.000 | 0.000 | 0.000 | 0.000 | 0.000 | 0.000 |
| 64 | 3314 | RPOB | 53.360 | 17.411 | 0.149 | 0.049 | 1.000 | 0.000 | 1.000 | 0.000 | 0.000 | 0.000 | 0.000 |
| 65 | 3332 | TET35 | 20.240 | 10.948 | 0.180 | 0.096 | 1.000 | 1.000 | 0.000 | 0.000 | 0.000 | 0.000 | 0.000 |
| 66 | 3370 | CFRA | 34.840 | 15.407 | 0.327 | 0.146 | 1.000 | 1.000 | 0.000 | 0.000 | 0.000 | 0.000 | 0.000 |
| 67 | 3513 | BRP | 8.720 | 3.398 | 0.213 | 0.083 | 1.000 | 1.000 | 0.000 | 0.000 | 0.000 | 0.000 | 0.000 |
| 68 | 3613 | APH3-PRIME | 30.560 | 16.000 | 0.375 | 0.195 | 1.000 | 1.000 | 0.000 | 0.000 | 0.000 | 0.000 | 0.000 |
| 69 | 3697 | TETM | 46.840 | 19.433 | 0.243 | 0.100 | 1.000 | 0.560 | 0.440 | 0.000 | 0.000 | 0.000 | 0.000 |
| 70 | 3778 | IND | 17.000 | 5.715 | 0.229 | 0.075 | 1.000 | 0.160 | 0.840 | 0.000 | 0.000 | 0.000 | 0.000 |

Simulation settings:

k-mers: ‘constant’, k=10
Gene coverage: 1
Number of genes: 1
Errors: ‘off’
Penalty score: 0.1
Thresholding: multiplier=1, all factors (1-on)
Entropy screening: ‘rand’

**Table S4.** Resistance genes simulations with 8-mer blocks

| No. | Database No. | Gene sub-class | Avg block for identification | StDev block for identification | Avg coverage for identification | StDev coverage for identification | Accuracy | Identification level (fraction of trials) | | | | Avg false positives | StDev false positives |
|---|---|---|---|---|---|---|---|---|---|---|---|---|---|
| | | | | | | | | Specific gene | Sub-class | Class | Incorrect | | |
| 1 | 4 | VANZA | 20.360 | 10.327 | 0.331 | 0.170 | 1.000 | 1.000 | 0.000 | 0.000 | 0.000 | 0.000 | 0.000 |
| 2 | 52 | VANWG | 37.800 | 17.963 | 0.356 | 0.170 | 1.000 | 0.440 | 0.560 | 0.000 | 0.000 | 0.000 | 0.000 |
| 3 | 62 | CTX | 37.280 | 13.252 | 0.339 | 0.120 | 1.000 | 0.000 | 1.000 | 0.000 | 0.000 | 0.000 | 0.000 |
| 4 | 68 | MEFA | 43.960 | 19.385 | 0.288 | 0.127 | 1.000 | 0.520 | 0.480 | 0.000 | 0.000 | 0.000 | 0.000 |
| 5 | 76 | OXA | 35.640 | 18.259 | 0.343 | 0.176 | 1.000 | 0.000 | 1.000 | 0.000 | 0.000 | 0.000 | 0.000 |
| 6 | 92 | CATB | 22.160 | 9.711 | 0.278 | 0.122 | 1.000 | 0.320 | 0.680 | 0.000 | 0.000 | 0.000 | 0.000 |
| 7 | 162 | EREA | 38.720 | 17.387 | 0.251 | 0.113 | 1.000 | 0.120 | 0.880 | 0.000 | 0.000 | 0.000 | 0.000 |
| 8 | 174 | TEM | 37.640 | 13.778 | 0.359 | 0.131 | 1.000 | 0.000 | 1.000 | 0.000 | 0.000 | 0.000 | 0.000 |
| 9 | 193 | CML | 26.160 | 15.032 | 0.159 | 0.091 | 1.000 | 0.000 | 1.000 | 0.000 | 0.000 | 0.000 | 0.000 |
| 10 | 207 | NIMA | 13.800 | 5.951 | 0.236 | 0.102 | 1.000 | 0.960 | 0.040 | 0.000 | 0.000 | 0.000 | 0.000 |
| 11 | 239 | DHFR | 23.680 | 9.040 | 0.393 | 0.151 | 1.000 | 0.320 | 0.680 | 0.000 | 0.000 | 0.000 | 0.000 |
| 12 | 246 | FOLP | 40.833 | 19.699 | 0.274 | 0.132 | 0.960 | 0.920 | 0.040 | 0.000 | 0.040 | 0.000 | 0.000 |
| 13 | 274 | PARC | 72.720 | 31.383 | 0.262 | 0.113 | 1.000 | 0.280 | 0.720 | 0.000 | 0.000 | 0.000 | 0.000 |
| 14 | 276 | SHV | 32.400 | 14.939 | 0.297 | 0.137 | 1.000 | 0.000 | 1.000 | 0.000 | 0.000 | 0.000 | 0.000 |
| 15 | 284 | DFRA | 29.080 | 9.639 | 0.416 | 0.137 | 1.000 | 1.000 | 0.000 | 0.000 | 0.000 | 0.000 | 0.000 |
| 16 | 295 | SOXS | 18.320 | 8.764 | 0.446 | 0.215 | 1.000 | 1.000 | 0.000 | 0.000 | 0.000 | 0.000 | 0.000 |
| 17 | 321 | TSNR | 38.636 | 20.254 | 0.391 | 0.206 | 0.880 | 0.880 | 0.000 | 0.000 | 0.120 | 0.000 | 0.000 |
| 18 | 413 | OMPD | 37.560 | 12.203 | 0.279 | 0.090 | 1.000 | 1.000 | 0.000 | 0.000 | 0.000 | 0.000 | 0.000 |
| 19 | 486 | LMRA | 16.800 | 7.130 | 0.234 | 0.098 | 1.000 | 1.000 | 0.000 | 0.000 | 0.000 | 0.000 | 0.000 |
| 20 | 505 | CARB | 31.200 | 11.576 | 0.276 | 0.104 | 1.000 | 1.000 | 0.000 | 0.000 | 0.000 | 0.000 | 0.000 |
| 21 | 506 | ANT3-DPRIME | 27.480 | 9.896 | 0.275 | 0.099 | 1.000 | 0.960 | 0.040 | 0.000 | 0.000 | 0.000 | 0.000 |
| 22 | 555 | QNRS | 28.680 | 11.821 | 0.345 | 0.143 | 1.000 | 0.360 | 0.640 | 0.000 | 0.000 | 0.000 | 0.000 |
| 23 | 558 | VANWI | 32.280 | 15.038 | 0.229 | 0.107 | 1.000 | 1.000 | 0.000 | 0.000 | 0.000 | 0.000 | 0.000 |
| 24 | 578 | TETX | 48.560 | 14.463 | 0.332 | 0.099 | 1.000 | 0.240 | 0.760 | 0.000 | 0.000 | 0.000 | 0.000 |
| 25 | 652 | VGA | 61.739 | 32.433 | 0.312 | 0.164 | 0.920 | 0.840 | 0.080 | 0.000 | 0.080 | 0.000 | 0.000 |
| 26 | 682 | MOX | 55.400 | 20.145 | 0.384 | 0.140 | 1.000 | 0.320 | 0.680 | 0.000 | 0.000 | 0.000 | 0.000 |
| 27 | 694 | ACT | 43.320 | 22.090 | 0.277 | 0.141 | 1.000 | 0.080 | 0.920 | 0.000 | 0.000 | 0.000 | 0.000 |
| 28 | 717 | VANRE | 31.400 | 17.772 | 0.361 | 0.205 | 1.000 | 1.000 | 0.000 | 0.000 | 0.000 | 0.000 | 0.000 |
| 29 | 749 | RMTB | 11.040 | 5.111 | 0.344 | 0.161 | 1.000 | 1.000 | 0.000 | 0.000 | 0.000 | 0.000 | 0.000 |
| 30 | 778 | DHA | 41.920 | 20.004 | 0.293 | 0.140 | 1.000 | 0.040 | 0.960 | 0.000 | 0.000 | 0.000 | 0.000 |
| 31 | 789 | CMY | 39.640 | 16.153 | 0.275 | 0.112 | 1.000 | 0.000 | 1.000 | 0.000 | 0.000 | 0.000 | 0.000 |
| 32 | 797 | FACT | 40.167 | 25.862 | 0.190 | 0.122 | 0.960 | 0.960 | 0.000 | 0.000 | 0.040 | 0.000 | 0.000 |
| 33 | 819 | OKP | 45.760 | 22.244 | 0.424 | 0.206 | 1.000 | 0.040 | 0.960 | 0.000 | 0.000 | 0.000 | 0.000 |
| 34 | 973 | IMP | 31.520 | 12.210 | 0.339 | 0.131 | 1.000 | 0.040 | 0.960 | 0.000 | 0.000 | 0.000 | 0.000 |
| 35 | 1048 | VIM | 27.920 | 14.250 | 0.277 | 0.142 | 1.000 | 0.080 | 0.920 | 0.000 | 0.000 | 0.000 | 0.000 |
| 36 | 1135 | PBP1B | 46.520 | 22.006 | 0.151 | 0.071 | 1.000 | 1.000 | 0.000 | 0.000 | 0.000 | 0.000 | 0.000 |
| 37 | 1146 | MPHE | 43.720 | 17.119 | 0.394 | 0.154 | 1.000 | 0.280 | 0.720 | 0.000 | 0.000 | 0.000 | 0.000 |
| 38 | 1182 | VATE | 20.800 | 8.818 | 0.257 | 0.109 | 1.000 | 0.200 | 0.800 | 0.000 | 0.000 | 0.000 | 0.000 |
| 39 | 1214 | OPRJ | 69.320 | 31.201 | 0.383 | 0.172 | 1.000 | 1.000 | 0.000 | 0.000 | 0.000 | 0.000 | 0.000 |
| 40 | 1254 | FOSB | 17.880 | 7.839 | 0.337 | 0.148 | 1.000 | 0.920 | 0.080 | 0.000 | 0.000 | 0.000 | 0.000 |
| 41 | 1271 | VPH | 30.000 | 16.427 | 0.275 | 0.150 | 1.000 | 1.000 | 0.000 | 0.000 | 0.000 | 0.000 | 0.000 |
| 42 | 1283 | SULI | 45.080 | 17.949 | 0.426 | 0.170 | 1.000 | 0.280 | 0.720 | 0.000 | 0.000 | 0.000 | 0.000 |
| 43 | 1297 | TET40 | 34.080 | 18.841 | 0.223 | 0.123 | 1.000 | 0.560 | 0.440 | 0.000 | 0.000 | 0.000 | 0.000 |
| 44 | 1389 | CPXAR | 27.000 | 8.211 | 0.306 | 0.093 | 1.000 | 0.360 | 0.640 | 0.000 | 0.000 | 0.000 | 0.000 |
| 45 | 1392 | AAC6-PRIME | 17.160 | 9.168 | 0.245 | 0.130 | 1.000 | 0.640 | 0.360 | 0.000 | 0.000 | 0.000 | 0.000 |
| 46 | 1422 | VGBB | 39.560 | 16.153 | 0.354 | 0.144 | 1.000 | 1.000 | 0.000 | 0.000 | 0.000 | 0.000 | 0.000 |
| 47 | 1440 | FOSC | 17.360 | 9.691 | 0.252 | 0.140 | 1.000 | 1.000 | 0.000 | 0.000 | 0.000 | 0.000 | 0.000 |

| | | | | | | | | | | | | | |
|---|---|---|---|---|---|---|---|---|---|---|---|---|---|
| 48 | 1535 | LNUA | 15.280 | 8.975 | 0.250 | 0.148 | 1.000 | 0.400 | 0.600 | 0.000 | 0.000 | 0.000 | 0.000 |
| 49 | 1569 | PARE | 72.800 | 23.189 | 0.308 | 0.098 | 1.000 | 0.320 | 0.680 | 0.000 | 0.000 | 0.000 | 0.000 |
| 50 | 1695 | NDM | 25.280 | 9.410 | 0.248 | 0.092 | 1.000 | 1.000 | 0.000 | 0.000 | 0.000 | 0.000 | 0.000 |
| 51 | 1702 | SPG | 29.480 | 13.574 | 0.273 | 0.125 | 1.000 | 1.000 | 0.000 | 0.000 | 0.000 | 0.000 | 0.000 |
| 52 | 1753 | VEB | 33.280 | 15.568 | 0.295 | 0.138 | 1.000 | 0.000 | 1.000 | 0.000 | 0.000 | 0.000 | 0.000 |
| 53 | 1953 | LNUB | 36.480 | 13.226 | 0.362 | 0.131 | 1.000 | 0.560 | 0.440 | 0.000 | 0.000 | 0.000 | 0.000 |
| 54 | 2026 | ERMA | 31.000 | 14.018 | 0.337 | 0.152 | 1.000 | 0.720 | 0.280 | 0.000 | 0.000 | 0.000 | 0.000 |
| 55 | 2357 | SULII | 30.417 | 13.445 | 0.296 | 0.131 | 0.960 | 0.000 | 0.960 | 0.000 | 0.040 | 0.000 | 0.000 |
| 56 | 2517 | TET37 | 17.440 | 6.868 | 0.415 | 0.165 | 1.000 | 1.000 | 0.000 | 0.000 | 0.000 | 0.000 | 0.000 |
| 57 | 2822 | EMRK | 41.160 | 19.686 | 0.310 | 0.148 | 1.000 | 1.000 | 0.000 | 0.000 | 0.000 | 0.000 | 0.000 |
| 58 | 2999 | MPHB | 29.667 | 14.577 | 0.260 | 0.128 | 0.960 | 0.960 | 0.000 | 0.000 | 0.040 | 0.000 | 0.000 |
| 59 | 3024 | VANYM | 46.680 | 19.991 | 0.528 | 0.224 | 1.000 | 0.960 | 0.040 | 0.000 | 0.000 | 0.160 | 0.374 |
| 60 | 3041 | MECC | 214.667 | 68.542 | 0.853 | 0.272 | 0.960 | 0.280 | 0.680 | 0.000 | 0.040 | 0.000 | 0.000 |
| 61 | 3128 | TUFAB | 33.760 | 14.042 | 0.226 | 0.095 | 1.000 | 0.400 | 0.600 | 0.000 | 0.000 | 0.000 | 0.000 |
| 62 | 3176 | AMRB | 97.320 | 39.918 | 0.248 | 0.102 | 1.000 | 1.000 | 0.000 | 0.000 | 0.000 | 0.000 | 0.000 |
| 63 | 3270 | IRI | 62.125 | 22.462 | 0.344 | 0.125 | 0.960 | 0.960 | 0.000 | 0.000 | 0.040 | 0.000 | 0.000 |
| 64 | 3314 | RPOB | 89.320 | 57.045 | 0.201 | 0.128 | 1.000 | 0.160 | 0.840 | 0.000 | 0.000 | 0.000 | 0.000 |
| 65 | 3332 | TET35 | 22.240 | 11.598 | 0.159 | 0.083 | 1.000 | 1.000 | 0.000 | 0.000 | 0.000 | 0.000 | 0.000 |
| 66 | 3370 | CFRA | 70.960 | 19.659 | 0.537 | 0.149 | 1.000 | 1.000 | 0.000 | 0.000 | 0.000 | 0.000 | 0.000 |
| 67 | 3513 | BRP | 12.320 | 6.549 | 0.242 | 0.127 | 1.000 | 1.000 | 0.000 | 0.000 | 0.000 | 0.000 | 0.000 |
| 68 | 3613 | APH3-PRIME | 40.120 | 17.050 | 0.397 | 0.169 | 1.000 | 1.000 | 0.000 | 0.000 | 0.000 | 0.000 | 0.000 |
| 69 | 3697 | TETM | 77.200 | 37.236 | 0.320 | 0.155 | 1.000 | 0.400 | 0.600 | 0.000 | 0.000 | 0.000 | 0.000 |
| 70 | 3778 | IND | 36.920 | 12.124 | 0.402 | 0.132 | 1.000 | 0.120 | 0.880 | 0.000 | 0.000 | 0.000 | 0.000 |

Simulation settings:

k-mers: ‘constant’, k=8
Gene coverage: 1
Number of genes: 1
Errors: ‘off’
Penalty score: 0.1
Thresholding: multiplier=1, all factors (1-on)
Entropy screening: ‘rand’

**Table S5.** Resistance genes simulations with 12-mer blocks

| No. | Database No. | Gene sub-class | Avg block for identification | StDev block for identification | Avg coverage for identification | StDev coverage for identification | Accuracy | Identification level (fraction of trials) | | | | Avg false positives | StDev false positives |
|---|---|---|---|---|---|---|---|---|---|---|---|---|---|
| | | | | | | | | Specific gene | Sub-class | Class | Incorrect | | |
| 1 | 4 | VANZA | 13.080 | 3.818 | 0.312 | 0.094 | 1.000 | 1.000 | 0.000 | 0.000 | 0.000 | 0.000 | 0.000 |
| 2 | 52 | VANWG | 8.440 | 2.987 | 0.115 | 0.042 | 1.000 | 0.720 | 0.280 | 0.000 | 0.000 | 0.000 | 0.000 |
| 3 | 62 | CTX | 11.640 | 6.915 | 0.150 | 0.094 | 1.000 | 0.000 | 1.000 | 0.000 | 0.000 | 0.000 | 0.000 |
| 4 | 68 | MEFA | 12.640 | 4.212 | 0.121 | 0.041 | 1.000 | 0.480 | 0.520 | 0.000 | 0.000 | 0.000 | 0.000 |
| 5 | 76 | OXA | 7.200 | 4.203 | 0.098 | 0.058 | 1.000 | 0.000 | 1.000 | 0.000 | 0.000 | 0.000 | 0.000 |
| 6 | 92 | CATB | 6.520 | 2.756 | 0.113 | 0.050 | 1.000 | 0.080 | 0.920 | 0.000 | 0.000 | 0.000 | 0.000 |
| 7 | 162 | EREA | 5.200 | 2.021 | 0.046 | 0.019 | 1.000 | 0.440 | 0.560 | 0.000 | 0.000 | 0.000 | 0.000 |
| 8 | 174 | TEM | 11.640 | 6.415 | 0.159 | 0.092 | 1.000 | 0.000 | 1.000 | 0.000 | 0.000 | 0.000 | 0.000 |
| 9 | 193 | CML | 5.520 | 4.360 | 0.047 | 0.039 | 1.000 | 0.000 | 1.000 | 0.000 | 0.000 | 0.000 | 0.000 |
| 10 | 207 | NIMA | 8.600 | 5.845 | 0.203 | 0.153 | 1.000 | 1.000 | 0.000 | 0.000 | 0.000 | 0.000 | 0.000 |
| 11 | 239 | DHFR | 10.800 | 4.330 | 0.262 | 0.109 | 1.000 | 0.280 | 0.720 | 0.000 | 0.000 | 0.000 | 0.000 |
| 12 | 246 | FOLP | 14.160 | 5.843 | 0.137 | 0.056 | 1.000 | 1.000 | 0.000 | 0.000 | 0.000 | 0.000 | 0.000 |
| 13 | 274 | PARC | 20.400 | 7.539 | 0.106 | 0.039 | 1.000 | 0.560 | 0.440 | 0.000 | 0.000 | 0.000 | 0.000 |
| 14 | 276 | SHV | 11.800 | 9.000 | 0.157 | 0.123 | 1.000 | 0.000 | 1.000 | 0.000 | 0.000 | 0.000 | 0.000 |
| 15 | 284 | DFRA | 11.240 | 5.585 | 0.225 | 0.120 | 1.000 | 1.000 | 0.000 | 0.000 | 0.000 | 0.000 | 0.000 |
| 16 | 295 | SOXS | 10.120 | 5.418 | 0.340 | 0.200 | 1.000 | 1.000 | 0.000 | 0.000 | 0.000 | 0.000 | 0.000 |
| 17 | 321 | TSNR | 10.560 | 6.678 | 0.154 | 0.101 | 1.000 | 1.000 | 0.000 | 0.000 | 0.000 | 0.000 | 0.000 |
| 18 | 413 | OMPD | 10.640 | 4.202 | 0.114 | 0.047 | 1.000 | 1.000 | 0.000 | 0.000 | 0.000 | 0.000 | 0.000 |
| 19 | 486 | LMRA | 9.000 | 3.651 | 0.176 | 0.077 | 1.000 | 1.000 | 0.000 | 0.000 | 0.000 | 0.000 | 0.000 |
| 20 | 505 | CARB | 10.040 | 5.827 | 0.124 | 0.078 | 1.000 | 1.000 | 0.000 | 0.000 | 0.000 | 0.000 | 0.000 |
| 21 | 506 | ANT3-DPRIME | 10.880 | 5.126 | 0.151 | 0.078 | 1.000 | 0.920 | 0.080 | 0.000 | 0.000 | 0.000 | 0.000 |
| 22 | 555 | QNRS | 13.200 | 6.770 | 0.228 | 0.117 | 1.000 | 0.320 | 0.680 | 0.000 | 0.000 | 0.000 | 0.000 |
| 23 | 558 | VANWI | 11.680 | 4.190 | 0.120 | 0.045 | 1.000 | 1.000 | 0.000 | 0.000 | 0.000 | 0.000 | 0.000 |
| 24 | 578 | TETX | 16.240 | 7.535 | 0.159 | 0.077 | 1.000 | 0.280 | 0.720 | 0.000 | 0.000 | 0.000 | 0.000 |
| 25 | 652 | VGA | 15.320 | 6.067 | 0.113 | 0.045 | 1.000 | 0.880 | 0.120 | 0.000 | 0.000 | 0.000 | 0.000 |
| 26 | 682 | MOX | 18.720 | 9.280 | 0.190 | 0.097 | 1.000 | 0.200 | 0.800 | 0.000 | 0.000 | 0.000 | 0.000 |
| 27 | 694 | ACT | 22.520 | 10.798 | 0.212 | 0.104 | 1.000 | 0.080 | 0.920 | 0.000 | 0.000 | 0.000 | 0.000 |
| 28 | 717 | VANRE | 10.600 | 4.311 | 0.178 | 0.074 | 1.000 | 1.000 | 0.000 | 0.000 | 0.000 | 0.000 | 0.000 |
| 29 | 749 | RMTB | 3.520 | 3.607 | 0.140 | 0.168 | 1.000 | 1.000 | 0.000 | 0.000 | 0.000 | 0.000 | 0.000 |
| 30 | 778 | DHA | 9.320 | 6.939 | 0.091 | 0.072 | 1.000 | 0.080 | 0.920 | 0.000 | 0.000 | 0.000 | 0.000 |
| 31 | 789 | CMY | 9.360 | 8.093 | 0.095 | 0.084 | 1.000 | 0.000 | 1.000 | 0.000 | 0.000 | 0.000 | 0.000 |
| 32 | 797 | FACT | 20.640 | 9.780 | 0.141 | 0.070 | 1.000 | 1.000 | 0.000 | 0.000 | 0.000 | 0.000 | 0.000 |
| 33 | 819 | OKP | 6.640 | 3.872 | 0.088 | 0.050 | 1.000 | 0.080 | 0.920 | 0.000 | 0.000 | 0.000 | 0.000 |
| 34 | 973 | IMP | 8.360 | 3.774 | 0.126 | 0.055 | 1.000 | 0.240 | 0.760 | 0.000 | 0.000 | 0.000 | 0.000 |
| 35 | 1048 | VIM | 4.160 | 1.491 | 0.058 | 0.020 | 1.000 | 0.080 | 0.920 | 0.000 | 0.000 | 0.000 | 0.000 |
| 36 | 1135 | PBP1B | 6.160 | 1.795 | 0.029 | 0.008 | 1.000 | 1.000 | 0.000 | 0.000 | 0.000 | 0.000 | 0.000 |
| 37 | 1146 | MPHE | 16.600 | 3.559 | 0.216 | 0.050 | 1.000 | 0.240 | 0.760 | 0.000 | 0.000 | 0.000 | 0.000 |
| 38 | 1182 | VATE | 9.200 | 6.007 | 0.157 | 0.104 | 1.000 | 0.680 | 0.320 | 0.000 | 0.000 | 0.000 | 0.000 |
| 39 | 1214 | OPRJ | 21.040 | 9.948 | 0.169 | 0.083 | 1.000 | 1.000 | 0.000 | 0.000 | 0.000 | 0.000 | 0.000 |
| 40 | 1254 | FOSB | 13.560 | 4.292 | 0.366 | 0.122 | 1.000 | 1.000 | 0.000 | 0.000 | 0.000 | 0.000 | 0.000 |
| 41 | 1271 | VPH | 10.760 | 4.447 | 0.138 | 0.060 | 1.000 | 1.000 | 0.000 | 0.000 | 0.000 | 0.000 | 0.000 |
| 42 | 1283 | SULI | 19.520 | 7.779 | 0.268 | 0.111 | 1.000 | 0.280 | 0.720 | 0.000 | 0.000 | 0.000 | 0.000 |
| 43 | 1297 | TET40 | 4.200 | 1.581 | 0.039 | 0.014 | 1.000 | 0.840 | 0.160 | 0.000 | 0.000 | 0.000 | 0.000 |
| 44 | 1389 | CPXAR | 10.840 | 4.469 | 0.173 | 0.077 | 1.000 | 0.320 | 0.680 | 0.000 | 0.000 | 0.000 | 0.000 |
| 45 | 1392 | AAC6-PRIME | 3.160 | 1.993 | 0.062 | 0.038 | 1.000 | 0.720 | 0.280 | 0.000 | 0.000 | 0.000 | 0.000 |
| 46 | 1422 | VGBB | 15.640 | 6.291 | 0.199 | 0.085 | 1.000 | 1.000 | 0.000 | 0.000 | 0.000 | 0.000 | 0.000 |
| 47 | 1440 | FOSC | 8.200 | 5.649 | 0.166 | 0.119 | 1.000 | 1.000 | 0.000 | 0.000 | 0.000 | 0.000 | 0.000 |

| | | | | | | | | | | | | | |
|---|---|---|---|---|---|---|---|---|---|---|---|---|---|
| 48 | 1535 | LNUA | 4.040 | 2.091 | 0.094 | 0.048 | 1.000 | 0.440 | 0.560 | 0.000 | 0.000 | 0.000 | 0.000 |
| 49 | 1569 | PARE | 15.240 | 8.997 | 0.095 | 0.057 | 1.000 | 0.800 | 0.200 | 0.000 | 0.000 | 0.000 | 0.000 |
| 50 | 1695 | NDM | 9.680 | 5.313 | 0.133 | 0.072 | 1.000 | 1.000 | 0.000 | 0.000 | 0.000 | 0.000 | 0.000 |
| 51 | 1702 | SPG | 9.040 | 9.889 | 0.122 | 0.137 | 1.000 | 1.000 | 0.000 | 0.000 | 0.000 | 0.000 | 0.000 |
| 52 | 1753 | VEB | 12.520 | 5.221 | 0.161 | 0.066 | 1.000 | 0.080 | 0.920 | 0.000 | 0.000 | 0.000 | 0.000 |
| 53 | 1953 | LNUB | 15.280 | 5.136 | 0.218 | 0.077 | 1.000 | 0.400 | 0.600 | 0.000 | 0.000 | 0.000 | 0.000 |
| 54 | 2026 | ERMA | 8.240 | 4.503 | 0.127 | 0.071 | 1.000 | 0.920 | 0.080 | 0.000 | 0.000 | 0.000 | 0.000 |
| 55 | 2357 | SULII | 9.920 | 7.371 | 0.135 | 0.105 | 1.000 | 0.040 | 0.960 | 0.000 | 0.000 | 0.000 | 0.000 |
| 56 | 2517 | TET37 | 7.600 | 3.488 | 0.255 | 0.127 | 1.000 | 1.000 | 0.000 | 0.000 | 0.000 | 0.000 | 0.000 |
| 57 | 2822 | EMRK | 17.600 | 5.583 | 0.189 | 0.063 | 1.000 | 1.000 | 0.000 | 0.000 | 0.000 | 0.000 | 0.000 |
| 58 | 2999 | MPHB | 11.920 | 6.006 | 0.150 | 0.077 | 1.000 | 1.000 | 0.000 | 0.000 | 0.000 | 0.000 | 0.000 |
| 59 | 3024 | VANYM | 12.840 | 3.508 | 0.212 | 0.060 | 1.000 | 1.000 | 0.000 | 0.000 | 0.000 | 0.000 | 0.000 |
| 60 | 3041 | MECC | 124.360 | 52.273 | 0.739 | 0.312 | 1.000 | 0.520 | 0.480 | 0.000 | 0.000 | 0.000 | 0.000 |
| 61 | 3128 | TUFAB | 12.080 | 8.103 | 0.118 | 0.083 | 1.000 | 0.640 | 0.360 | 0.000 | 0.000 | 0.000 | 0.000 |
| 62 | 3176 | AMRB | 25.400 | 14.751 | 0.096 | 0.056 | 1.000 | 1.000 | 0.000 | 0.000 | 0.000 | 0.000 | 0.000 |
| 63 | 3270 | IRI | 11.320 | 5.865 | 0.088 | 0.048 | 1.000 | 1.000 | 0.000 | 0.000 | 0.000 | 0.000 | 0.000 |
| 64 | 3314 | RPOB | 20.960 | 21.784 | 0.069 | 0.074 | 1.000 | 0.320 | 0.680 | 0.000 | 0.000 | 0.000 | 0.000 |
| 65 | 3332 | TET35 | 8.080 | 4.907 | 0.083 | 0.053 | 1.000 | 1.000 | 0.000 | 0.000 | 0.000 | 0.000 | 0.000 |
| 66 | 3370 | CFRA | 19.320 | 5.692 | 0.215 | 0.065 | 1.000 | 1.000 | 0.000 | 0.000 | 0.000 | 0.000 | 0.000 |
| 67 | 3513 | BRP | 9.040 | 2.865 | 0.251 | 0.086 | 1.000 | 1.000 | 0.000 | 0.000 | 0.000 | 0.000 | 0.000 |
| 68 | 3613 | APH3-PRIME | 15.760 | 5.101 | 0.227 | 0.074 | 1.000 | 1.000 | 0.000 | 0.000 | 0.000 | 0.000 | 0.000 |
| 69 | 3697 | TETM | 16.320 | 10.590 | 0.098 | 0.065 | 1.000 | 0.520 | 0.480 | 0.000 | 0.000 | 0.000 | 0.000 |
| 70 | 3778 | IND | 11.080 | 3.639 | 0.170 | 0.057 | 1.000 | 0.040 | 0.960 | 0.000 | 0.000 | 0.000 | 0.000 |

Simulation settings:

k-mers: ‘constant’, k=12
Gene coverage: 1
Number of genes: 1
Errors: ‘off’
Penalty score: 0.1
Thresholding: multiplier=1, all factors (1-on)
Entropy screening: ‘rand’

**Table S6.** Resistance genes simulations with variable-mer blocks centered around k=10

| No. | Database No. | Gene sub-class | Avg block for identification | StDev block for identification | Avg coverage for identification | StDev coverage for identification | Accuracy | Identification level (fraction of trials) | | | | Avg false positives | StDev false positives |
|---|---|---|---|---|---|---|---|---|---|---|---|---|---|
| | | | | | | | | Specific gene | Sub-class | Class | Incorrect | | |
| 1 | 4 | VANZA | 19.120 | 8.268 | 0.350 | 0.158 | 1.000 | 1.000 | 0.000 | 0.000 | 0.000 | 0.000 | 0.000 |
| 2 | 52 | VANWG | 21.040 | 12.408 | 0.224 | 0.131 | 1.000 | 0.120 | 0.880 | 0.000 | 0.000 | 0.000 | 0.000 |
| 3 | 62 | CTX | 20.800 | 10.178 | 0.206 | 0.098 | 1.000 | 0.000 | 1.000 | 0.000 | 0.000 | 0.000 | 0.000 |
| 4 | 68 | MEFA | 31.960 | 15.624 | 0.231 | 0.114 | 1.000 | 0.560 | 0.440 | 0.000 | 0.000 | 0.000 | 0.000 |
| 5 | 76 | OXA | 21.440 | 10.587 | 0.228 | 0.111 | 1.000 | 0.000 | 1.000 | 0.000 | 0.000 | 0.000 | 0.000 |
| 6 | 92 | CATB | 15.320 | 6.460 | 0.207 | 0.085 | 1.000 | 0.120 | 0.880 | 0.000 | 0.000 | 0.000 | 0.000 |
| 7 | 162 | EREA | 22.240 | 11.530 | 0.157 | 0.080 | 1.000 | 0.360 | 0.640 | 0.000 | 0.000 | 0.000 | 0.000 |
| 8 | 174 | TEM | 21.280 | 7.591 | 0.221 | 0.079 | 1.000 | 0.000 | 1.000 | 0.000 | 0.000 | 0.000 | 0.000 |
| 9 | 193 | CML | 18.840 | 9.150 | 0.124 | 0.060 | 1.000 | 0.200 | 0.800 | 0.000 | 0.000 | 0.000 | 0.000 |
| 10 | 207 | NIMA | 10.560 | 4.360 | 0.195 | 0.082 | 1.000 | 0.960 | 0.040 | 0.000 | 0.000 | 0.000 | 0.000 |
| 11 | 239 | DHFR | 12.400 | 4.882 | 0.227 | 0.090 | 1.000 | 0.320 | 0.680 | 0.000 | 0.000 | 0.000 | 0.000 |
| 12 | 246 | FOLP | 31.960 | 21.458 | 0.246 | 0.165 | 1.000 | 0.920 | 0.080 | 0.000 | 0.000 | 0.000 | 0.000 |
| 13 | 274 | PARC | 49.040 | 20.733 | 0.191 | 0.080 | 1.000 | 0.200 | 0.800 | 0.000 | 0.000 | 0.000 | 0.000 |
| 14 | 276 | SHV | 22.560 | 9.933 | 0.228 | 0.105 | 1.000 | 0.080 | 0.920 | 0.000 | 0.000 | 0.000 | 0.000 |
| 15 | 284 | DFRA | 19.280 | 8.872 | 0.304 | 0.153 | 1.000 | 1.000 | 0.000 | 0.000 | 0.000 | 0.000 | 0.000 |
| 16 | 295 | SOXS | 13.200 | 4.770 | 0.353 | 0.134 | 1.000 | 1.000 | 0.000 | 0.000 | 0.000 | 0.000 | 0.000 |
| 17 | 321 | TSNR | 21.120 | 10.113 | 0.237 | 0.114 | 1.000 | 1.000 | 0.000 | 0.000 | 0.000 | 0.000 | 0.000 |
| 18 | 413 | OMPD | 23.360 | 9.827 | 0.189 | 0.078 | 1.000 | 1.000 | 0.000 | 0.000 | 0.000 | 0.000 | 0.000 |
| 19 | 486 | LMRA | 11.760 | 5.904 | 0.177 | 0.085 | 1.000 | 1.000 | 0.000 | 0.000 | 0.000 | 0.000 | 0.000 |
| 20 | 505 | CARB | 20.040 | 10.470 | 0.193 | 0.098 | 1.000 | 1.000 | 0.000 | 0.000 | 0.000 | 0.000 | 0.000 |
| 21 | 506 | ANT3-DPRIME | 20.000 | 7.303 | 0.216 | 0.079 | 1.000 | 0.920 | 0.080 | 0.000 | 0.000 | 0.000 | 0.000 |
| 22 | 555 | QNRS | 17.280 | 7.056 | 0.223 | 0.088 | 1.000 | 0.480 | 0.520 | 0.000 | 0.000 | 0.000 | 0.000 |
| 23 | 558 | VANWI | 22.360 | 10.012 | 0.170 | 0.078 | 1.000 | 1.000 | 0.000 | 0.000 | 0.000 | 0.000 | 0.000 |
| 24 | 578 | TETX | 39.200 | 17.830 | 0.296 | 0.136 | 1.000 | 0.400 | 0.600 | 0.000 | 0.000 | 0.000 | 0.000 |
| 25 | 652 | VGA | 34.680 | 12.730 | 0.198 | 0.074 | 1.000 | 0.800 | 0.200 | 0.000 | 0.000 | 0.000 | 0.000 |
| 26 | 682 | MOX | 33.680 | 15.063 | 0.254 | 0.111 | 1.000 | 0.240 | 0.760 | 0.000 | 0.000 | 0.000 | 0.000 |
| 27 | 694 | ACT | 35.720 | 17.021 | 0.247 | 0.125 | 1.000 | 0.080 | 0.920 | 0.000 | 0.000 | 0.000 | 0.000 |
| 28 | 717 | VANRE | 25.720 | 11.516 | 0.334 | 0.151 | 1.000 | 1.000 | 0.000 | 0.000 | 0.000 | 0.000 | 0.000 |
| 29 | 749 | RMTB | 7.880 | 4.324 | 0.274 | 0.153 | 1.000 | 0.920 | 0.080 | 0.000 | 0.000 | 0.000 | 0.000 |
| 30 | 778 | DHA | 21.640 | 10.950 | 0.165 | 0.081 | 1.000 | 0.040 | 0.960 | 0.000 | 0.000 | 0.000 | 0.000 |
| 31 | 789 | CMY | 23.080 | 14.130 | 0.174 | 0.112 | 1.000 | 0.000 | 1.000 | 0.000 | 0.000 | 0.000 | 0.000 |
| 32 | 797 | FACT | 36.240 | 27.470 | 0.193 | 0.145 | 1.000 | 1.000 | 0.000 | 0.000 | 0.000 | 0.000 | 0.000 |
| 33 | 819 | OKP | 27.560 | 13.482 | 0.283 | 0.142 | 1.000 | 0.080 | 0.920 | 0.000 | 0.000 | 0.000 | 0.000 |
| 34 | 973 | IMP | 16.640 | 7.262 | 0.199 | 0.082 | 1.000 | 0.160 | 0.840 | 0.000 | 0.000 | 0.000 | 0.000 |
| 35 | 1048 | VIM | 15.280 | 6.175 | 0.164 | 0.068 | 1.000 | 0.040 | 0.960 | 0.000 | 0.000 | 0.000 | 0.000 |
| 36 | 1135 | PBP1B | 31.200 | 14.491 | 0.110 | 0.050 | 1.000 | 1.000 | 0.000 | 0.000 | 0.000 | 0.000 | 0.000 |
| 37 | 1146 | MPHE | 25.120 | 14.432 | 0.251 | 0.145 | 1.000 | 0.360 | 0.640 | 0.000 | 0.000 | 0.000 | 0.000 |
| 38 | 1182 | VATE | 13.800 | 7.439 | 0.188 | 0.101 | 1.000 | 0.480 | 0.520 | 0.000 | 0.000 | 0.000 | 0.000 |
| 39 | 1214 | OPRJ | 50.280 | 25.842 | 0.310 | 0.158 | 1.000 | 1.000 | 0.000 | 0.000 | 0.000 | 0.000 | 0.000 |
| 40 | 1254 | FOSB | 18.320 | 8.265 | 0.387 | 0.173 | 1.000 | 0.960 | 0.040 | 0.000 | 0.000 | 0.000 | 0.000 |
| 41 | 1271 | VPH | 25.520 | 13.080 | 0.264 | 0.137 | 1.000 | 1.000 | 0.000 | 0.000 | 0.000 | 0.000 | 0.000 |
| 42 | 1283 | SULI | 40.680 | 19.786 | 0.439 | 0.240 | 1.000 | 0.120 | 0.880 | 0.000 | 0.000 | 0.000 | 0.000 |
| 43 | 1297 | TET40 | 17.560 | 8.466 | 0.125 | 0.060 | 1.000 | 0.360 | 0.640 | 0.000 | 0.000 | 0.000 | 0.000 |
| 44 | 1389 | CPXAR | 19.240 | 8.141 | 0.234 | 0.096 | 1.000 | 0.400 | 0.600 | 0.000 | 0.000 | 0.000 | 0.000 |
| 45 | 1392 | AAC6-PRIME | 10.680 | 4.589 | 0.167 | 0.072 | 1.000 | 0.680 | 0.320 | 0.000 | 0.000 | 0.000 | 0.000 |
| 46 | 1422 | VGBB | 26.480 | 10.813 | 0.260 | 0.105 | 1.000 | 1.000 | 0.000 | 0.000 | 0.000 | 0.000 | 0.000 |
| 47 | 1440 | FOSC | 14.520 | 7.343 | 0.223 | 0.112 | 1.000 | 1.000 | 0.000 | 0.000 | 0.000 | 0.000 | 0.000 |

| | | | | | | | | | | | | | |
|---|---|---|---|---|---|---|---|---|---|---|---|---|---|
| 48 | 1535 | LNUA | 10.400 | 4.444 | 0.196 | 0.080 | 1.000 | 0.440 | 0.560 | 0.000 | 0.000 | 0.000 | 0.000 |
| 49 | 1569 | PARE | 38.560 | 13.863 | 0.179 | 0.065 | 1.000 | 0.640 | 0.360 | 0.000 | 0.000 | 0.000 | 0.000 |
| 50 | 1695 | NDM | 22.400 | 7.303 | 0.238 | 0.077 | 1.000 | 1.000 | 0.000 | 0.000 | 0.000 | 0.000 | 0.000 |
| 51 | 1702 | SPG | 19.400 | 12.295 | 0.197 | 0.128 | 1.000 | 1.000 | 0.000 | 0.000 | 0.000 | 0.000 | 0.000 |
| 52 | 1753 | VEB | 28.760 | 17.408 | 0.286 | 0.175 | 1.000 | 0.040 | 0.960 | 0.000 | 0.000 | 0.000 | 0.000 |
| 53 | 1953 | LNUB | 29.760 | 15.517 | 0.334 | 0.177 | 1.000 | 0.640 | 0.360 | 0.000 | 0.000 | 0.000 | 0.000 |
| 54 | 2026 | ERMA | 22.400 | 10.704 | 0.272 | 0.128 | 1.000 | 0.720 | 0.280 | 0.000 | 0.000 | 0.000 | 0.000 |
| 55 | 2357 | SULII | 20.800 | 10.165 | 0.222 | 0.108 | 1.000 | 0.160 | 0.840 | 0.000 | 0.000 | 0.000 | 0.000 |
| 56 | 2517 | TET37 | 9.560 | 4.620 | 0.248 | 0.116 | 1.000 | 1.000 | 0.000 | 0.000 | 0.000 | 0.000 | 0.000 |
| 57 | 2822 | EMRK | 30.040 | 12.684 | 0.250 | 0.108 | 1.000 | 1.000 | 0.000 | 0.000 | 0.000 | 0.000 | 0.000 |
| 58 | 2999 | MPHB | 22.880 | 8.710 | 0.219 | 0.082 | 1.000 | 0.960 | 0.040 | 0.000 | 0.000 | 0.000 | 0.000 |
| 59 | 3024 | VANYM | 21.800 | 9.954 | 0.273 | 0.132 | 1.000 | 1.000 | 0.000 | 0.000 | 0.000 | 0.000 | 0.000 |
| 60 | 3041 | MECC | 127.480 | 77.287 | 0.624 | 0.394 | 1.000 | 0.560 | 0.440 | 0.000 | 0.000 | 0.000 | 0.000 |
| 61 | 3128 | TUFAB | 25.120 | 11.344 | 0.182 | 0.081 | 1.000 | 0.560 | 0.440 | 0.000 | 0.000 | 0.000 | 0.000 |
| 62 | 3176 | AMRB | 60.080 | 30.228 | 0.169 | 0.084 | 1.000 | 1.000 | 0.000 | 0.000 | 0.000 | 0.000 | 0.000 |
| 63 | 3270 | IRI | 44.600 | 17.550 | 0.267 | 0.107 | 1.000 | 1.000 | 0.000 | 0.000 | 0.000 | 0.000 | 0.000 |
| 64 | 3314 | RPOB | 46.760 | 17.439 | 0.114 | 0.043 | 1.000 | 0.080 | 0.920 | 0.000 | 0.000 | 0.080 | 0.277 |
| 65 | 3332 | TET35 | 19.320 | 8.697 | 0.148 | 0.066 | 1.000 | 1.000 | 0.000 | 0.000 | 0.000 | 0.000 | 0.000 |
| 66 | 3370 | CFRA | 35.440 | 13.476 | 0.301 | 0.116 | 1.000 | 1.000 | 0.000 | 0.000 | 0.000 | 0.000 | 0.000 |
| 67 | 3513 | BRP | 9.760 | 4.893 | 0.212 | 0.107 | 1.000 | 1.000 | 0.000 | 0.000 | 0.000 | 0.000 | 0.000 |
| 68 | 3613 | APH3-PRIME | 24.520 | 9.005 | 0.269 | 0.098 | 1.000 | 0.960 | 0.040 | 0.000 | 0.000 | 0.000 | 0.000 |
| 69 | 3697 | TETM | 47.120 | 20.376 | 0.219 | 0.094 | 1.000 | 0.600 | 0.400 | 0.000 | 0.000 | 0.000 | 0.000 |
| 70 | 3778 | IND | 17.160 | 6.743 | 0.208 | 0.079 | 1.000 | 0.040 | 0.960 | 0.000 | 0.000 | 0.000 | 0.000 |

Simulation settings:

k-mers: ‘variable’, k=10
Gene coverage: 1
Number of genes: 1
Errors: ‘off’
Penalty score: 0.1
Thresholding: multiplier=1, all factors (1-on)
Entropy screening: ‘rand’

**Table S7.** Resistance genes simulations with 0.01 (2%) error rate

| No. | Database No. | Gene sub-class | Avg block for identification | StDev block for identification | Avg coverage for identification | StDev coverage for identification | Accuracy | Identification level (fraction of trials) | | | | Avg false positives | StDev false positives |
|---|---|---|---|---|---|---|---|---|---|---|---|---|---|
| | | | | | | | | Specific gene | Sub-class | Class | Incorrect | | |
| 1 | 4 | VANZA | 23.500 | 12.786 | 0.478 | 0.262 | 0.960 | 0.960 | 0.000 | 0.000 | 0.040 | 0.000 | 0.000 |
| 2 | 52 | VANWG | 47.280 | 18.573 | 0.555 | 0.218 | 1.000 | 0.400 | 0.600 | 0.000 | 0.000 | 0.000 | 0.000 |
| 3 | 62 | CTX | 56.480 | 12.600 | 0.640 | 0.144 | 1.000 | 0.000 | 1.000 | 0.000 | 0.000 | 0.040 | 0.200 |
| 4 | 68 | MEFA | 78.560 | 17.280 | 0.640 | 0.141 | 1.000 | 0.640 | 0.360 | 0.000 | 0.000 | 0.000 | 0.000 |
| 5 | 76 | OXA | 46.320 | 16.175 | 0.556 | 0.194 | 1.000 | 0.040 | 0.960 | 0.000 | 0.000 | 0.000 | 0.000 |
| 6 | 92 | CATB | 18.880 | 12.601 | 0.291 | 0.195 | 1.000 | 0.280 | 0.720 | 0.000 | 0.000 | 0.000 | 0.000 |
| 7 | 162 | EREA | 26.440 | 15.578 | 0.214 | 0.126 | 1.000 | 0.480 | 0.520 | 0.000 | 0.000 | 0.000 | 0.000 |
| 8 | 174 | TEM | 35.960 | 7.829 | 0.424 | 0.093 | 1.000 | 0.280 | 0.720 | 0.000 | 0.000 | 0.000 | 0.000 |
| 9 | 193 | CML | 66.320 | 17.587 | 0.500 | 0.133 | 1.000 | 0.240 | 0.720 | 0.040 | 0.000 | 0.000 | 0.000 |
| 10 | 207 | NIMA | 12.720 | 6.222 | 0.272 | 0.136 | 1.000 | 1.000 | 0.000 | 0.000 | 0.000 | 0.000 | 0.000 |
| 11 | 239 | DHFR | 26.000 | 7.348 | 0.536 | 0.154 | 1.000 | 0.080 | 0.920 | 0.000 | 0.000 | 0.000 | 0.000 |
| 12 | 246 | FOLP | 90.174 | 16.420 | 0.755 | 0.140 | 0.920 | 0.920 | 0.000 | 0.000 | 0.080 | 0.000 | 0.000 |
| 13 | 274 | PARC | 189.333 | 43.389 | 0.849 | 0.196 | 0.840 | 0.040 | 0.520 | 0.280 | 0.160 | 1.560 | 2.142 |
| 14 | 276 | SHV | 78.840 | 12.229 | 0.899 | 0.140 | 1.000 | 0.040 | 0.960 | 0.000 | 0.000 | 0.560 | 0.583 |
| 15 | 284 | DFRA | 30.880 | 9.198 | 0.546 | 0.165 | 1.000 | 1.000 | 0.000 | 0.000 | 0.000 | 0.000 | 0.000 |
| 16 | 295 | SOXS | 17.360 | 6.794 | 0.520 | 0.208 | 1.000 | 1.000 | 0.000 | 0.000 | 0.000 | 0.000 | 0.000 |
| 17 | 321 | TSNR | 47.833 | 12.426 | 0.603 | 0.157 | 0.960 | 0.960 | 0.000 | 0.000 | 0.040 | 0.000 | 0.000 |
| 18 | 413 | OMPD | 57.250 | 9.755 | 0.528 | 0.090 | 0.960 | 0.960 | 0.000 | 0.000 | 0.040 | 0.000 | 0.000 |
| 19 | 486 | LMRA | 20.280 | 8.463 | 0.350 | 0.144 | 1.000 | 1.000 | 0.000 | 0.000 | 0.000 | 0.000 | 0.000 |
| 20 | 505 | CARB | 52.200 | 8.446 | 0.574 | 0.092 | 1.000 | 0.960 | 0.000 | 0.040 | 0.000 | 0.000 | 0.000 |
| 21 | 506 | ANT3-DPRIME | 41.880 | 13.318 | 0.519 | 0.168 | 1.000 | 0.920 | 0.080 | 0.000 | 0.000 | 0.240 | 0.831 |
| 22 | 555 | QNRS | 35.840 | 5.749 | 0.534 | 0.093 | 1.000 | 0.600 | 0.400 | 0.000 | 0.000 | 0.000 | 0.000 |
| 23 | 558 | VANWI | 31.520 | 10.190 | 0.277 | 0.090 | 1.000 | 1.000 | 0.000 | 0.000 | 0.000 | 0.000 | 0.000 |
| 24 | 578 | TETX | 84.280 | 14.932 | 0.717 | 0.129 | 1.000 | 0.840 | 0.160 | 0.000 | 0.000 | 0.080 | 0.277 |
| 25 | 652 | VGA | 101.040 | 36.600 | 0.639 | 0.231 | 1.000 | 0.920 | 0.080 | 0.000 | 0.000 | 0.560 | 1.960 |
| 26 | 682 | MOX | 96.800 | 16.889 | 0.836 | 0.145 | 1.000 | 0.440 | 0.560 | 0.000 | 0.000 | 0.280 | 0.458 |
| 27 | 694 | ACT | 86.240 | 14.878 | 0.686 | 0.119 | 1.000 | 0.240 | 0.760 | 0.000 | 0.000 | 0.040 | 0.200 |
| 28 | 717 | VANRE | 51.083 | 9.329 | 0.727 | 0.133 | 0.960 | 0.960 | 0.000 | 0.000 | 0.040 | 0.000 | 0.000 |
| 29 | 749 | RMTB | 10.542 | 3.538 | 0.394 | 0.138 | 0.960 | 0.960 | 0.000 | 0.000 | 0.040 | 0.000 | 0.000 |
| 30 | 778 | DHA | 48.080 | 16.457 | 0.416 | 0.144 | 1.000 | 0.040 | 0.960 | 0.000 | 0.000 | 0.000 | 0.000 |
| 31 | 789 | CMY | 33.520 | 14.477 | 0.290 | 0.126 | 1.000 | 0.000 | 1.000 | 0.000 | 0.000 | 0.000 | 0.000 |
| 32 | 797 | FACT | 139.222 | 17.795 | 0.823 | 0.106 | 0.720 | 0.680 | 0.040 | 0.000 | 0.280 | 0.080 | 0.400 |
| 33 | 819 | OKP | 65.583 | 9.169 | 0.751 | 0.106 | 0.960 | 0.120 | 0.840 | 0.000 | 0.040 | 0.080 | 0.277 |
| 34 | 973 | IMP | 28.400 | 16.946 | 0.378 | 0.225 | 1.000 | 0.240 | 0.760 | 0.000 | 0.000 | 0.000 | 0.000 |
| 35 | 1048 | VIM | 31.920 | 12.945 | 0.390 | 0.158 | 1.000 | 0.040 | 0.960 | 0.000 | 0.000 | 0.000 | 0.000 |
| 36 | 1135 | PBP1B | 79.000 | 58.407 | 0.319 | 0.235 | 0.840 | 0.760 | 0.040 | 0.040 | 0.160 | 0.040 | 0.200 |
| 37 | 1146 | MPHE | 69.000 | 6.285 | 0.774 | 0.071 | 1.000 | 0.440 | 0.560 | 0.000 | 0.000 | 0.000 | 0.000 |
| 38 | 1182 | VATE | 39.917 | 8.387 | 0.611 | 0.129 | 0.960 | 0.640 | 0.320 | 0.000 | 0.040 | 0.000 | 0.000 |
| 39 | 1214 | OPRJ | 130.773 | 14.784 | 0.903 | 0.103 | 0.880 | 0.480 | 0.400 | 0.000 | 0.120 | 0.480 | 0.586 |
| 40 | 1254 | FOSB | 31.120 | 9.816 | 0.722 | 0.229 | 1.000 | 0.800 | 0.200 | 0.000 | 0.000 | 0.000 | 0.000 |
| 41 | 1271 | VPH | 46.208 | 28.717 | 0.530 | 0.330 | 0.960 | 0.920 | 0.040 | 0.000 | 0.040 | 0.080 | 0.277 |
| 42 | 1283 | SULI | 61.080 | 9.617 | 0.717 | 0.116 | 1.000 | 0.240 | 0.760 | 0.000 | 0.000 | 0.040 | 0.200 |
| 43 | 1297 | TET40 | 51.960 | 30.194 | 0.420 | 0.245 | 1.000 | 0.640 | 0.360 | 0.000 | 0.000 | 0.000 | 0.000 |
| 44 | 1389 | CPXAR | 31.320 | 11.089 | 0.438 | 0.159 | 1.000 | 0.320 | 0.640 | 0.040 | 0.000 | 0.360 | 1.440 |
| 45 | 1392 | AAC6-PRIME | 11.240 | 7.639 | 0.198 | 0.136 | 1.000 | 0.760 | 0.240 | 0.000 | 0.000 | 0.000 | 0.000 |
| 46 | 1422 | VGBB | 64.750 | 8.828 | 0.723 | 0.100 | 0.960 | 0.920 | 0.040 | 0.000 | 0.040 | 0.120 | 0.600 |
| 47 | 1440 | FOSC | 24.920 | 7.433 | 0.438 | 0.131 | 1.000 | 1.000 | 0.000 | 0.000 | 0.000 | 0.000 | 0.000 |

| | | | | | | | | | | | | | |
|---|---|---|---|---|---|---|---|---|---|---|---|---|---|
| 48 | 1535 | LNUA | 13.960 | 9.312 | 0.285 | 0.190 | 1.000 | 0.720 | 0.280 | 0.000 | 0.000 | 0.000 | 0.000 |
| 49 | 1569 | PARE | 136.143 | 36.753 | 0.719 | 0.195 | 0.280 | 0.200 | 0.080 | 0.000 | 0.720 | 0.680 | 2.358 |
| 50 | 1695 | NDM | 45.875 | 9.396 | 0.557 | 0.114 | 0.960 | 0.960 | 0.000 | 0.000 | 0.040 | 0.000 | 0.000 |
| 51 | 1702 | SPG | 60.667 | 10.655 | 0.700 | 0.124 | 0.960 | 0.920 | 0.040 | 0.000 | 0.040 | 0.040 | 0.200 |
| 52 | 1753 | VEB | 76.520 | 9.430 | 0.844 | 0.103 | 1.000 | 0.000 | 1.000 | 0.000 | 0.000 | 0.040 | 0.200 |
| 53 | 1953 | LNUB | 63.760 | 6.437 | 0.786 | 0.080 | 1.000 | 0.960 | 0.040 | 0.000 | 0.000 | 0.000 | 0.000 |
| 54 | 2026 | ERMA | 47.000 | 18.815 | 0.633 | 0.254 | 1.000 | 0.880 | 0.080 | 0.040 | 0.000 | 0.200 | 0.500 |
| 55 | 2357 | SULII | 53.800 | 10.079 | 0.654 | 0.124 | 1.000 | 0.080 | 0.920 | 0.000 | 0.000 | 0.080 | 0.277 |
| 56 | 2517 | TET37 | 18.958 | 2.911 | 0.560 | 0.091 | 0.960 | 0.960 | 0.000 | 0.000 | 0.040 | 0.000 | 0.000 |
| 57 | 2822 | EMRK | 71.920 | 12.566 | 0.676 | 0.120 | 1.000 | 0.920 | 0.000 | 0.080 | 0.000 | 0.320 | 1.145 |
| 58 | 2999 | MPHB | 57.120 | 8.550 | 0.619 | 0.096 | 1.000 | 1.000 | 0.000 | 0.000 | 0.000 | 0.000 | 0.000 |
| 59 | 3024 | VANYM | 51.640 | 7.979 | 0.724 | 0.114 | 1.000 | 0.960 | 0.040 | 0.000 | 0.000 | 0.040 | 0.200 |
| 60 | 3041 | MECC | 175.360 | 40.240 | 0.870 | 0.200 | 1.000 | 0.440 | 0.560 | 0.000 | 0.000 | 0.000 | 0.000 |
| 61 | 3128 | TUFAB | 51.833 | 21.184 | 0.434 | 0.178 | 0.960 | 0.760 | 0.200 | 0.000 | 0.040 | 0.000 | 0.000 |
| 62 | 3176 | AMRB | 296.600 | 54.895 | 0.943 | 0.175 | 1.000 | 0.040 | 0.840 | 0.120 | 0.000 | 0.080 | 0.400 |
| 63 | 3270 | IRI | 125.708 | 22.542 | 0.866 | 0.156 | 0.960 | 0.520 | 0.440 | 0.000 | 0.040 | 0.480 | 0.586 |
| 64 | 3314 | RPOB | 336.286 | 52.159 | 0.944 | 0.147 | 0.280 | 0.000 | 0.280 | 0.000 | 0.720 | 1.480 | 2.756 |
| 65 | 3332 | TET35 | 43.043 | 9.943 | 0.383 | 0.088 | 0.920 | 0.920 | 0.000 | 0.000 | 0.080 | 0.000 | 0.000 |
| 66 | 3370 | CFRA | 78.773 | 13.245 | 0.745 | 0.124 | 0.880 | 0.840 | 0.040 | 0.000 | 0.120 | 0.040 | 0.200 |
| 67 | 3513 | BRP | 15.440 | 8.150 | 0.373 | 0.198 | 1.000 | 1.000 | 0.000 | 0.000 | 0.000 | 0.000 | 0.000 |
| 68 | 3613 | APH3-PRIME | 56.800 | 4.213 | 0.698 | 0.053 | 1.000 | 1.000 | 0.000 | 0.000 | 0.000 | 0.000 | 0.000 |
| 69 | 3697 | TETM | 168.400 | 19.807 | 0.872 | 0.103 | 1.000 | 0.760 | 0.240 | 0.000 | 0.000 | 0.360 | 0.700 |
| 70 | 3778 | IND | 51.680 | 10.523 | 0.695 | 0.143 | 1.000 | 0.080 | 0.920 | 0.000 | 0.000 | 0.000 | 0.000 |

Simulation settings:

k-mers: ‘constant’, k=10
Gene coverage: 1
Number of genes: 1
Errors: ‘on’, 0.01
Penalty score: 0.1
Thresholding: multiplier=2, all factors (1-on)
Entropy screening: ‘rand’

**Table S8.** Resistance genes simulations with 0.025 (5%) error rate

| No. | Database No. | Gene sub-class | Avg block for identification | StDev block for identification | Avg coverage for identification | StDev coverage for identification | Accuracy | Identification level (fraction of trials) | | | | Avg false positives | StDev false positives |
|---|---|---|---|---|---|---|---|---|---|---|---|---|---|
| | | | | | | | | Specific gene | Sub-class | Class | Incorrect | | |
| 1 | 4 | VANZA | 25.250 | 10.707 | 0.514 | 0.218 | 0.960 | 0.960 | 0.000 | 0.000 | 0.040 | 0.040 | 0.200 |
| 2 | 52 | VANWG | 53.960 | 14.438 | 0.633 | 0.170 | 1.000 | 0.440 | 0.560 | 0.000 | 0.000 | 0.000 | 0.000 |
| 3 | 62 | CTX | 60.440 | 21.370 | 0.686 | 0.243 | 1.000 | 0.000 | 1.000 | 0.000 | 0.000 | 0.080 | 0.277 |
| 4 | 68 | MEFA | 85.040 | 11.040 | 0.694 | 0.090 | 1.000 | 0.560 | 0.440 | 0.000 | 0.000 | 0.040 | 0.200 |
| 5 | 76 | OXA | 44.800 | 19.462 | 0.538 | 0.235 | 1.000 | 0.000 | 1.000 | 0.000 | 0.000 | 0.200 | 1.000 |
| 6 | 92 | CATB | 24.160 | 13.966 | 0.373 | 0.218 | 1.000 | 0.360 | 0.640 | 0.000 | 0.000 | 0.000 | 0.000 |
| 7 | 162 | EREA | 25.174 | 15.614 | 0.204 | 0.125 | 0.920 | 0.520 | 0.400 | 0.000 | 0.080 | 0.040 | 0.200 |
| 8 | 174 | TEM | 44.320 | 16.663 | 0.525 | 0.198 | 1.000 | 0.120 | 0.880 | 0.000 | 0.000 | 0.160 | 0.374 |
| 9 | 193 | CML | 61.840 | 26.100 | 0.467 | 0.197 | 1.000 | 0.160 | 0.800 | 0.040 | 0.000 | 0.040 | 0.200 |
| 10 | 207 | NIMA | 17.375 | 10.034 | 0.373 | 0.219 | 0.960 | 0.920 | 0.040 | 0.000 | 0.040 | 0.000 | 0.000 |
| 11 | 239 | DHFR | 24.800 | 8.860 | 0.513 | 0.183 | 1.000 | 0.320 | 0.680 | 0.000 | 0.000 | 0.000 | 0.000 |
| 12 | 246 | FOLP | 89.080 | 24.605 | 0.746 | 0.208 | 1.000 | 0.920 | 0.080 | 0.000 | 0.000 | 0.080 | 0.277 |
| 13 | 274 | PARC | 202.778 | 34.739 | 0.909 | 0.157 | 0.720 | 0.080 | 0.400 | 0.240 | 0.280 | 2.080 | 2.465 |
| 14 | 276 | SHV | 76.640 | 17.464 | 0.874 | 0.198 | 1.000 | 0.000 | 1.000 | 0.000 | 0.000 | 1.000 | 1.958 |
| 15 | 284 | DFRA | 33.480 | 11.748 | 0.594 | 0.211 | 1.000 | 0.920 | 0.080 | 0.000 | 0.000 | 0.240 | 0.597 |
| 16 | 295 | SOXS | 18.560 | 5.752 | 0.556 | 0.177 | 1.000 | 0.960 | 0.040 | 0.000 | 0.000 | 0.240 | 1.200 |
| 17 | 321 | TSNR | 57.280 | 13.107 | 0.723 | 0.167 | 1.000 | 0.880 | 0.120 | 0.000 | 0.000 | 0.280 | 0.542 |
| 18 | 413 | OMPD | 63.292 | 12.896 | 0.584 | 0.120 | 0.960 | 0.920 | 0.040 | 0.000 | 0.040 | 0.040 | 0.200 |
| 19 | 486 | LMRA | 24.708 | 7.214 | 0.422 | 0.124 | 0.960 | 0.960 | 0.000 | 0.000 | 0.040 | 0.000 | 0.000 |
| 20 | 505 | CARB | 54.200 | 18.538 | 0.598 | 0.204 | 1.000 | 0.880 | 0.040 | 0.080 | 0.000 | 0.320 | 0.900 |
| 21 | 506 | ANT3-DPRIME | 44.080 | 12.486 | 0.547 | 0.157 | 1.000 | 0.920 | 0.080 | 0.000 | 0.000 | 0.240 | 1.200 |
| 22 | 555 | QNRS | 39.080 | 6.416 | 0.588 | 0.098 | 1.000 | 0.480 | 0.520 | 0.000 | 0.000 | 0.080 | 0.277 |
| 23 | 558 | VANWI | 40.760 | 10.345 | 0.358 | 0.092 | 1.000 | 1.000 | 0.000 | 0.000 | 0.000 | 0.040 | 0.200 |
| 24 | 578 | TETX | 89.960 | 18.620 | 0.767 | 0.161 | 1.000 | 0.680 | 0.320 | 0.000 | 0.000 | 0.200 | 0.408 |
| 25 | 652 | VGA | 102.120 | 32.917 | 0.646 | 0.208 | 1.000 | 0.920 | 0.080 | 0.000 | 0.000 | 0.120 | 0.332 |
| 26 | 682 | MOX | 106.080 | 13.617 | 0.918 | 0.119 | 1.000 | 0.240 | 0.760 | 0.000 | 0.000 | 0.640 | 0.638 |
| 27 | 694 | ACT | 94.800 | 17.424 | 0.756 | 0.140 | 1.000 | 0.360 | 0.640 | 0.000 | 0.000 | 0.160 | 0.374 |
| 28 | 717 | VANRE | 56.720 | 7.992 | 0.808 | 0.116 | 1.000 | 0.880 | 0.120 | 0.000 | 0.000 | 0.160 | 0.374 |
| 29 | 749 | RMTB | 11.440 | 6.905 | 0.433 | 0.265 | 1.000 | 0.880 | 0.120 | 0.000 | 0.000 | 0.520 | 2.220 |
| 30 | 778 | DHA | 41.760 | 17.429 | 0.361 | 0.151 | 1.000 | 0.000 | 1.000 | 0.000 | 0.000 | 0.080 | 0.400 |
| 31 | 789 | CMY | 38.640 | 21.022 | 0.335 | 0.183 | 1.000 | 0.000 | 0.960 | 0.040 | 0.000 | 0.000 | 0.000 |
| 32 | 797 | FACT | 147.944 | 22.161 | 0.875 | 0.132 | 0.720 | 0.480 | 0.240 | 0.000 | 0.280 | 0.280 | 0.542 |
| 33 | 819 | OKP | 79.375 | 9.050 | 0.911 | 0.105 | 0.960 | 0.120 | 0.800 | 0.040 | 0.040 | 0.520 | 0.653 |
| 34 | 973 | IMP | 42.040 | 17.862 | 0.557 | 0.239 | 1.000 | 0.120 | 0.880 | 0.000 | 0.000 | 0.040 | 0.200 |
| 35 | 1048 | VIM | 36.840 | 12.233 | 0.450 | 0.151 | 1.000 | 0.000 | 1.000 | 0.000 | 0.000 | 0.000 | 0.000 |
| 36 | 1135 | PBP1B | 87.389 | 31.797 | 0.353 | 0.128 | 0.720 | 0.720 | 0.000 | 0.000 | 0.280 | 0.080 | 0.400 |
| 37 | 1146 | MPHE | 69.833 | 7.063 | 0.783 | 0.080 | 0.960 | 0.560 | 0.360 | 0.040 | 0.040 | 0.280 | 1.400 |
| 38 | 1182 | VATE | 44.200 | 8.067 | 0.678 | 0.124 | 1.000 | 0.520 | 0.480 | 0.000 | 0.000 | 0.040 | 0.200 |
| 39 | 1214 | OPRJ | 138.304 | 15.423 | 0.955 | 0.107 | 0.920 | 0.200 | 0.720 | 0.000 | 0.080 | 0.880 | 0.726 |
| 40 | 1254 | FOSB | 34.480 | 7.495 | 0.805 | 0.175 | 1.000 | 0.960 | 0.040 | 0.000 | 0.000 | 0.000 | 0.000 |
| 41 | 1271 | VPH | 52.957 | 25.121 | 0.607 | 0.289 | 0.920 | 0.840 | 0.080 | 0.000 | 0.080 | 0.200 | 0.577 |
| 42 | 1283 | SULI | 65.840 | 12.202 | 0.774 | 0.145 | 1.000 | 0.120 | 0.880 | 0.000 | 0.000 | 0.240 | 0.663 |
| 43 | 1297 | TET40 | 35.833 | 24.925 | 0.290 | 0.201 | 0.960 | 0.400 | 0.560 | 0.000 | 0.040 | 0.000 | 0.000 |
| 44 | 1389 | CPXAR | 35.360 | 15.047 | 0.498 | 0.212 | 1.000 | 0.400 | 0.560 | 0.040 | 0.000 | 0.120 | 0.440 |
| 45 | 1392 | AAC6-PRIME | 12.960 | 6.661 | 0.230 | 0.119 | 1.000 | 0.920 | 0.080 | 0.000 | 0.000 | 0.000 | 0.000 |
| 46 | 1422 | VGBB | 68.640 | 7.416 | 0.766 | 0.084 | 1.000 | 0.960 | 0.040 | 0.000 | 0.000 | 0.080 | 0.277 |
| 47 | 1440 | FOSC | 30.600 | 5.635 | 0.543 | 0.099 | 1.000 | 1.000 | 0.000 | 0.000 | 0.000 | 0.000 | 0.000 |

| | | | | | | | | | | | | | |
|---|---|---|---|---|---|---|---|---|---|---|---|---|---|
| 48 | 1535 | LNUA | 18.120 | 10.553 | 0.369 | 0.215 | 1.000 | 0.560 | 0.440 | 0.000 | 0.000 | 0.080 | 0.277 |
| 49 | 1569 | PARE | 151.000 | 41.661 | 0.798 | 0.221 | 0.240 | 0.040 | 0.120 | 0.080 | 0.760 | 0.680 | 2.096 |
| 50 | 1695 | NDM | 50.625 | 16.248 | 0.615 | 0.198 | 0.960 | 0.920 | 0.000 | 0.040 | 0.040 | 0.200 | 1.000 |
| 51 | 1702 | SPG | 70.042 | 13.013 | 0.808 | 0.152 | 0.960 | 0.760 | 0.160 | 0.040 | 0.040 | 0.600 | 1.528 |
| 52 | 1753 | VEB | 72.920 | 15.242 | 0.804 | 0.168 | 1.000 | 0.000 | 0.960 | 0.040 | 0.000 | 0.320 | 0.988 |
| 53 | 1953 | LNUB | 66.480 | 11.435 | 0.820 | 0.142 | 1.000 | 0.720 | 0.280 | 0.000 | 0.000 | 0.200 | 0.408 |
| 54 | 2026 | ERMA | 54.440 | 13.238 | 0.734 | 0.180 | 1.000 | 0.720 | 0.240 | 0.040 | 0.000 | 0.160 | 0.473 |
| 55 | 2357 | SULII | 53.792 | 11.310 | 0.654 | 0.138 | 0.960 | 0.040 | 0.920 | 0.000 | 0.040 | 0.040 | 0.200 |
| 56 | 2517 | TET37 | 19.200 | 6.344 | 0.571 | 0.195 | 1.000 | 1.000 | 0.000 | 0.000 | 0.000 | 0.000 | 0.000 |
| 57 | 2822 | EMRK | 69.833 | 13.127 | 0.656 | 0.125 | 0.960 | 0.880 | 0.040 | 0.040 | 0.040 | 0.160 | 0.624 |
| 58 | 2999 | MPHB | 57.583 | 15.234 | 0.628 | 0.169 | 0.960 | 0.920 | 0.040 | 0.000 | 0.040 | 0.040 | 0.200 |
| 59 | 3024 | VANYM | 52.320 | 8.669 | 0.734 | 0.123 | 1.000 | 0.960 | 0.040 | 0.000 | 0.000 | 0.160 | 0.624 |
| 60 | 3041 | MECC | 172.200 | 24.767 | 0.855 | 0.124 | 1.000 | 0.600 | 0.360 | 0.040 | 0.000 | 0.200 | 1.000 |
| 61 | 3128 | TUFAB | 67.160 | 25.151 | 0.563 | 0.212 | 1.000 | 0.720 | 0.280 | 0.000 | 0.000 | 0.280 | 1.400 |
| 62 | 3176 | AMRB | 308.640 | 16.830 | 0.981 | 0.054 | 1.000 | 0.120 | 0.840 | 0.040 | 0.000 | 0.080 | 0.400 |
| 63 | 3270 | IRI | 133.200 | 16.427 | 0.918 | 0.114 | 1.000 | 0.440 | 0.560 | 0.000 | 0.000 | 0.680 | 0.748 |
| 64 | 3314 | RPOB | 331.667 | 46.523 | 0.931 | 0.131 | 0.480 | 0.000 | 0.480 | 0.000 | 0.520 | 2.440 | 3.429 |
| 65 | 3332 | TET35 | 48.917 | 11.938 | 0.435 | 0.108 | 0.960 | 0.960 | 0.000 | 0.000 | 0.040 | 0.000 | 0.000 |
| 66 | 3370 | CFRA | 87.857 | 8.248 | 0.832 | 0.079 | 0.840 | 0.840 | 0.000 | 0.000 | 0.160 | 0.000 | 0.000 |
| 67 | 3513 | BRP | 17.000 | 8.886 | 0.414 | 0.217 | 0.960 | 0.960 | 0.000 | 0.000 | 0.040 | 0.000 | 0.000 |
| 68 | 3613 | APH3-PRIME | 60.240 | 7.833 | 0.741 | 0.097 | 1.000 | 0.920 | 0.080 | 0.000 | 0.000 | 0.080 | 0.277 |
| 69 | 3697 | TETM | 175.880 | 19.951 | 0.912 | 0.104 | 1.000 | 0.400 | 0.600 | 0.000 | 0.000 | 1.400 | 1.979 |
| 70 | 3778 | IND | 56.840 | 10.862 | 0.766 | 0.148 | 1.000 | 0.080 | 0.880 | 0.040 | 0.000 | 0.360 | 1.036 |

Simulation settings:

k-mers: ‘constant’, k=10
Gene coverage: 1
Number of genes: 1
Errors: ‘on’, 0.025
Penalty score: 0.1
Thresholding: multiplier=2, all factors (1-on)
Entropy screening: ‘rand’

**Table S9.** Resistance genes simulations with 0.05 (10%) error rate

| No. | Database No. | Gene sub-class | Avg block for identification | StDev block for identification | Avg coverage for identification | StDev coverage for identification | Accuracy | Identification level (fraction of trials) | | | | Avg false positives | StDev false positives |
|---|---|---|---|---|---|---|---|---|---|---|---|---|---|
| | | | | | | | | Specific gene | Sub-class | Class | Incorrect | | |
| 1 | 4 | VANZA | 30.783 | 14.248 | 0.628 | 0.290 | 0.920 | 0.760 | 0.160 | 0.000 | 0.080 | 0.440 | 1.261 |
| 2 | 52 | VANWG | 60.500 | 19.636 | 0.711 | 0.232 | 0.960 | 0.400 | 0.560 | 0.000 | 0.040 | 0.120 | 0.332 |
| 3 | 62 | CTX | 62.240 | 22.946 | 0.706 | 0.261 | 1.000 | 0.000 | 1.000 | 0.000 | 0.000 | 0.480 | 0.823 |
| 4 | 68 | MEFA | 88.600 | 25.492 | 0.723 | 0.209 | 1.000 | 0.360 | 0.640 | 0.000 | 0.000 | 0.280 | 0.542 |
| 5 | 76 | OXA | 50.880 | 20.767 | 0.612 | 0.250 | 1.000 | 0.000 | 1.000 | 0.000 | 0.000 | 0.360 | 0.700 |
| 6 | 92 | CATB | 30.960 | 18.571 | 0.481 | 0.290 | 1.000 | 0.280 | 0.680 | 0.040 | 0.000 | 0.240 | 0.831 |
| 7 | 162 | EREA | 56.364 | 38.857 | 0.457 | 0.315 | 0.880 | 0.320 | 0.560 | 0.000 | 0.120 | 0.160 | 0.624 |
| 8 | 174 | TEM | 55.958 | 15.267 | 0.663 | 0.183 | 0.960 | 0.040 | 0.920 | 0.000 | 0.040 | 0.200 | 0.645 |
| 9 | 193 | CML | 83.800 | 23.836 | 0.633 | 0.181 | 1.000 | 0.280 | 0.680 | 0.040 | 0.000 | 0.080 | 0.400 |
| 10 | 207 | NIMA | 23.917 | 12.014 | 0.513 | 0.263 | 0.960 | 0.760 | 0.160 | 0.040 | 0.040 | 0.480 | 1.686 |
| 11 | 239 | DHFR | 28.360 | 11.018 | 0.587 | 0.230 | 1.000 | 0.400 | 0.600 | 0.000 | 0.000 | 0.000 | 0.000 |
| 12 | 246 | FOLP | 89.720 | 24.630 | 0.749 | 0.206 | 1.000 | 0.960 | 0.040 | 0.000 | 0.000 | 0.080 | 0.400 |
| 13 | 274 | PARC | 207.278 | 32.207 | 0.930 | 0.146 | 0.720 | 0.000 | 0.560 | 0.160 | 0.280 | 2.240 | 2.368 |
| 14 | 276 | SHV | 85.167 | 7.597 | 0.972 | 0.085 | 0.960 | 0.000 | 0.960 | 0.000 | 0.040 | 2.080 | 2.900 |
| 15 | 284 | DFRA | 41.667 | 12.940 | 0.741 | 0.234 | 0.960 | 0.680 | 0.280 | 0.000 | 0.040 | 0.640 | 1.381 |
| 16 | 295 | SOXS | 26.560 | 7.119 | 0.802 | 0.219 | 1.000 | 0.640 | 0.360 | 0.000 | 0.000 | 1.000 | 2.141 |
| 17 | 321 | TSNR | 62.000 | 16.321 | 0.783 | 0.208 | 0.880 | 0.560 | 0.320 | 0.000 | 0.120 | 0.600 | 0.866 |
| 18 | 413 | OMPD | 63.520 | 16.259 | 0.586 | 0.151 | 1.000 | 1.000 | 0.000 | 0.000 | 0.000 | 0.000 | 0.000 |
| 19 | 486 | LMRA | 27.000 | 12.312 | 0.466 | 0.212 | 1.000 | 0.960 | 0.040 | 0.000 | 0.000 | 0.040 | 0.200 |
| 20 | 505 | CARB | 68.583 | 19.440 | 0.758 | 0.214 | 0.960 | 0.680 | 0.280 | 0.000 | 0.040 | 0.520 | 1.046 |
| 21 | 506 | ANT3-DPRIME | 51.240 | 13.252 | 0.637 | 0.167 | 1.000 | 0.920 | 0.080 | 0.000 | 0.000 | 0.120 | 0.332 |
| 22 | 555 | QNRS | 45.960 | 9.334 | 0.693 | 0.143 | 1.000 | 0.560 | 0.440 | 0.000 | 0.000 | 0.040 | 0.200 |
| 23 | 558 | VANWI | 55.318 | 20.051 | 0.487 | 0.177 | 0.880 | 0.840 | 0.040 | 0.000 | 0.120 | 0.080 | 0.400 |
| 24 | 578 | TETX | 100.960 | 17.714 | 0.862 | 0.153 | 1.000 | 0.360 | 0.640 | 0.000 | 0.000 | 2.080 | 4.271 |
| 25 | 652 | VGA | 111.520 | 32.218 | 0.705 | 0.204 | 1.000 | 0.880 | 0.120 | 0.000 | 0.000 | 0.920 | 2.344 |
| 26 | 682 | MOX | 111.160 | 10.850 | 0.960 | 0.093 | 1.000 | 0.160 | 0.800 | 0.040 | 0.000 | 1.680 | 1.952 |
| 27 | 694 | ACT | 106.720 | 16.794 | 0.851 | 0.134 | 1.000 | 0.120 | 0.880 | 0.000 | 0.000 | 0.440 | 1.003 |
| 28 | 717 | VANRE | 63.542 | 6.547 | 0.907 | 0.095 | 0.960 | 0.640 | 0.320 | 0.000 | 0.040 | 0.640 | 1.150 |
| 29 | 749 | RMTB | 16.810 | 7.420 | 0.640 | 0.287 | 0.840 | 0.640 | 0.200 | 0.000 | 0.160 | 0.800 | 2.021 |
| 30 | 778 | DHA | 62.400 | 29.537 | 0.542 | 0.259 | 1.000 | 0.000 | 1.000 | 0.000 | 0.000 | 0.320 | 0.748 |
| 31 | 789 | CMY | 63.080 | 27.296 | 0.548 | 0.237 | 1.000 | 0.000 | 1.000 | 0.000 | 0.000 | 0.160 | 0.624 |
| 32 | 797 | FACT | 150.000 | 20.613 | 0.887 | 0.123 | 0.760 | 0.480 | 0.280 | 0.000 | 0.240 | 0.400 | 0.764 |
| 33 | 819 | OKP | 84.833 | 4.833 | 0.975 | 0.056 | 0.960 | 0.000 | 0.960 | 0.000 | 0.040 | 2.440 | 2.293 |
| 34 | 973 | IMP | 48.520 | 21.804 | 0.645 | 0.291 | 1.000 | 0.000 | 1.000 | 0.000 | 0.000 | 0.400 | 1.080 |
| 35 | 1048 | VIM | 45.080 | 15.756 | 0.553 | 0.195 | 1.000 | 0.040 | 0.960 | 0.000 | 0.000 | 0.080 | 0.277 |
| 36 | 1135 | PBP1B | 96.444 | 35.196 | 0.389 | 0.142 | 0.720 | 0.720 | 0.000 | 0.000 | 0.280 | 0.080 | 0.400 |
| 37 | 1146 | MPHE | 71.840 | 9.419 | 0.806 | 0.106 | 1.000 | 0.520 | 0.480 | 0.000 | 0.000 | 0.280 | 0.843 |
| 38 | 1182 | VATE | 45.409 | 10.671 | 0.696 | 0.165 | 0.880 | 0.520 | 0.360 | 0.000 | 0.120 | 0.280 | 0.678 |
| 39 | 1214 | OPRJ | 136.227 | 13.596 | 0.940 | 0.094 | 0.880 | 0.360 | 0.520 | 0.000 | 0.120 | 0.600 | 0.645 |
| 40 | 1254 | FOSB | 38.560 | 5.229 | 0.897 | 0.125 | 1.000 | 0.560 | 0.440 | 0.000 | 0.000 | 1.960 | 3.434 |
| 41 | 1271 | VPH | 56.727 | 25.317 | 0.651 | 0.291 | 0.880 | 0.760 | 0.120 | 0.000 | 0.120 | 0.600 | 1.041 |
| 42 | 1283 | SULI | 71.083 | 12.991 | 0.836 | 0.154 | 0.960 | 0.160 | 0.760 | 0.040 | 0.040 | 0.560 | 1.044 |
| 43 | 1297 | TET40 | 41.043 | 21.582 | 0.332 | 0.174 | 0.920 | 0.480 | 0.440 | 0.000 | 0.080 | 0.040 | 0.200 |
| 44 | 1389 | CPXAR | 45.542 | 13.825 | 0.643 | 0.199 | 0.960 | 0.240 | 0.720 | 0.000 | 0.040 | 0.440 | 0.712 |
| 45 | 1392 | AAC6-PRIME | 14.760 | 7.423 | 0.261 | 0.132 | 1.000 | 0.800 | 0.200 | 0.000 | 0.000 | 0.000 | 0.000 |
| 46 | 1422 | VGBB | 78.600 | 11.281 | 0.875 | 0.130 | 1.000 | 0.680 | 0.320 | 0.000 | 0.000 | 1.400 | 3.109 |
| 47 | 1440 | FOSC | 28.960 | 12.431 | 0.514 | 0.223 | 1.000 | 0.920 | 0.080 | 0.000 | 0.000 | 0.120 | 0.440 |

| | | | | | | | | | | | | | |
|---|---|---|---|---|---|---|---|---|---|---|---|---|---|
| 48 | 1535 | LNUA | 25.760 | 12.367 | 0.525 | 0.252 | 1.000 | 0.400 | 0.600 | 0.000 | 0.000 | 0.160 | 0.473 |
| 49 | 1569 | PARE | 164.167 | 32.093 | 0.868 | 0.171 | 0.480 | 0.200 | 0.240 | 0.040 | 0.520 | 1.200 | 2.517 |
| 50 | 1695 | NDM | 59.292 | 13.687 | 0.721 | 0.168 | 0.960 | 0.840 | 0.080 | 0.040 | 0.040 | 0.240 | 0.663 |
| 51 | 1702 | SPG | 76.458 | 14.741 | 0.885 | 0.172 | 0.960 | 0.440 | 0.480 | 0.040 | 0.040 | 0.920 | 0.997 |
| 52 | 1753 | VEB | 77.680 | 19.491 | 0.855 | 0.214 | 1.000 | 0.040 | 0.960 | 0.000 | 0.000 | 0.360 | 0.757 |
| 53 | 1953 | LNUB | 68.042 | 12.267 | 0.839 | 0.152 | 0.960 | 0.720 | 0.240 | 0.000 | 0.040 | 0.160 | 0.374 |
| 54 | 2026 | ERMA | 54.292 | 18.155 | 0.732 | 0.246 | 0.960 | 0.720 | 0.240 | 0.000 | 0.040 | 0.160 | 0.374 |
| 55 | 2357 | SULII | 65.760 | 13.245 | 0.801 | 0.162 | 1.000 | 0.040 | 0.960 | 0.000 | 0.000 | 0.440 | 0.870 |
| 56 | 2517 | TET37 | 22.542 | 6.554 | 0.673 | 0.201 | 0.960 | 0.880 | 0.080 | 0.000 | 0.040 | 0.560 | 2.123 |
| 57 | 2822 | EMRK | 85.500 | 19.269 | 0.804 | 0.184 | 0.880 | 0.640 | 0.160 | 0.080 | 0.120 | 0.320 | 0.627 |
| 58 | 2999 | MPHB | 68.958 | 15.058 | 0.752 | 0.168 | 0.960 | 0.800 | 0.160 | 0.000 | 0.040 | 0.320 | 1.069 |
| 59 | 3024 | VANYM | 62.000 | 9.239 | 0.872 | 0.132 | 0.920 | 0.600 | 0.320 | 0.000 | 0.080 | 0.400 | 0.645 |
| 60 | 3041 | MECC | 189.160 | 19.796 | 0.939 | 0.098 | 1.000 | 0.280 | 0.680 | 0.040 | 0.000 | 0.040 | 0.200 |
| 61 | 3128 | TUFAB | 71.040 | 19.711 | 0.595 | 0.166 | 1.000 | 0.760 | 0.240 | 0.000 | 0.000 | 0.280 | 0.542 |
| 62 | 3176 | AMRB | 305.440 | 22.387 | 0.971 | 0.072 | 1.000 | 0.160 | 0.800 | 0.040 | 0.000 | 0.000 | 0.000 |
| 63 | 3270 | IRI | 140.957 | 8.450 | 0.972 | 0.059 | 0.920 | 0.240 | 0.680 | 0.000 | 0.080 | 1.440 | 1.660 |
| 64 | 3314 | RPOB | 356.000 | 0.000 | 1.000 | 0.000 | 0.200 | 0.000 | 0.200 | 0.000 | 0.800 | 1.320 | 2.750 |
| 65 | 3332 | TET35 | 62.208 | 17.093 | 0.553 | 0.153 | 0.960 | 0.960 | 0.000 | 0.000 | 0.040 | 0.000 | 0.000 |
| 66 | 3370 | CFRA | 86.542 | 16.519 | 0.817 | 0.157 | 0.960 | 0.840 | 0.120 | 0.000 | 0.040 | 0.200 | 0.645 |
| 67 | 3513 | BRP | 20.364 | 9.796 | 0.494 | 0.238 | 0.880 | 0.880 | 0.000 | 0.000 | 0.120 | 0.080 | 0.400 |
| 68 | 3613 | APH3-PRIME | 65.583 | 8.075 | 0.808 | 0.101 | 0.960 | 0.880 | 0.080 | 0.000 | 0.040 | 0.440 | 1.446 |
| 69 | 3697 | TETM | 184.880 | 15.584 | 0.959 | 0.082 | 1.000 | 0.120 | 0.880 | 0.000 | 0.000 | 1.760 | 2.087 |
| 70 | 3778 | IND | 60.880 | 11.791 | 0.821 | 0.161 | 1.000 | 0.040 | 0.960 | 0.000 | 0.000 | 0.480 | 0.963 |

Simulation settings:

k-mers: ‘constant’, k=10
Gene coverage: 1
Number of genes: 1
Errors: ‘on’, 0.05
Penalty score: 0.1
Thresholding: multiplier=2, all factors (1-on)
Entropy screening: ‘rand’

**Table S10.** Resistance genes simulations with 0.10 (20%) error rate

| No. | Database No. | Gene sub-class | Avg block for identification | StDev block for identification | Avg coverage for identification | StDev coverage for identification | Accuracy | Identification level (fraction of trials) | | | | Avg false positives | StDev false positives |
|---|---|---|---|---|---|---|---|---|---|---|---|---|---|
| | | | | | | | | Specific gene | Sub-class | Class | Incorrect | | |
| 1 | 4 | VANZA | 33.958 | 16.992 | 0.692 | 0.347 | 0.960 | 0.640 | 0.320 | 0.000 | 0.040 | 3.600 | 7.539 |
| 2 | 52 | VANWG | 76.136 | 11.503 | 0.895 | 0.136 | 0.880 | 0.160 | 0.720 | 0.000 | 0.120 | 4.000 | 7.767 |
| 3 | 62 | CTX | 78.160 | 13.530 | 0.888 | 0.154 | 1.000 | 0.000 | 1.000 | 0.000 | 0.000 | 0.760 | 0.970 |
| 4 | 68 | MEFA | 103.600 | 21.747 | 0.845 | 0.178 | 1.000 | 0.200 | 0.800 | 0.000 | 0.000 | 1.960 | 2.894 |
| 5 | 76 | OXA | 65.208 | 23.052 | 0.785 | 0.279 | 0.960 | 0.000 | 0.960 | 0.000 | 0.040 | 2.360 | 3.988 |
| 6 | 92 | CATB | 41.600 | 20.516 | 0.647 | 0.322 | 1.000 | 0.040 | 0.960 | 0.000 | 0.000 | 0.840 | 1.650 |
| 7 | 162 | EREA | 82.438 | 39.744 | 0.670 | 0.324 | 0.640 | 0.280 | 0.360 | 0.000 | 0.360 | 0.920 | 1.778 |
| 8 | 174 | TEM | 67.080 | 20.866 | 0.797 | 0.250 | 1.000 | 0.040 | 0.960 | 0.000 | 0.000 | 1.840 | 2.285 |
| 9 | 193 | CML | 100.520 | 30.206 | 0.761 | 0.229 | 1.000 | 0.040 | 0.880 | 0.080 | 0.000 | 0.200 | 0.408 |
| 10 | 207 | NIMA | 28.400 | 14.483 | 0.614 | 0.316 | 1.000 | 0.640 | 0.280 | 0.080 | 0.000 | 2.120 | 5.876 |
| 11 | 239 | DHFR | 34.320 | 10.135 | 0.712 | 0.213 | 1.000 | 0.120 | 0.880 | 0.000 | 0.000 | 0.920 | 1.730 |
| 12 | 246 | FOLP | 103.750 | 16.308 | 0.868 | 0.137 | 0.960 | 0.600 | 0.360 | 0.000 | 0.040 | 1.320 | 2.376 |
| 13 | 274 | PARC | 209.100 | 28.026 | 0.938 | 0.126 | 0.800 | 0.000 | 0.400 | 0.400 | 0.200 | 2.040 | 2.169 |
| 14 | 276 | SHV | 82.042 | 19.356 | 0.937 | 0.221 | 0.960 | 0.000 | 0.800 | 0.160 | 0.040 | 4.440 | 3.630 |
| 15 | 284 | DFRA | 53.478 | 6.755 | 0.954 | 0.122 | 0.920 | 0.160 | 0.720 | 0.040 | 0.080 | 4.920 | 4.573 |
| 16 | 295 | SOXS | 29.652 | 6.860 | 0.897 | 0.211 | 0.920 | 0.320 | 0.600 | 0.000 | 0.080 | 4.800 | 6.331 |
| 17 | 321 | TSNR | 73.870 | 9.172 | 0.935 | 0.117 | 0.920 | 0.320 | 0.600 | 0.000 | 0.080 | 2.080 | 2.914 |
| 18 | 413 | OMPD | 77.227 | 18.662 | 0.713 | 0.174 | 0.880 | 0.680 | 0.200 | 0.000 | 0.120 | 1.120 | 2.369 |
| 19 | 486 | LMRA | 37.696 | 10.877 | 0.652 | 0.194 | 0.920 | 0.800 | 0.120 | 0.000 | 0.080 | 0.440 | 1.294 |
| 20 | 505 | CARB | 78.875 | 16.894 | 0.870 | 0.189 | 0.960 | 0.360 | 0.520 | 0.080 | 0.040 | 1.680 | 2.495 |
| 21 | 506 | ANT3-DPRIME | 65.333 | 13.321 | 0.815 | 0.168 | 0.960 | 0.520 | 0.440 | 0.000 | 0.040 | 0.840 | 1.248 |
| 22 | 555 | QNRS | 56.833 | 10.945 | 0.857 | 0.166 | 0.960 | 0.240 | 0.680 | 0.040 | 0.040 | 1.360 | 2.343 |
| 23 | 558 | VANWI | 73.833 | 24.815 | 0.651 | 0.221 | 0.960 | 0.760 | 0.200 | 0.000 | 0.040 | 0.640 | 1.150 |
| 24 | 578 | TETX | 108.130 | 19.398 | 0.922 | 0.166 | 0.920 | 0.120 | 0.800 | 0.000 | 0.080 | 4.280 | 4.686 |
| 25 | 652 | VGA | 124.520 | 36.860 | 0.788 | 0.234 | 1.000 | 0.640 | 0.360 | 0.000 | 0.000 | 2.240 | 2.697 |
| 26 | 682 | MOX | 112.320 | 7.941 | 0.971 | 0.070 | 1.000 | 0.040 | 0.880 | 0.080 | 0.000 | 2.040 | 2.189 |
| 27 | 694 | ACT | 114.600 | 16.304 | 0.914 | 0.131 | 1.000 | 0.080 | 0.920 | 0.000 | 0.000 | 1.400 | 1.500 |
| 28 | 717 | VANRE | 66.280 | 6.374 | 0.946 | 0.092 | 1.000 | 0.400 | 0.600 | 0.000 | 0.000 | 2.400 | 3.379 |
| 29 | 749 | RMTB | 20.320 | 7.915 | 0.778 | 0.308 | 1.000 | 0.520 | 0.480 | 0.000 | 0.000 | 3.000 | 4.397 |
| 30 | 778 | DHA | 88.160 | 34.827 | 0.767 | 0.304 | 1.000 | 0.000 | 0.920 | 0.080 | 0.000 | 1.760 | 3.072 |
| 31 | 789 | CMY | 73.920 | 30.791 | 0.642 | 0.268 | 1.000 | 0.000 | 0.840 | 0.160 | 0.000 | 0.840 | 1.344 |
| 32 | 797 | FACT | 141.263 | 36.689 | 0.835 | 0.217 | 0.760 | 0.440 | 0.320 | 0.000 | 0.240 | 0.680 | 0.988 |
| 33 | 819 | OKP | 84.458 | 7.052 | 0.970 | 0.082 | 0.960 | 0.000 | 0.840 | 0.120 | 0.040 | 4.360 | 4.009 |
| 34 | 973 | IMP | 43.958 | 23.704 | 0.584 | 0.317 | 0.960 | 0.120 | 0.840 | 0.000 | 0.040 | 0.600 | 1.323 |
| 35 | 1048 | VIM | 60.240 | 15.613 | 0.741 | 0.195 | 1.000 | 0.000 | 1.000 | 0.000 | 0.000 | 0.600 | 1.443 |
| 36 | 1135 | PBP1B | 157.091 | 60.552 | 0.634 | 0.245 | 0.440 | 0.360 | 0.080 | 0.000 | 0.560 | 0.240 | 1.012 |
| 37 | 1146 | MPHE | 82.917 | 9.036 | 0.931 | 0.102 | 0.960 | 0.120 | 0.840 | 0.000 | 0.040 | 4.960 | 8.152 |
| 38 | 1182 | VATE | 52.500 | 15.291 | 0.807 | 0.236 | 0.960 | 0.400 | 0.560 | 0.000 | 0.040 | 1.080 | 1.801 |
| 39 | 1214 | OPRJ | 144.773 | 0.429 | 1.000 | 0.000 | 0.880 | 0.000 | 0.880 | 0.000 | 0.120 | 4.160 | 4.160 |
| 40 | 1254 | FOSB | 40.680 | 5.289 | 0.946 | 0.126 | 1.000 | 0.200 | 0.800 | 0.000 | 0.000 | 3.720 | 5.512 |
| 41 | 1271 | VPH | 63.542 | 28.278 | 0.730 | 0.325 | 0.960 | 0.560 | 0.400 | 0.000 | 0.040 | 1.720 | 2.424 |
| 42 | 1283 | SULI | 80.000 | 7.751 | 0.943 | 0.093 | 1.000 | 0.120 | 0.880 | 0.000 | 0.000 | 1.200 | 1.258 |
| 43 | 1297 | TET40 | 53.875 | 29.858 | 0.436 | 0.243 | 0.960 | 0.440 | 0.520 | 0.000 | 0.040 | 0.440 | 1.044 |
| 44 | 1389 | CPXAR | 58.190 | 13.144 | 0.821 | 0.189 | 0.840 | 0.240 | 0.520 | 0.080 | 0.160 | 0.960 | 1.695 |
| 45 | 1392 | AAC6-PRIME | 25.600 | 17.448 | 0.455 | 0.313 | 1.000 | 0.560 | 0.440 | 0.000 | 0.000 | 0.240 | 0.723 |
| 46 | 1422 | VGBB | 83.870 | 9.739 | 0.935 | 0.110 | 0.920 | 0.320 | 0.600 | 0.000 | 0.080 | 6.880 | 8.192 |
| 47 | 1440 | FOSC | 39.792 | 13.325 | 0.709 | 0.239 | 0.960 | 0.720 | 0.240 | 0.000 | 0.040 | 1.000 | 1.756 |

| | | | | | | | | | | | | | |
|---|---|---|---|---|---|---|---|---|---|---|---|---|---|
| 48 | 1535 | LNUA | 32.160 | 16.790 | 0.655 | 0.343 | 1.000 | 0.240 | 0.760 | 0.000 | 0.000 | 2.800 | 4.573 |
| 49 | 1569 | PARE | 183.750 | 14.849 | 0.972 | 0.079 | 0.320 | 0.040 | 0.240 | 0.040 | 0.680 | 1.400 | 2.432 |
| 50 | 1695 | NDM | 68.261 | 20.100 | 0.832 | 0.246 | 0.920 | 0.400 | 0.440 | 0.080 | 0.080 | 2.280 | 3.565 |
| 51 | 1702 | SPG | 80.720 | 11.059 | 0.935 | 0.129 | 1.000 | 0.320 | 0.680 | 0.000 | 0.000 | 2.600 | 2.858 |
| 52 | 1753 | VEB | 79.720 | 14.458 | 0.877 | 0.161 | 1.000 | 0.000 | 1.000 | 0.000 | 0.000 | 1.520 | 3.466 |
| 53 | 1953 | LNUB | 78.400 | 5.909 | 0.968 | 0.073 | 1.000 | 0.200 | 0.760 | 0.040 | 0.000 | 3.440 | 5.394 |
| 54 | 2026 | ERMA | 63.200 | 16.345 | 0.853 | 0.222 | 1.000 | 0.440 | 0.520 | 0.040 | 0.000 | 0.960 | 1.513 |
| 55 | 2357 | SULII | 70.958 | 12.757 | 0.865 | 0.156 | 0.960 | 0.000 | 0.920 | 0.040 | 0.040 | 1.720 | 2.558 |
| 56 | 2517 | TET37 | 27.609 | 6.073 | 0.824 | 0.190 | 0.920 | 0.520 | 0.400 | 0.000 | 0.080 | 4.880 | 10.902 |
| 57 | 2822 | EMRK | 86.348 | 18.458 | 0.813 | 0.174 | 0.920 | 0.560 | 0.280 | 0.080 | 0.080 | 0.800 | 1.581 |
| 58 | 2999 | MPHB | 74.609 | 13.550 | 0.813 | 0.152 | 0.920 | 0.640 | 0.280 | 0.000 | 0.080 | 0.760 | 1.234 |
| 59 | 3024 | VANYM | 64.391 | 9.380 | 0.906 | 0.134 | 0.920 | 0.360 | 0.560 | 0.000 | 0.080 | 6.480 | 8.510 |
| 60 | 3041 | MECC | 193.640 | 16.520 | 0.961 | 0.082 | 1.000 | 0.240 | 0.760 | 0.000 | 0.000 | 0.240 | 0.523 |
| 61 | 3128 | TUFAB | 90.292 | 23.447 | 0.758 | 0.198 | 0.960 | 0.320 | 0.640 | 0.000 | 0.040 | 0.800 | 1.258 |
| 62 | 3176 | AMRB | 294.640 | 70.201 | 0.937 | 0.223 | 1.000 | 0.000 | 0.920 | 0.080 | 0.000 | 0.000 | 0.000 |
| 63 | 3270 | IRI | 142.682 | 7.852 | 0.984 | 0.054 | 0.880 | 0.080 | 0.800 | 0.000 | 0.120 | 4.200 | 3.640 |
| 64 | 3314 | RPOB | 293.000 | 55.648 | 0.823 | 0.157 | 0.160 | 0.000 | 0.160 | 0.000 | 0.840 | 0.320 | 1.600 |
| 65 | 3332 | TET35 | 74.600 | 29.305 | 0.665 | 0.262 | 0.800 | 0.560 | 0.240 | 0.000 | 0.200 | 0.480 | 0.963 |
| 66 | 3370 | CFRA | 95.227 | 16.115 | 0.900 | 0.154 | 0.880 | 0.360 | 0.520 | 0.000 | 0.120 | 4.640 | 6.800 |
| 67 | 3513 | BRP | 29.286 | 9.023 | 0.718 | 0.222 | 0.840 | 0.640 | 0.200 | 0.000 | 0.160 | 2.080 | 7.405 |
| 68 | 3613 | APH3-PRIME | 72.792 | 10.266 | 0.898 | 0.128 | 0.960 | 0.520 | 0.440 | 0.000 | 0.040 | 2.720 | 4.440 |
| 69 | 3697 | TETM | 183.920 | 18.907 | 0.954 | 0.099 | 1.000 | 0.120 | 0.840 | 0.040 | 0.000 | 3.400 | 2.533 |
| 70 | 3778 | IND | 64.160 | 16.178 | 0.866 | 0.219 | 1.000 | 0.000 | 1.000 | 0.000 | 0.000 | 2.760 | 3.407 |

Simulation settings:

k-mers: ‘constant’, k=10
Gene coverage: 1
Number of genes: 1
Errors: ‘on’, 0.10
Penalty score: 0.1
Thresholding: multiplier=2, all factors (1-on)
Entropy screening: ‘rand’

**Table S11.** Simulations with 2 resistance genes (gene combinations and results)

| Combo No. | Database No. gene 1 | Database No. gene 2 | Gene sub-class gene 1 | Gene sub-class gene 2 |
|---|---|---|---|---|
| 1 | 76 | 505 | OXA | CARB |
| 2 | 92 | 3024 | CATB | VANYM |
| 3 | 1048 | 506 | VIM | ANT3-DPRIME |
| 4 | 1182 | 778 | VATE | DHA |
| 5 | 1702 | 3270 | SPG | IRI |
| 6 | 2357 | 694 | SULII | ACT |
| 7 | 2999 | 68 | MPHB | MEFA |
| 8 | 3041 | 1048 | MECC | VIM |
| 9 | 3128 | 284 | TUFAB | DFRA |
| 10 | 3370 | 3024 | CFRA | VANYM |

| Combo No. | Avg block for identification | StDev block for identification | Avg coverage for identification | StDev coverage for identification | Accuracy | Identification level (fraction of trials) | | | | Avg false positives | StDev false positives |
|---|---|---|---|---|---|---|---|---|---|---|---|
| | | | | | | Specific gene | Sub-class | Class | Incorrect | | |
| 1 | 91.042 | 47.288 | 0.534 | 0.265 | 0.960 | 0.100 | 0.760 | 0.100 | 0.040 | 1.000 | 2.000 |
| 2 | 64.188 | 31.773 | 0.481 | 0.255 | 0.960 | 0.300 | 0.640 | 0.020 | 0.040 | 0.280 | 1.021 |
| 3 | 81.959 | 39.022 | 0.511 | 0.232 | 0.980 | 0.060 | 0.900 | 0.020 | 0.020 | 0.880 | 1.900 |
| 4 | 96.060 | 38.859 | 0.536 | 0.212 | 1.000 | 0.140 | 0.840 | 0.020 | 0.000 | 1.240 | 1.715 |
| 5 | 191.283 | 32.959 | 0.821 | 0.147 | 0.920 | 0.640 | 0.260 | 0.020 | 0.080 | 1.080 | 0.812 |
| 6 | 120.040 | 41.134 | 0.597 | 0.177 | 1.000 | 0.060 | 0.940 | 0.000 | 0.000 | 0.000 | 0.000 |
| 7 | 128.771 | 43.605 | 0.605 | 0.205 | 0.960 | 0.280 | 0.680 | 0.000 | 0.040 | 0.040 | 0.200 |
| 8 | 228.604 | 59.409 | 0.831 | 0.199 | 0.960 | 0.440 | 0.060 | 0.460 | 0.040 | 0.600 | 0.957 |
| 9 | 85.634 | 26.347 | 0.489 | 0.162 | 0.820 | 0.700 | 0.120 | 0.000 | 0.180 | 0.120 | 0.440 |
| 10 | 131.500 | 46.095 | 0.754 | 0.256 | 0.880 | 0.580 | 0.300 | 0.000 | 0.120 | 0.440 | 1.193 |

Simulation settings:

k-mers: ‘constant’, k=10
Gene coverage: 1
Number of genes: 2
Errors: ‘off’
Penalty score: 0.1
Thresholding: multiplier=4, all factors (1-on)
Entropy screening: ‘rand’

**Table S12.** Simulations with 5 resistance genes (gene combinations and results)

| Combo No. | Database No. gene 1 | Database No. gene 2 | Database No. gene 3 | Database No. gene 4 | Database No. gene 5 | Gene sub-class gene 1 | Gene sub-class gene 2 | Gene sub-class gene 3 | Gene sub-class gene 4 | Gene sub-class gene 5 |
|---|---|---|---|---|---|---|---|---|---|---|
| 1 | 295 | 1182 | 819 | 555 | 239 | SOXS | VATE | OKP | QNRS | DHFR |
| 2 | 973 | 2026 | 3041 | 1753 | 694 | IMP | ERMA | MECC | VEB | ACT |
| 3 | 1048 | 3778 | 3270 | 789 | 2517 | VIM | IND | IRI | CMY | TET37 |
| 4 | 3370 | 276 | 1422 | 3778 | 1702 | CFRA | SHV | VGBB | IND | SPG |
| 5 | 3778 | 506 | 274 | 694 | 778 | IND | ANT3-DPRIME | PARC | ACT | DHA |

| Combo No. | Avg block for identification | StDev block for identification | Avg coverage for identification | StDev coverage for identification | Accuracy | Identification level (fraction of trials) | | | | Avg false positives | StDev false positives |
|---|---|---|---|---|---|---|---|---|---|---|---|
| | | | | | | Specific gene | Sub-class | Class | Incorrect | | |
| 1 | 158.889 | 91.030 | 0.524 | 0.298 | 0.792 | 0.048 | 0.656 | 0.088 | 0.208 | 4.520 | 4.224 |
| 2 | 441.777 | 186.481 | 0.793 | 0.324 | 0.824 | 0.032 | 0.720 | 0.072 | 0.176 | 9.560 | 5.738 |
| 3 | 245.739 | 130.588 | 0.550 | 0.286 | 0.736 | 0.008 | 0.640 | 0.088 | 0.264 | 3.800 | 3.862 |
| 4 | 281.263 | 138.899 | 0.644 | 0.312 | 0.912 | 0.104 | 0.640 | 0.168 | 0.088 | 11.640 | 7.059 |
| 5 | 330.667 | 168.319 | 0.549 | 0.266 | 0.672 | 0.000 | 0.640 | 0.032 | 0.328 | 4.760 | 5.372 |

Simulation settings:

k-mers: ‘constant’, k=10
Gene coverage: 1
Number of genes: 5
Errors: ‘off’
Penalty score: 0.1
Thresholding: multiplier=25, all factors (1-on)
Entropy screening: ‘rand’

**Table S13.** 10 randomly-selected cancer genes

| No. | Gene database No. | Sub-class | Full gene name (from COSMIC database) |
|---|---|---|---|
| 1 | 1049 | CMPK1 | CMPK1 ENST00000371873 1:47333946-47376745(+) |
| 2 | 2851 | C1orf115 | C1orf115 ENST00000294889 1:220690403-220696731(+) |
| 3 | 5025 | MTMR14 | MTMR14 ENST00000296003 3:9649584-9701973(+) |
| 4 | 7924 | CARTPT | CARTPT ENST00000296777 5:71719294-71720615(+) |
| 5 | 9305 | C6orf25 | C6orf25_ENST00000375806 ENST00000375806 6:31723384-31725074(+) |
| 6 | 15404 | FRG2B | FRG2B ENST00000425520 10:133625099-133626742(-) |
| 7 | 19240 | RBM23 | RBM23 ENST00000359890 14:22901730-22911393(-) |
| 8 | 21814 | PDXDC2 | PDXDC2 ENST00000331116 16:69996455-70065776(-) |
| 9 | 24929 | SLC7A10 | SLC7A10 ENST00000253188 19:33208891-33225703(-) |
| 10 | 27882 | CSF2RA | CSF2RA ENST00000381529 23:1282704-1309479(+) |

**Table S14.** Cancer genes simulations

| No. | Database No. | Gene sub-class | Avg block for identification | StDev block for identification | Avg coverage for identification | StDev coverage for identification | Accuracy | Identification level (fraction of trials) | | | | Avg false positives | StDev false positives |
|---|---|---|---|---|---|---|---|---|---|---|---|---|---|
| | | | | | | | | Specific gene | Sub-class | Class | Incorrect | | |
| 1 | 1049 | CMPK1 | 11.700 | 3.889 | 0.169 | 0.057 | 1.000 | 1.000 | 0.000 | 0.000 | 0.000 | 0.000 | 0.000 |
| 2 | 2851 | C1orf115 | 12.100 | 6.297 | 0.268 | 0.146 | 1.000 | 1.000 | 0.000 | 0.000 | 0.000 | 0.000 | 0.000 |
| 3 | 5025 | MTMR14 | 58.200 | 21.872 | 0.295 | 0.112 | 1.000 | 1.000 | 0.000 | 0.000 | 0.000 | 0.000 | 0.000 |
| 4 | 7924 | CARTPT | 19.600 | 5.758 | 0.536 | 0.160 | 1.000 | 1.000 | 0.000 | 0.000 | 0.000 | 0.000 | 0.000 |
| 5 | 9305 | C6orf25 | 34.200 | 10.326 | 0.467 | 0.142 | 1.000 | 0.900 | 0.100 | 0.000 | 0.000 | 0.000 | 0.000 |
| 6 | 15404 | FRG2B | 25.800 | 8.217 | 0.306 | 0.097 | 1.000 | 1.000 | 0.000 | 0.000 | 0.000 | 0.000 | 0.000 |
| 7 | 19240 | RBM23 | 50.900 | 13.110 | 0.379 | 0.097 | 1.000 | 1.000 | 0.000 | 0.000 | 0.000 | 0.000 | 0.000 |
| 8 | 21814 | PDXDC2 | 39.000 | 10.770 | 0.274 | 0.075 | 1.000 | 1.000 | 0.000 | 0.000 | 0.000 | 0.000 | 0.000 |
| 9 | 24929 | SLC7A10 | 51.300 | 22.081 | 0.324 | 0.139 | 1.000 | 1.000 | 0.000 | 0.000 | 0.000 | 0.000 | 0.000 |
| 10 | 27882 | CSF2RA | 45.900 | 17.451 | 0.377 | 0.143 | 1.000 | 0.400 | 0.600 | 0.000 | 0.000 | 0.000 | 0.000 |

Simulation settings:

k-mers: ‘constant’, k=10
Gene coverage: 1
Number of genes: 1
Errors: ‘off’
Penalty score: 0.1
Thresholding: multiplier=1, all factors (1-on)
Entropy screening: ‘rand’

**Table S15.** 10 randomly-selected genetic disease genes

| No. | Gene databse No. | Sub-class | Full gene name (from custom compiled database) |
|---|---|---|---|
| 1 | 28 | TBR1 | NG_046904.1 Homo sapiens T-box, brain 1 (TBR1), RefSeqGene on chromosome 2 |
| 2 | 109 | SHH | NG_007504.2 Homo sapiens sonic hedgehog (SHH), RefSeqGene on chromosome 7 |
| 3 | 110 | SIX3 | NG_016222.1 Homo sapiens SIX homeobox 3 (SIX3), RefSeqGene on chromosome 2 |
| 4 | 112 | ZIC2 | NG_007085.3 Homo sapiens Zic family member 2 (ZIC2), RefSeqGene on chromosome 13 |
| 5 | 121 | KRAS | NG_007524.1 Homo sapiens KRAS proto-oncogene, GTPase (KRAS), RefSeqGene on chromosome 12 |
| 6 | 143 | ALAD | NG_008716.1 Homo sapiens aminolevulinate dehydratase (ALAD), RefSeqGene on chromosome 9 |
| 7 | 163 | IGF2 | NG_008849.1 Homo sapiens insulin like growth factor 2 (IGF2), RefSeqGene on chromosome 11 |
| 8 | 202 | PDE6G | NG_009834.1 Homo sapiens phosphodiesterase 6G (PDE6G), RefSeqGene on chromosome 17 |
| 9 | 214 | ROM1 | NG_009845.1 Homo sapiens retinal outer segment membrane protein 1 (ROM1), RefSeqGene on chromosome 11 |
| 10 | 242 | UBA1 | NG_009161.1 Homo sapiens ubiquitin like modifier activating enzyme 1 (UBA1), RefSeqGene on chromosome X |

**Table S16**. Genetic disease genes simulations

| No. | Database No. | Gene sub-class | Avg block for identification | StDev block for identification | Avg coverage for identification | StDev coverage for identification | Accuracy | Identification level (fraction of trials) | | | | Avg false positives | StDev false positives |
|---|---|---|---|---|---|---|---|---|---|---|---|---|---|
| | | | | | | | | Specific gene | Sub-class | Class | Incorrect | | |
| 1 | 28 | TBR1 | 70.900 | 43.322 | 0.044 | 0.027 | 1.000 | 1.000 | 0.000 | 0.000 | 0.000 | 0.000 | 0.000 |
| 2 | 109 | SHH | 216.200 | 151.258 | 0.112 | 0.078 | 1.000 | 1.000 | 0.000 | 0.000 | 0.000 | 0.000 | 0.000 |
| 3 | 110 | SIX3 | 119.700 | 38.251 | 0.107 | 0.034 | 1.000 | 1.000 | 0.000 | 0.000 | 0.000 | 0.000 | 0.000 |
| 4 | 112 | ZIC2 | 28.900 | 25.562 | 0.024 | 0.021 | 1.000 | 1.000 | 0.000 | 0.000 | 0.000 | 0.000 | 0.000 |
| 5 | 121 | KRAS | 126.200 | 49.497 | 0.024 | 0.009 | 1.000 | 1.000 | 0.000 | 0.000 | 0.000 | 0.000 | 0.000 |
| 6 | 143 | ALAD | 327.200 | 159.264 | 0.148 | 0.072 | 1.000 | 1.000 | 0.000 | 0.000 | 0.000 | 0.000 | 0.000 |
| 7 | 163 | IGF2 | 186.900 | 161.765 | 0.068 | 0.059 | 1.000 | 1.000 | 0.000 | 0.000 | 0.000 | 0.000 | 0.000 |
| 8 | 202 | PDE6G | 140.500 | 104.140 | 0.107 | 0.079 | 1.000 | 1.000 | 0.000 | 0.000 | 0.000 | 0.000 | 0.000 |
| 9 | 214 | ROM1 | 185.200 | 156.556 | 0.197 | 0.167 | 1.000 | 1.000 | 0.000 | 0.000 | 0.000 | 0.000 | 0.000 |
| 10 | 242 | UBA1 | 1522.300 | 768.633 | 0.486 | 0.245 | 1.000 | 1.000 | 0.000 | 0.000 | 0.000 | 0.000 | 0.000 |

Simulation settings:

k-mers: ‘constant’, k=10
Gene coverage: 1
Number of genes: 1
Errors: ‘off’
Penalty score: 0.1
Thresholding: multiplier=3-5, all factors (1-on)
Entropy screening: ‘rand’
